# Single transcript-level metabolic responsive landscape of human liver transcriptome

**DOI:** 10.1101/2023.07.06.544842

**Authors:** Chengfei Jiang, Ping Li, Yonghe Ma, Nao Yoneda, Megumi Nishiwaki, Yasuyuki Ohnishi, Hiroshi Suemizu, Haiming Cao

**Affiliations:** Cardiovascular Branch National Heart, Lung and Blood Institute National Institutes of Health Bethesda, MD 20892, USA; Liver Engineering Laboratory Department of Applied Research for Laboratory Animals Central Institute for Experimental Animals (CIEA) 3-25-12 Tonomachi, Kawasaki-ku, Kawasaki 210-0821, Japan; Fujiyama Breeding Facility, CLEA Japan, Inc, 4839-23 Kitayama, Fujinomiya, Shizuoka 418-0122, Japan.

**Author notes:** To whom correspondence should be addressed: Haiming Cao, Ph.D. Cardiovascular Branch National Heart, Lung and Blood Institute National Institutes of Health Bethesda, MD 20892, USA.

## Abstract

Direct knowledge of gene regulation in human liver by metabolic stimuli could fundamentally advance our understanding of metabolic physiology. This information, however, is largely unknown, and this void is deeply rooted in a paradox that is caused by the inaccessibility of human liver to treatments and the insufficient annotation of liver transcriptome. Recent advances have uncovered immense complexity of transcriptome, i.e., multiple transcripts produced from one gene and extensive RNA modifications, which are all highly regulated and often condition-dependent. Establishing an inclusive annotation to study liver transcriptome dynamics thus requires human liver samples of diverse conditions, which are paradoxically unavailable due to the inaccessibility of human liver to treatments. In this work, we addressed these challenges by coupling an isogenic humanized mouse model with Nanopore single-molecule direct RNA sequencing (DRS). We first generated mice that carried humanized livers of identical genetic background, which were equivalent to clones of a single human liver, and then subjected the mice to representative metabolic treatments. We then analyzed the humanized livers with Nanopore DRS, which directly reads full-length native RNAs to determine the expression level, m6A modification and poly(A) tail length of all RNA transcript isoforms. Thus, our system allows for constructing a de novo annotation of human liver transcriptomes reflecting metabolic responses and studying transcriptome dynamics in conjunction. Our analysis uncovered a vast number of novel genes and transcripts that have not been previously reported. Our transcript-level analysis of human liver transcriptomes also identified a multitude of regulated metabolic pathways that were otherwise invisible using conventional short read RNA-seq. We also revealed for the first time the dynamic changes in m6A and poly(A) tail length of human liver transcripts many of which are transcribed from key metabolic genes. Furthermore, we performed comparative analyses of gene regulation between human and mouse and between two individuals using the liver-specific humanized mice. This revealed that transcriptome dynamics are highly species- and genetic background-dependent, which may only be faithfully studied in a humanized system that entails clones of the same human liver. Hence our work revealed a complex metabolic responsive landscape of human liver transcriptome and also provided a framework to understand transcriptome dynamics of human liver in response to physiologically relevant metabolic stimuli.

## Introduction

As the central metabolic organ, the liver can promptly adjust its transcriptome to respond to metabolic stimuli. Mounting evidence in animal studies support that deciphering the transcriptome dynamics is the key to unlock the molecular basis of liver metabolic physiology^1,2^. Direct information on gene regulation in human liver by metabolic stimuli, however, are largely unknown. As normal human livers are generally inaccessible to treatments, knowledge of stimulus-provoked changes in their transcriptomes heavily depend on the extrapolation from animal studies, particularly those done in mice. Growing evidence, however, has revealed substantial differences in gene regulation between human and mouse livers^3,4^. To directly study gene regulation in human liver, the only option is to analyze liver biopsy samples of patients and, very rarely, healthy controls. However, information from these studies is generally very limited and sometimes even misleading. The major hurdle limiting the power of patient-based studies is the diverse genetic backgrounds and environmental exposures of the patient population which often have substantial impact on gene regulation. It has been shown that in a general population, the vast majority of human genes (86.1%) are subject to the regulation of local genetic variants and exhibit differential expression among individuals^5^. While cohort statistics can be employed to identify the fraction of commonly regulated genes, the holistic picture of the metabolic responsive landscape of the transcriptome would have been lost. Ideally and hypothetically, such a study should be performed repeatedly on the same person or on multiple clones of the same individual, which is obviously impossible. Thus, direct knowledge of the liver transcriptome dynamics in humans is currently unknown and therefore hindering the overall understanding of human liver pathophysiology.

Aside from the limited accessibility to treatments discussed above, another critical obstacle to understanding gene regulation in human liver is the insufficient annotation of its transcriptome. The current mainstream approach to study global gene regulation is short-read RNA sequencing (RNA-seq), which requires an annotation to map the reads for gene identification. Recent advances have continued to unravel the complexity of the human transcriptome: a single gene can produce 10 or more transcript isoforms on average^6,7^ and RNA transcripts often go through extensive modifications, such as m6A and changes in length of the poly(A) tail, all of which impact RNA metabolism and have shown to play diverse roles in metabolic physiology^8–11^. Intriguingly, expression levels of transcript isoforms and RNA modifications are often highly regulated and condition-dependent. Although major annotations, such as GENCODE^12^ and RefSeq^13^, provide useful references for gene loci, they do not have a deep coverage of transcript isoforms and RNA modifications, especially those that specifically occur in liver under defined metabolic conditions. The reason for this insufficient annotation is partly caused by the limited accessibility of human livers to treatments precluding liver samples of diverse metabolic conditions to establish an inclusive annotation. This literally creates a paradox, i.e., reliable assessment of transcriptome dynamics in inaccessible human liver requires a comprehensive annotation of its transcriptome, which is contingent on the availability of liver samples of diverse conditions. The only approach to solve this paradox is to construct a de novo annotation of human liver transcriptomes under diverse metabolic conditions and simultaneously examine gene regulation at the levels of transcript isoforms and RNA modifications.

In this work, we produced mice that carry humanized livers of identical genetic background and subjected them to diverse metabolic stimuli, which essentially makes the human liver as accessible and genetically simple as an inbred mouse strain. We then performed Nanopore DRS on the humanized livers and uncovered a vast number of novel genes and transcript isoforms that were not previously reported. We also defined the regulation of human liver genes at transcript and RNA modification levels. Our work revealed substantial transcriptome dynamics that were not previously recognized, and this information could serve as a valuable resource to advance the understanding of human metabolic physiology.

## Results

### A de novo annotation of human liver transcriptomes reflecting pathophysiologically relevant metabolic responses

To enable a thorough examination of gene regulation in human liver by representative metabolic conditions, we first established an inclusive annotation of human liver transcriptomes reflecting these conditions. To make the human liver accessible to metabolic treatments, we employed a liver-specific humanized mouse model in which approximately 50% of the mouse hepatocytes were replaced by human ones^14–16^. All mice in the same experimental group were engrafted with hepatocytes from the same donor, a setting to mimic human clones and to reduce the impact of diverse genetic backgrounds. To capture transcriptomes reflecting pathophysiology and therapeutic development of metabolic diseases, we subjected the humanized mice to the following treatments: fasting (Fast) and ad libitum (AL), two ends of caloric cycle and known to regulate the expression of most metabolic genes^17^; activation of key transcriptional factors of metabolic pathways: PPARα, a Fast-induced master regulator of fatty acid oxidation^18^; PPARγ, an activator of lipogenic genes^19^; and FXR, a receptor for bile acids and a regulator of broad metabolic pathways that is being explored as a drug target for metabolic disorders^20,21^. In light of the increased complexity of RNA metabolism, we analyzed the transcriptome using Nanopore direct RNA sequencing (DRS). By dispensing with reverse transcription and cDNA amplification and directly reading native RNAs, Nanopore DRS has been shown to be able to discriminate similar transcript isoforms, detect novel genes and transcripts, and quantify differential gene expressions at single transcript-level with much greater fidelity and accuracy compared to conventional short-read RNA-Seq^22–24^

DRS analyses of humanized livers revealed that DRS was capable of detecting RNA transcripts of up to 15,000 bp long and most of them were in the range of 1,000 to 5,000 bp in both human and mouse (Fig. S1A). To verify the capability of DRS to ascertain differential gene expressions, we compared differentially expressed genes (DEGs) between Fast and AL detected by short-read RNA-seq and DRS, which showed similar expression patterns during fasting for both human and mouse genes (Fig. S1B). Furthermore, the differential expression genes (DEGs) generated from DRS were significantly enriched in fatty acid metabolism related pathways for human and mouse, which is similar to the results of short read RNA-seq (Fig. S1C). Moreover, GSVA results, indicating the biological function for each sample, also identified similar enriched patterns between DRS and short-read sequencing in Fast and AL group respectively (Fig. S1D). These data supported that Nanopore DRS is equally capable as short-read RNA-seq in capturing transcriptome dynamics at the gene level.

It is well known that most genes transcribe into multiple transcript isoforms but the full spectrum of the RNA transcripts in human livers, especially those that are conditionally expressed in response to metabolic stimuli, have not been carefully annotated. Here we constructed a de novo annotation of human and mouse transcriptomes based on the Nanopore DRS analysis of humanized livers. Our annotation resulted in a substantial expansion of RNA transcripts in both species: 54.6% of human and 70.8% of mouse transcripts isoforms are novel compared to those in reference annotations (GENCODE v33 and GENCODE vM24) (Fig. 1A and Table S1). Furthermore, over 30% of all transcripts in this novel annotation were only detectable under one or multiple specific treatment conditions (Fig. S1E), strongly arguing that gene regulation in human liver by metabolic treatments can only be adequately studied when an inclusive annotation that reflects these treatments is simultaneously established.

**Fig. 1.**
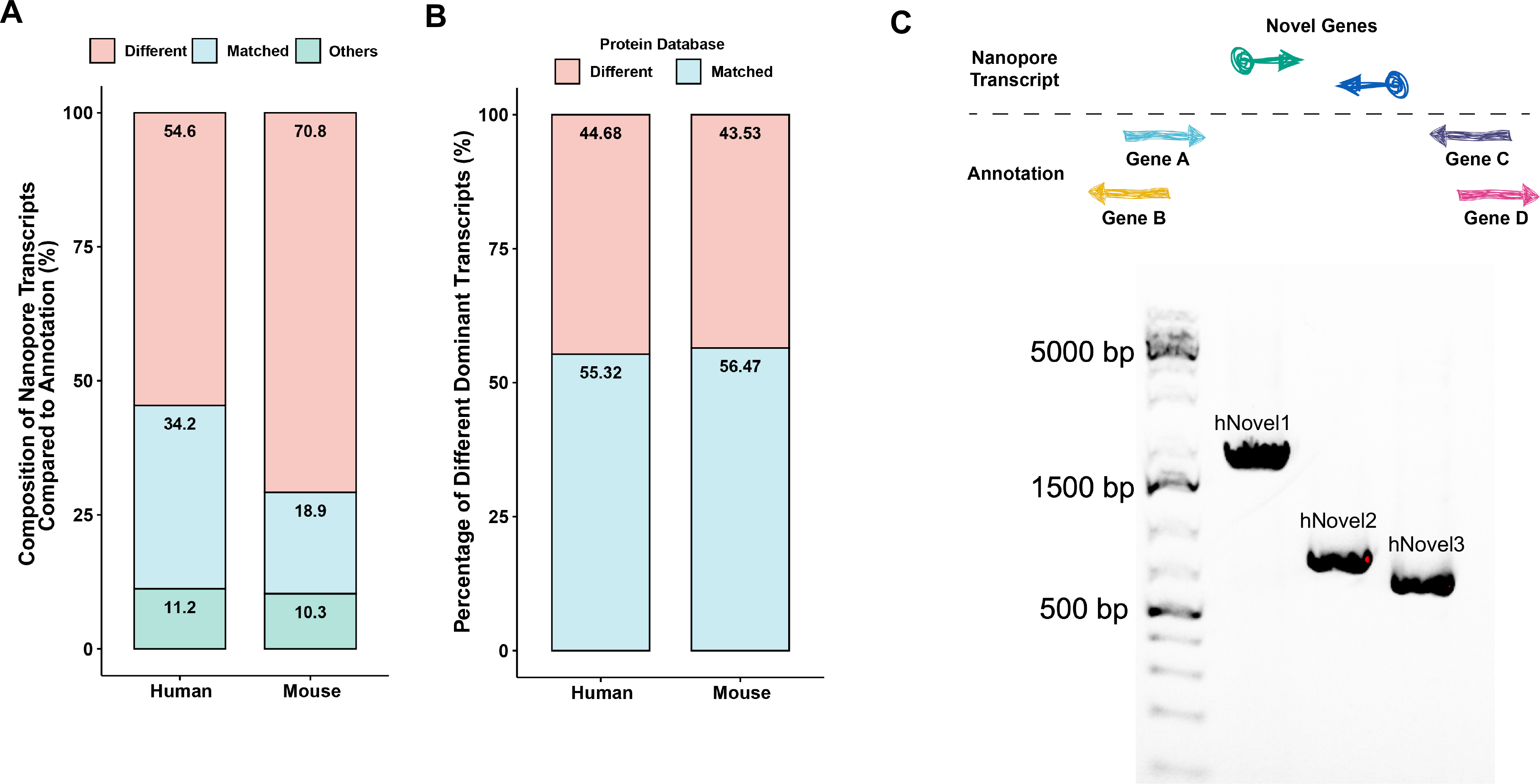
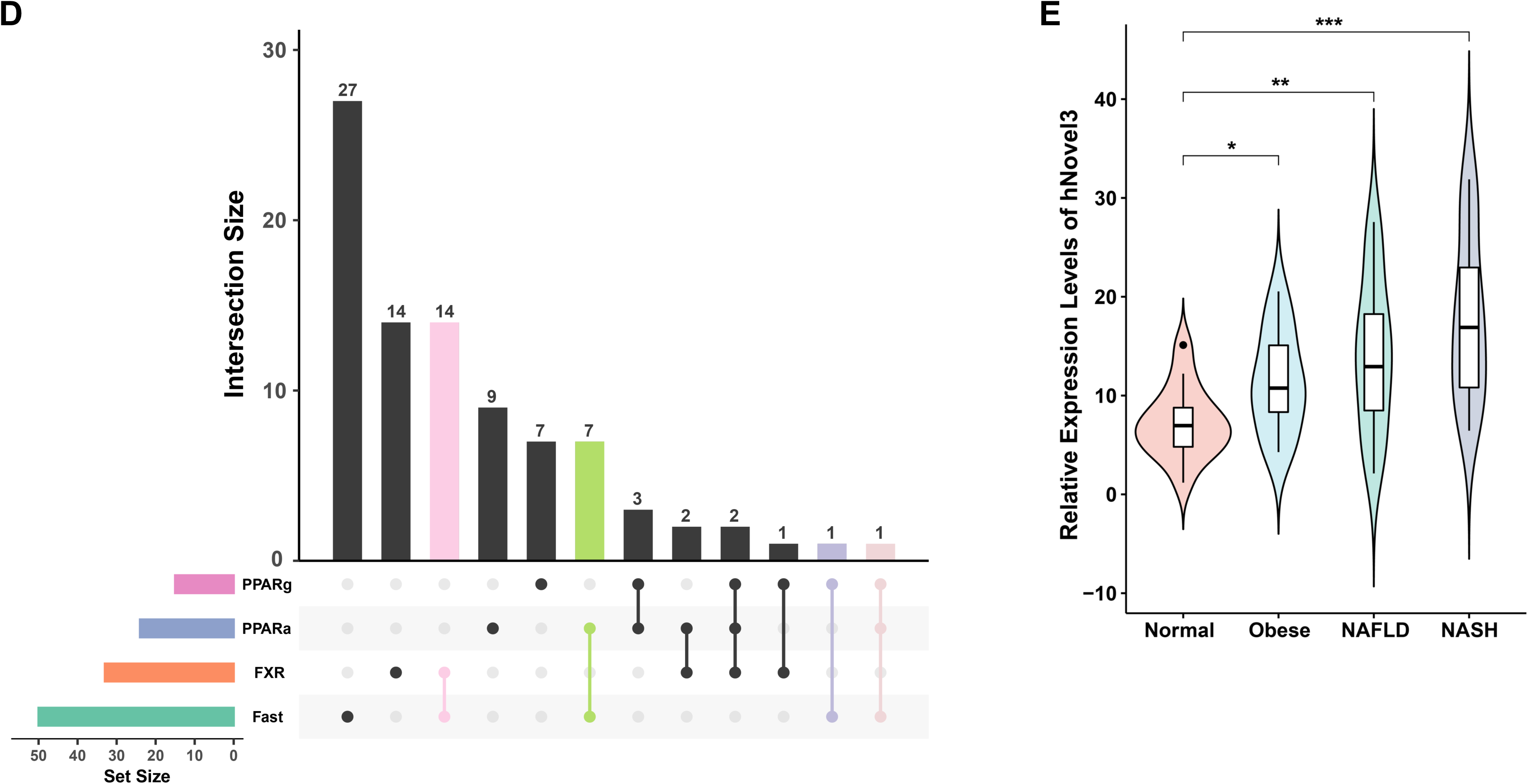
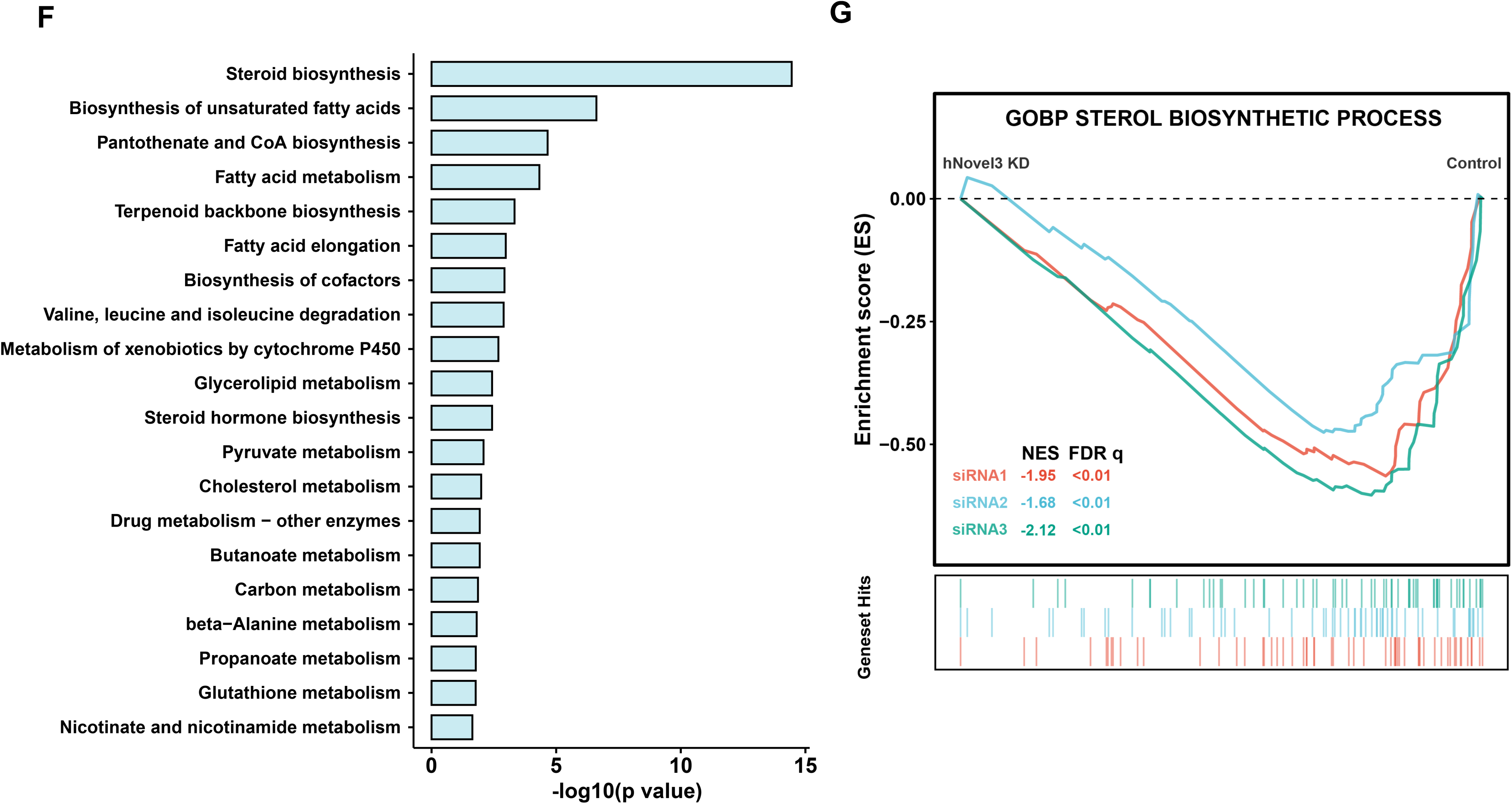
A de novo annotation of human liver transcriptomes reflecting pathophysiologically relevant metabolic responses. (A) Comparison of DRS and reference annotations (GENCODE v33 for human and GENCODE vM24 for mouse respectively). Transcripts that have been defined as matched transcripts (Matched) were those with completed and exact matched splicing sites to the ones in the reference annotation. The different transcripts (Different) were the ones that had been found to have different isoform structures compared to the genes collected in reference annotations. Other isoforms (Others) were those that had mismatched splicing sites or had no actual overlaps to entries in reference annotation. (B) Comparison of different/novel transcript protein sequences identified by DRS with those in reference databases (GENCODE v33 for human and GENCODE vM24 for mouse respectively). The dominantly expressed novel transcripts were sent to ORF predictions. The longest ORFs were chosen, and the matched ones (Matched) were characterized as having an E-value of less than 0.01 and alignment coverage of more than 90%. The remaining ones were considered different (Different). (C) Diagram schematic of the novel gene definition (Top). Gel analysis of the PCR cloned novel genes (Bottom). (D) UpSet plot displaying the overlaps of novel genes regulated by several metabolic interventions. Novel genes commonly regulated in Fast and other treatments were labeled by different colors. (E) Expression levels of human novel 3 (hNovel 3) in the livers of healthy (Normal), obese (Obese), nonalcoholic fatty liver (NAFL), and nonalcoholic steatohepatitis (NASH) patients (Non-alcoholic fatty liver disease database, PRJNA523510). Data represent mean ± SEM, * p<0.05, **p < 0.01, ***p < 0.001, two-tailed unpaired Student’s t-test. (F) The bar plot of the top enriched KEGG pathways correlated with hNovel3. Pearson’s correlation analysis was performed to predict the correlated genes of hNovel3 in humanized mice samples. After performing a pathway analysis on the top 250 hNovel3 correlated genes, the top 20 enriched pathways were shown. (G) Gene Set Enrichment Analysis (GSEA) enrichment plots for the Sterol Biosynthetic Process pathway using the transcriptome data of hNovel3 knockdown (siRNA1, siRNA2, and siRNA3) and control samples.

While a single gene often transcribes into multiple transcript isoforms, the functional importance of these isoforms varies to a great degree and the dominant transcripts usually have a greater chance to be translated to proteins and play an important functional role^25^. We evaluated the composition of dominant transcripts per gene for novel transcripts identified by DRS and found that 16.45% of the novel transcripts in human and 18.17% in mouse were the dominant ones for their represented genes (Fig. S1F). Moreover, we found that more than 40% of these novel dominant transcripts displayed protein coding sequences that are different from those in current reference databases (Fig. 1B) supporting the idea that these novel transcripts significantly contribute to protein diversities in human liver.

In addition to protein-coding genes, noncoding genes have also been shown to play an increasingly prominent role in liver metabolism^14,15,26,27^. To characterize novel transcripts located in noncoding regions, we analyzed the coding potential for these transcripts using five algorithms (CPC2, PLEK, CPAT, CNCI, FEELnc) and found that they were mostly noncoding transcripts. A small fraction of them (5.5%), however, indeed have coding potential (Fig. S1G). For example, we detected a novel transcript in the locus of noncoding gene AC005538.3, whose structure was different from that in the reference annotation (Fig. S1H). Short-read RNA-seq also detected several reads in the exons of this novel transcript but the read peaks were ambiguous compared to DRS ones. Interestingly, while the host gene was originally labelled as noncoding in the reference annotation, this novel transcript displayed very high coding potential suggesting our novel annotation could be used to identify novel functional elements in a genomic locus. It should be noted, vice versa, that 3% novel transcripts derived from the coding genes contain no clear coding sequence (CDS), thus potentially functioning as noncoding RNAs. Thus, these novel transcripts revealed by our de novo annotation could serve as an important resource to further characterize the function of coding and noncoding transcripts in the context of liver metabolism and metabolic disorders.

Furthermore, we were able to identify 212 novel genes in intergenic regions where no gene has been recorded in current reference annotations, again pointing to the insufficient annotations of human liver transcriptome (Table S2). To confirm the expression of these novel genes, we randomly cloned some of the novel genes from cDNA of humanized livers and confirmed that their sequences matched those revealed by Nanopore DRS (Fig. 1C and Table S3). To further explore the expression of these novel genes in human populations and their implications in liver disease, we reanalyzed a published liver RNA-seq dataset from a nonalcoholic fatty liver disease (NAFLD) study^28^ using our novel annotation, and readily detected 160 novel genes in at least one third of the livers in this NAFLD cohort. Intriguingly, the expression levels of 38.8% of these novel genes were significantly changed in patients with NAFLD (Fig. S1I), suggesting that dysregulation of these novel genes could potentially contribute to the pathogenesis of NAFLD.

Moreover, more than 40% of the novel genes were responsive to at least one metabolic stimulus (fasting or activation of key metabolic regulators) and some of them could be co-regulated by multiple treatments (Fig. 1D), indicating the potential roles of these novel genes in metabolism. We then took Human novel gene 3 (hNovel 3) as an example to assess the function of novel genes in liver cells. hNovel3 was down-regulated in livers of humanized mice treated with FXR agonist and was upregulated in patients with NAFLD (Fig. S1J and Fig. 1E). Pathway enrichment analysis using co-regulated protein coding genes in humanized mice suggested that hNovel 3 may be associated with steroid biosynthesis pathway (Fig. 1F). The specific expression of hNovel 3 gene was supported by Nanopore DRS as well as ATAC-seq and H3K27Ac ChIP-seq. In addition, short-read RNA-seq also corroborated the endogenous structure of hNovel 3 in either humanized livers or primary human hepatocytes (Fig. S1K). To further explore the biological function of hNovel 3, we knocked it down in primary human hepatocytes using three different siRNAs and then profiled gene expression. GSEA pathway enrichment analysis demonstrated that hNovel 3 negatively regulated the Sterol Biosynthetic Process, which was consistent with aforementioned pathway analysis in humanized mice (Fig. 1G).

By performing DRS on the livers of humanized mice under pathophysiologically relevant metabolic conditions, we have constructed a novel and comprehensive annotation of human liver transcriptomes reflecting these important metabolic responses. This effort has led to substantial expansion of human liver transcriptome identifying a large number of novel genes and transcripts. Our results also reinforced the notion that the current annotation of human liver transcriptome is far from being complete, and indicated that a de novo annotation of human liver transcripts reflecting key metabolic responses needs to be first established before transcriptome dynamics underlying these responses can be sufficiently studied.

### Human liver transcriptome dynamics in response to representative metabolic treatments

Equipped with this comprehensive annotation of human liver transcriptome encompassing transcripts that are expressed under representative metabolic treatments, we next examined how these treatments regulate gene expression at transcript level in human liver cells in an in vivo setting. Strikingly, the vast majority of significantly regulated transcripts displayed no corresponding gene level changes in any of the treatments: Fast, PPARα, PPARγ and FXR agonists. For example, less than 6% of the differentially regulated transcripts manifested gene-level changes in response to PPARα agonist treatment (Fig. 2A and Table S4-5). The percentage of genes that were consistently regulated at both gene and transcript levels were also similarly low for all other treatments (Fig. 2A). As an example, Glutathione S-Transferase Mu 1 (GSTM1) gene, a central regulator in the detoxification of electrophilic compounds, produced 20 transcript isoforms. While the levels of two isoforms were significantly changed in response to PPARα agonist, the remaining transcripts and the total gene expression level showed no change (Fig. 2B). Among these, transcript 913293c8 was the highest expressed one and was different from the one that is currently documented in the reference annotation, an observation that was also supported by an independent liver DRS dataset (BioSample, SAMD00127219). Interestingly, this new isoform was predicted to encode a 154 amino acid (aa) protein which was much shorter than the one in reference annotation (ENST00000309851, 219 aa) (Fig. 2C). Remarkably, three-dimension structure alignment analysis showed that the new isoform resulted in the loss of a large domain compared to the reference transcript (ENST00000309851) (Fig. 2D). This example illustrated the unique advantage of transcript-level analysis in detecting gene expression regulations of important functional consequence that are otherwise non-detectable by gene-level study.

**Fig. 2.**
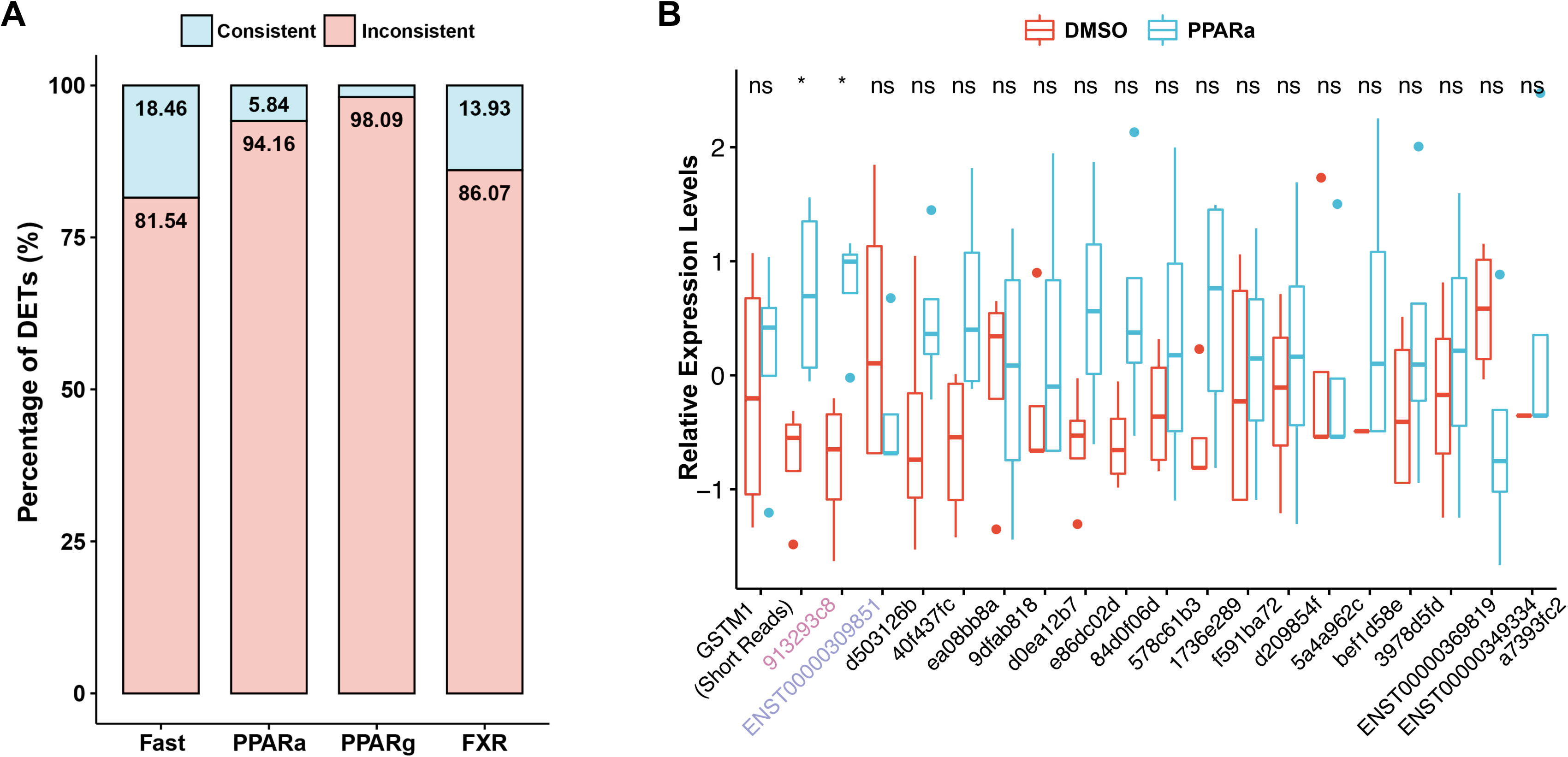
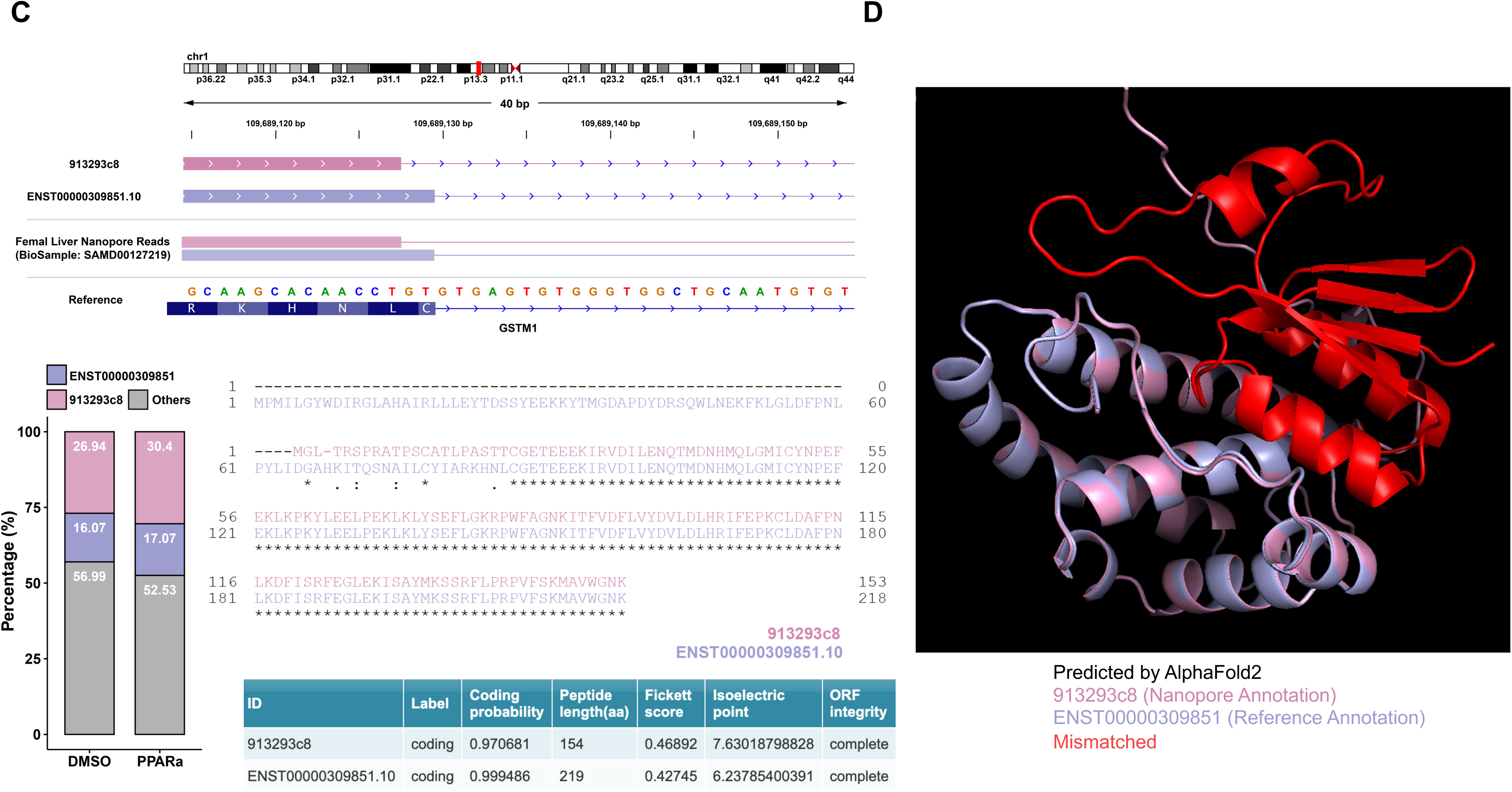
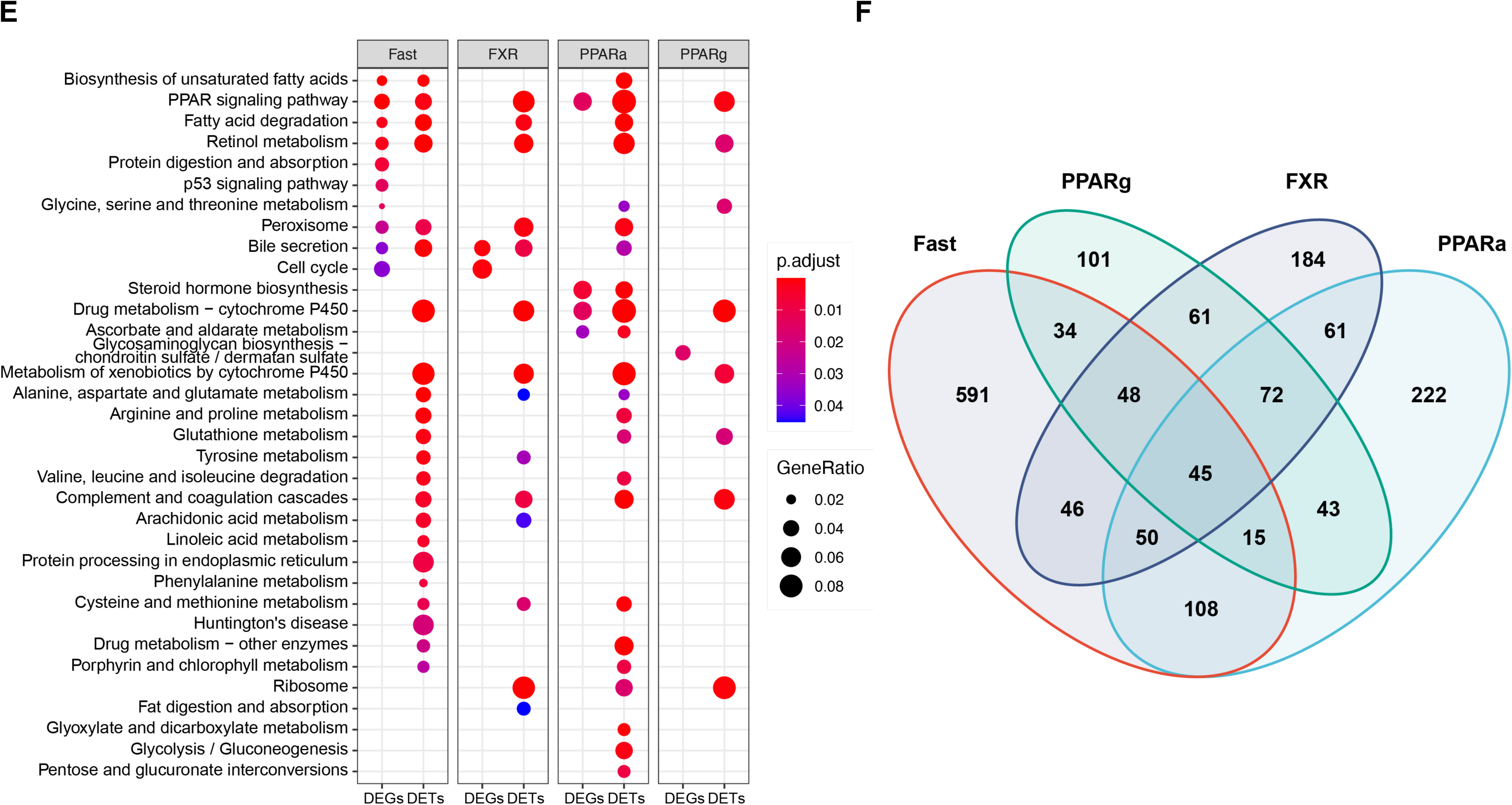
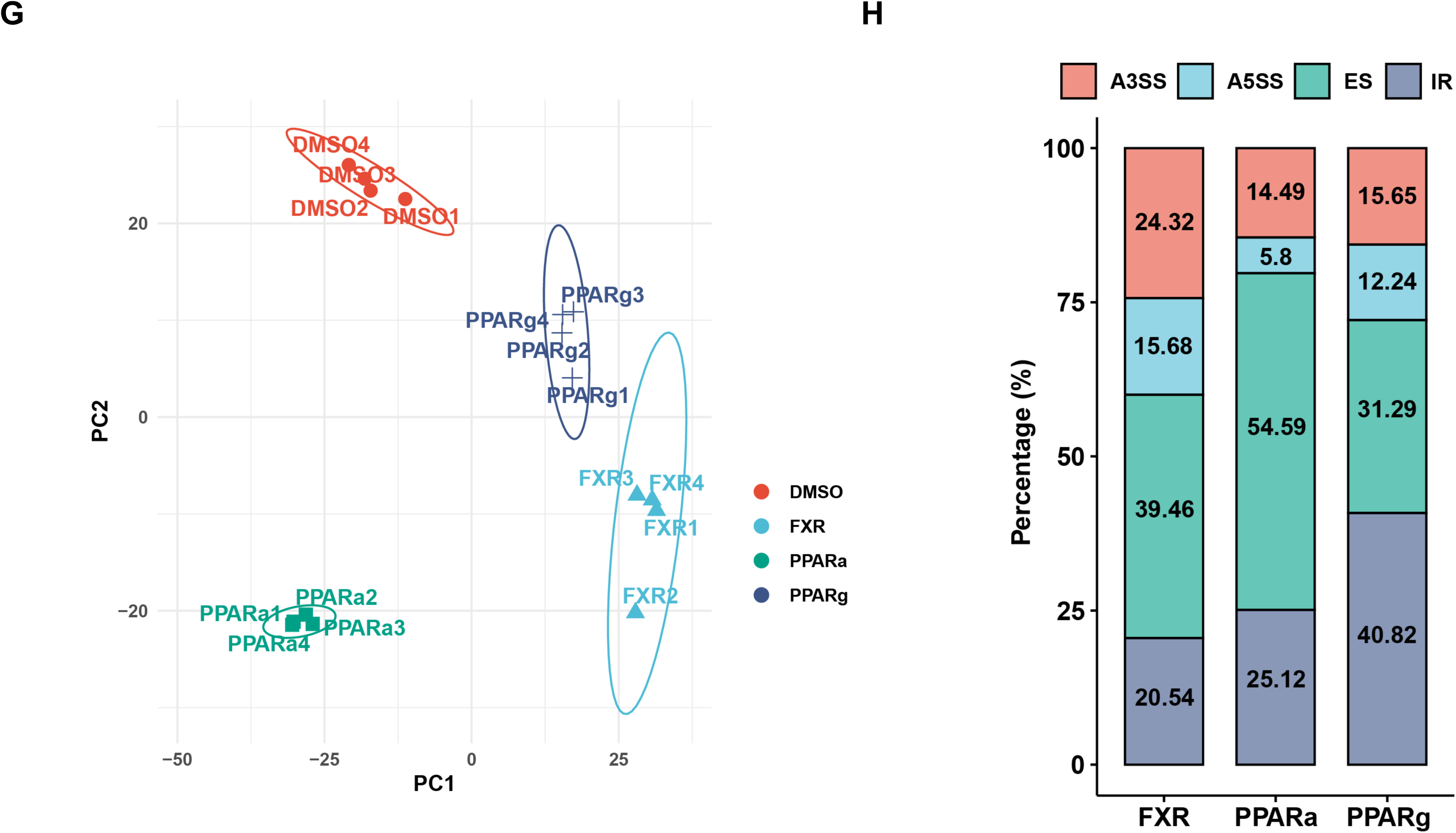
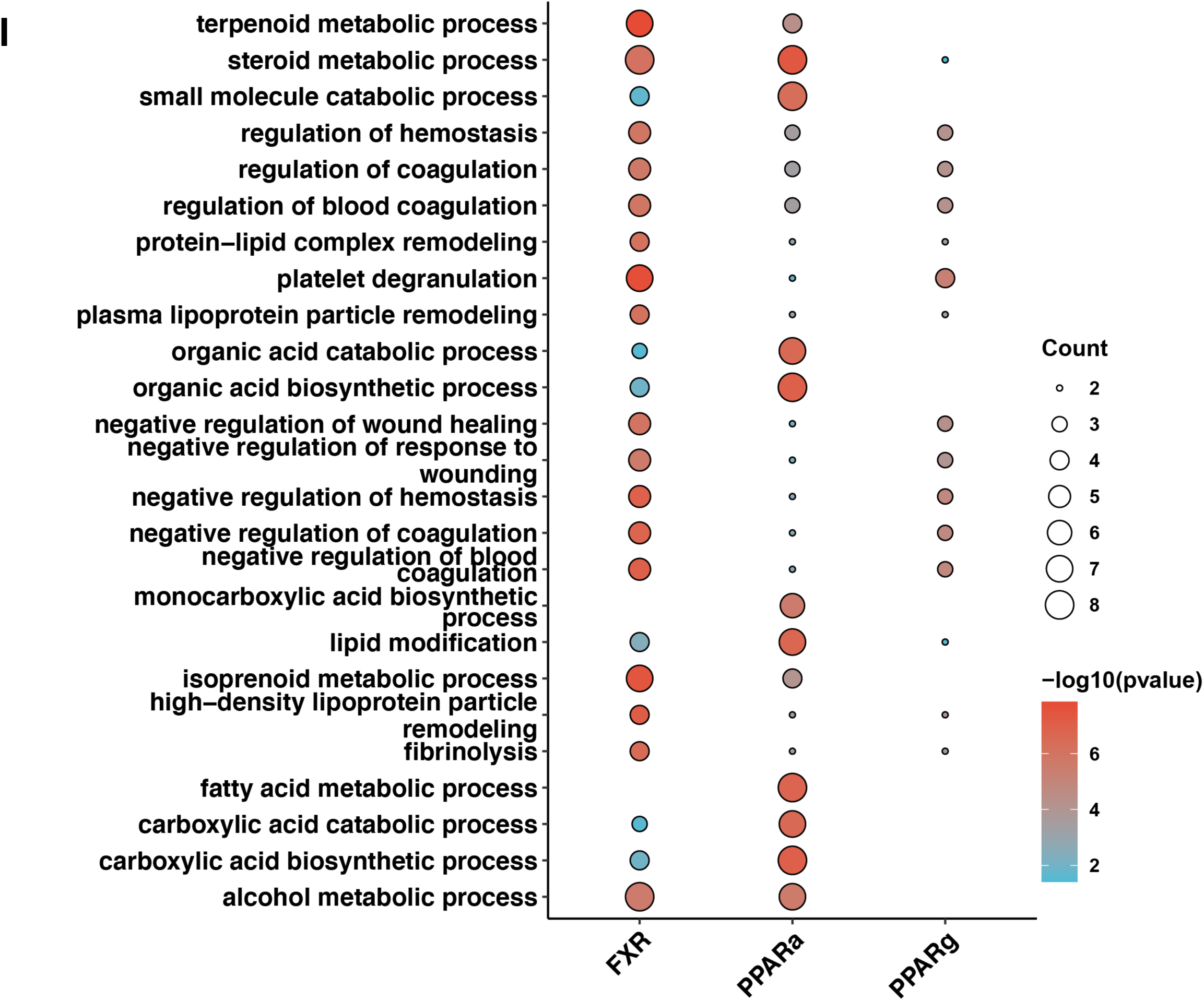
Human liver transcriptome dynamics in response to representative metabolic treatments. (A) Percentage of differentially expressed transcripts (DETs) with consistent gene level regulation patterns in short read RNA-Seq. The DETs that had the same fold change directions with the DEGs were referred to be consistent (Consistent), whereas the others were inconsistent (Inconsistent). A log2 (fold change) more than 0.5 and a p value less than 0.05 were used to define the DETs and DEGs. (B) Boxplot of GSTM1 gene and transcript expression levels in PPARα agonist and DMSO treatment samples. The GSTM1 gene expressions were measured by short read RNA-Seq and the transcript expressions by DRS. The two significantly changed transcripts were highlighted by purple and dark blue respectively. * p<0.05, two-tailed unpaired Student’s t-test. (C) Top: Isoform schematic of GSTM1 significantly changed transcripts (913293c8 in purple and ENST00000309851.10 in dark blue) from humanized mouse liver DRS and public human liver dataset (BioSample: SAMD00127219). Left bottom: The percentages of transcript expression levels of 913293c8, ENST00000309851.10 and other transcripts in GSTM1 gene. Right bottom: Table of the coding protein potential analysis results. Right top: Alignment of the protein sequence between 913293c8 and ENST00000309851.10. The matched aligned amino acids were marked with *. (D) Alignment of the three-dimensional protein structures of ENST00000309851 (dark blue) and 913293c8 (purple). The mismatched zones were highlighted by red. (E) Comparisons of the top enriched pathways for DETs and DEGs under different treatments. The pathway gene sets were downloaded from the Gene Ontology biological process (GO BP) database (http://geneontology.org/). (F) Venn diagram of overlapping human DETs in humanized mouse livers under various conditions. The DETs were defined as | log2 (fold change) | > 0.5 and p value 0.05. (G) PCA plot of human splicing events across samples treated with different transcription factor agonists. (H) Composition of significantly changed alternative splicing (AS) types, which includes exon skipping (ES), alternative 5’ splice site selection (A5SS), alternative 3’ splice site selection (A3SS), and intron retention (IR), during treatments with FXR, PPARα, and PPARγ agonists. (I) Comparisons of the top enriched pathways (GO BP) for transcripts containing AS events that were significantly changed between treatments.

Remarkably, pathways that were enriched by differentially expressed transcripts (DETs) and differentially expressed genes (DEGs) also displayed marked differences (Fig. 2E). For example, in the Fast treatment, DETs and DEGs shared several essential pathways such as PPAR signaling pathways, fatty acid degradation and bile secretion which were widely known to be regulated by Fast treatment. However, the dynamic transcripts induced by Fast also indicated important roles in certain crucial biological processes such as glutathione metabolism, tyrosine metabolism and linoleic acid metabolism, none of which were detected in DEG analysis. These data highlighted that transcript-level regulations revealed by DRS could uncover substantially more information on gene regulations that are often undetectable by gene-level analysis.

Metabolic processes in the body are intertwined, and metabolic genes are often regulated by multiple metabolic stimuli, a notion called metabolic sensitivity which can be used to identify genes, particularly lncRNA genes, that play a role in metabolism^15,29^. Hence, we examined the transcript-level regulations in response to different metabolic treatments and identified the commonly changed transcripts that may have greater chance to play a role in metabolism. We noticed that a sizable portion of transcripts were regulated by at least two treatments showing the high sensitivities to metabolic stimuli (Fig. 2F). Among these, 45 transcripts were changed in all treatments which were mainly enriched in glutathione related pathways, xenobiotic catabolic process and sulfur compound metabolic process (Fig. S2A). These commonly regulated transcripts displayed divergent regulatory patterns in response to different treatments (Fig. S2B). In addition to the commonly regulated transcripts, we also observed hundreds of transcripts that were specifically regulated by each treatment (Fig. S2C) and they were enriched in diverse metabolic pathways (Fig. S2D).

Alternative splicing (AS) is one of the most important regulatory mechanisms in transcripts diversity which plays a central role in regulating transcript expression levels, usages and variants^30,31^. One clear advantage of DRS is its capability to reliably define AS events. We found that the humanized livers with different metabolic treatments split clearly according to AS events in PCA analysis, which indicated that they were subject to specific regulation by these treatments (Fig. 2G and Fig. S2E). For example, while the dominant AS event regulated by PPARα agonist treatment was exon skipping (ES) (more than 50%), it was intron retention (IR) in PPARγ agonist treated samples (Fig. 2H). Pathway enrichment analysis showed that the significantly changed AS events were involved in important metabolism pathways and displayed differential enrichment patterns in response to different metabolic stimuli (Fig. 2I). For example, the changed AS in response to FXR activation showed greater enrichment in high-density lipoprotein (HDL) particle remodeling process than those in PPARα and PPARγ agonist treatments did. APOA1, one key regulator in HDL biogenesis, displayed more alternative 5′ splice sites (A5SS) in the end of the first exon during FXR activation which may affect APOA1 expression (Fig. S2F).

By building a de novo annotation of human liver transcriptomes reflecting diverse metabolic conditions and simultaneously comparing gene expressions at the transcript level, we were able to comprehensively profile transcriptome dynamics in human liver cells in response to metabolic treatments that are relevant to metabolic regulation and diseases. This analysis uncovered substantially more changes in gene network than those detected by traditional gene-level comparisons and also provided an accurate assessment of how AS contributes to transcript diversity in response to metabolic stimuli. Our results support that an accessible humanized system coupled with contemporary DRS technologies could provide a robust solution to the existing challenges in defining transcriptome dynamics in human liver under a physiological context.

### Dynamics of m6A modification and Poly(A) tail length of RNA transcripts in humanized livers

In the past decade, RNA modifications, such as m6A and changes in poly(A) tail length, have emerged to be key regulators of RNA processing, stability and translation^32–34^ and have been shown to play diverse roles in metabolic regulation^8–11^. But their dynamic changes in human liver in response to metabolic stimuli have never been comprehensively examined due to the inaccessibility of human livers to treatments and current technical limitations in detecting them. For example, most of the transcriptome-wide knowledge of m6A modifications were generated from antibody-based immunoprecipitations followed by short-reads RNA-seq (eg, meRIP-seq)^35,36^. These methods require reverse transcription of antibody-precipitated RNAs to cDNAs which are subject to PCR amplification biases and isoform ambiguity^37^. In addition, limited by the SBS sequencing methods of short-read sequencing, it is also challenging to detect the Poly(A) homopolymers which can be several hundred nucleotides long^38,39^. Different from these methods, Nanopore DRS is a novel real-time, single-molecule method that is capable of detecting these modifications on native RNAs with greater accuracy^40–42^.

We used DRS to analyze m6A modifications on all human RNA transcripts in the humanized livers^43^ and also defined how metabolic treatments regulate these modifications. Our results showed that the modification sites were enriched on CDS and 3’ UTR regions of human transcripts with a clear peak at the right end of CDS and the start of 3’ UTR (Fig. S3A), a pattern that was consistent with previous studies^35,44^. Interestingly, both dietary interventions and transcription factor activations shifted the modification distributions towards 3’UTR regions compared to the control groups (Fig. S3A). Moreover, PCA analysis of m6A modifications on 3’UTR showed significant difference among different treatments but no such distinction existed in the CDS regions of the transcripts (Fig. 3A and Fig. S3B). Using dietary intervention as an example, we further examined the genomic distribution of genes whose transcripts carried m6A modifications, and found that these genes were not evenly distributed on chromosome. Transcripts from certain genomic regions showed intensive modifications but others were sparse (Fig. S3C). While most of the m6A sites existed in both AL and Fast treatments, treatment-specific modifications also occurred, and Fast-specific modifications were generally closer to 3’ UTR compared to those in the AL group (Fig. 3B). To further understand the dynamics of m6A modifications during Fast, we performed differential modification analysis and identified 302 significantly changed modification loci. More than one third of these loci contained motif GGACA and more than 40% of them displayed transcript-or gene-level changes in Fast treatment suggesting m6A modifications on these RNA transcripts could modulate their expression levels (Fig. 3C and Table S6). Pathway analysis indicated that these differentially regulated m6A modifications occurred on transcripts of key metabolic genes in fatty acid and alcohol metabolic processes such as HMGCS2, PCK1 and SCD1 (Fig. 3D).

**Fig. 3.**
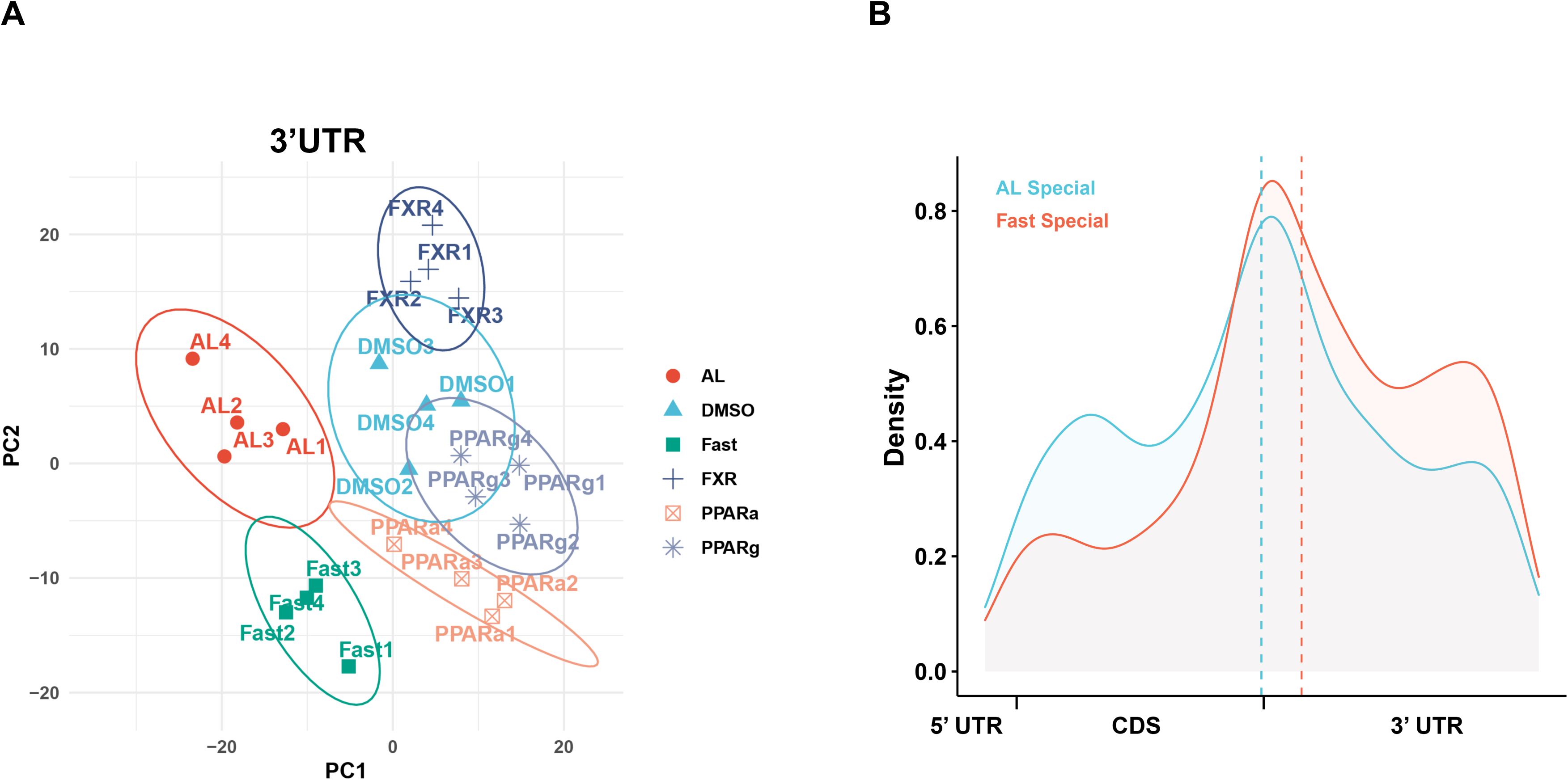
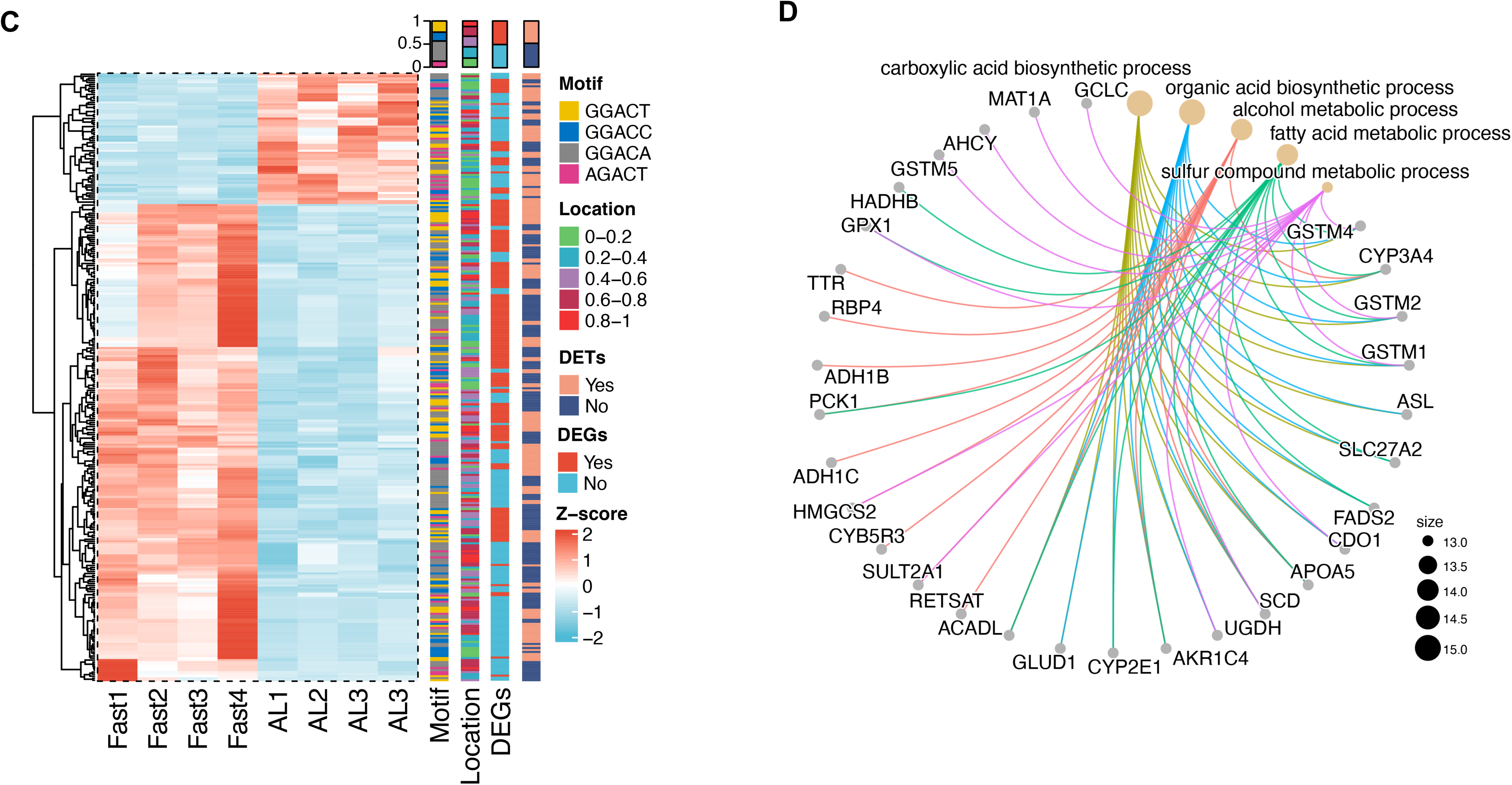
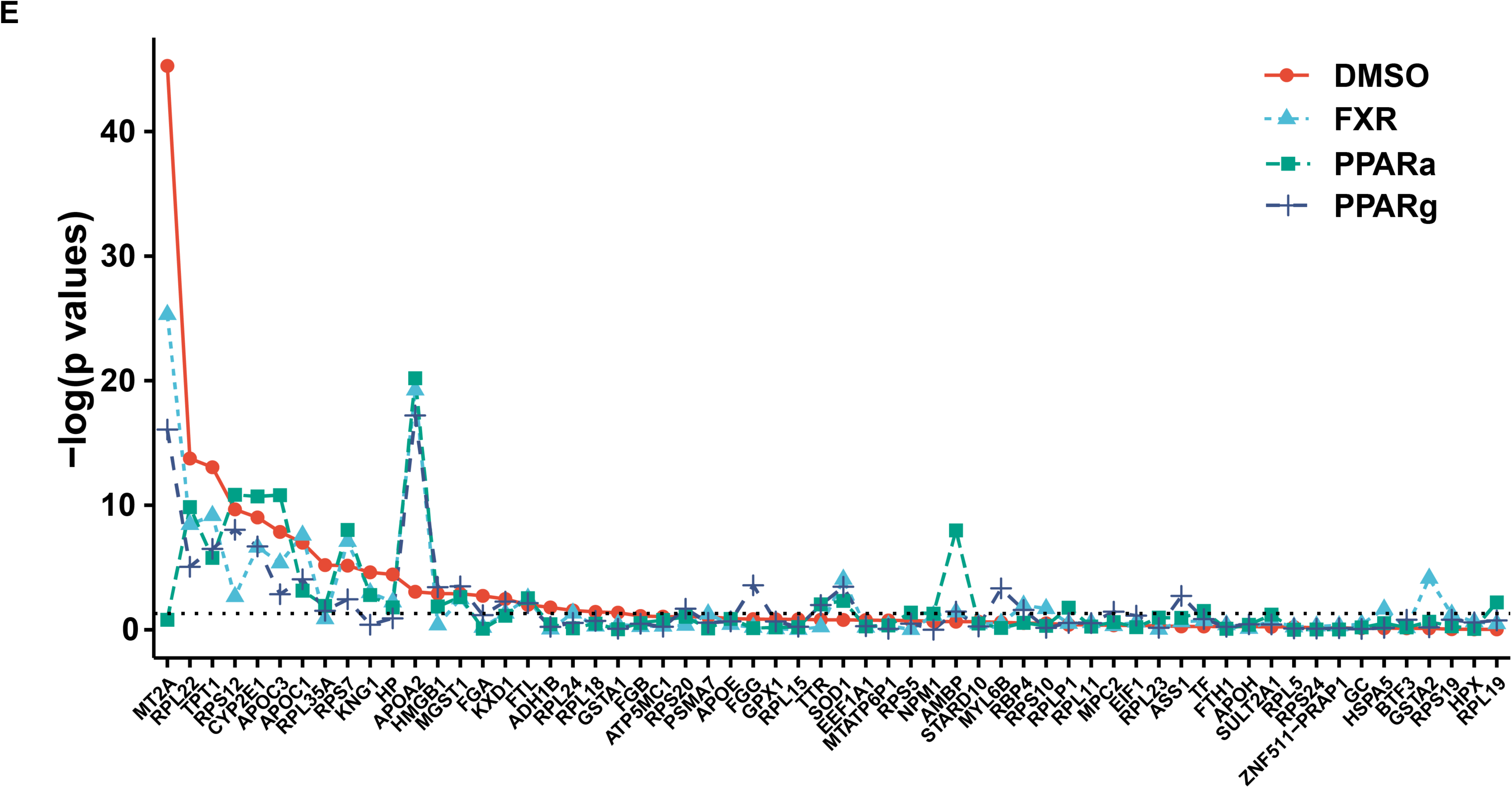
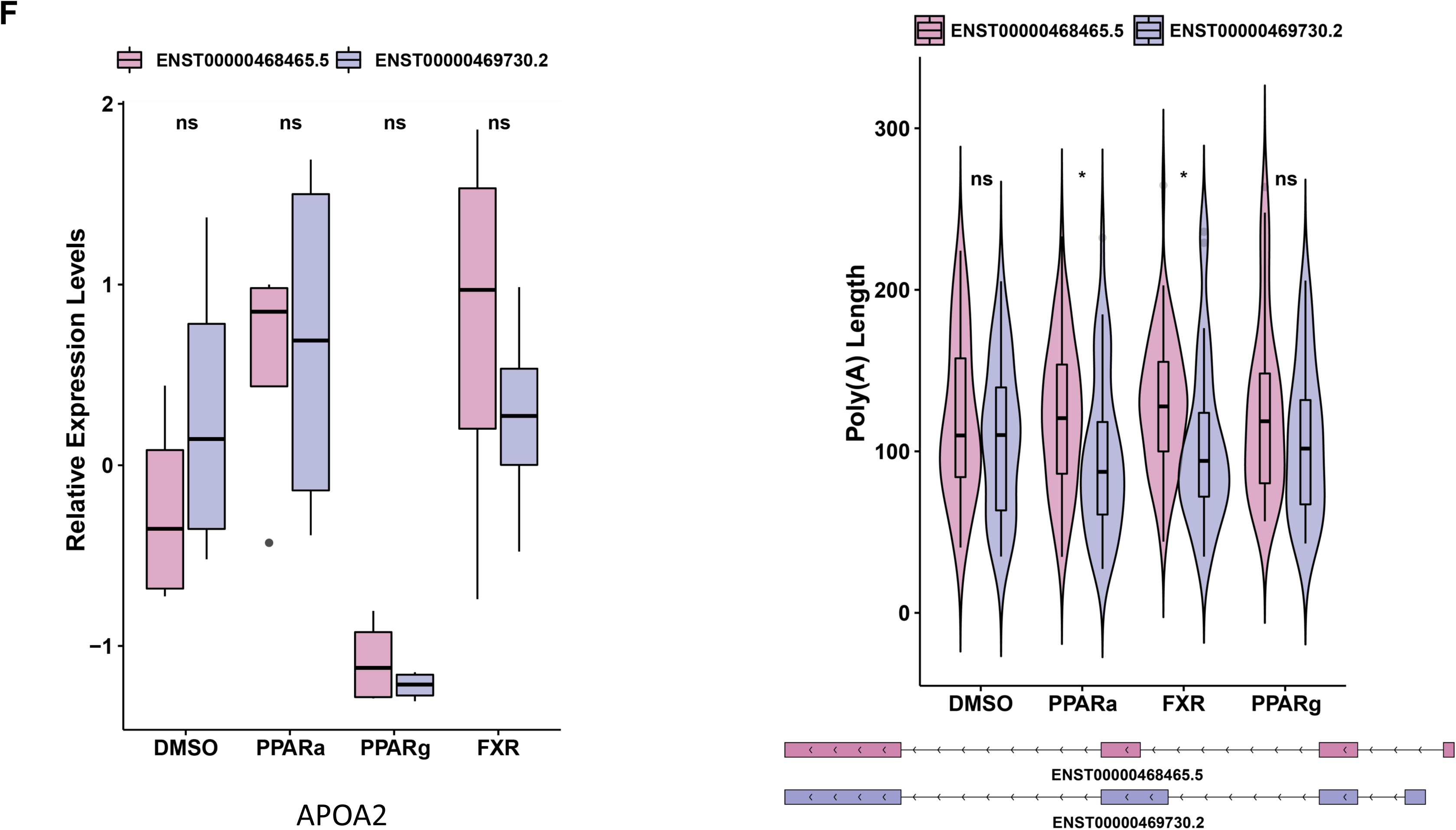
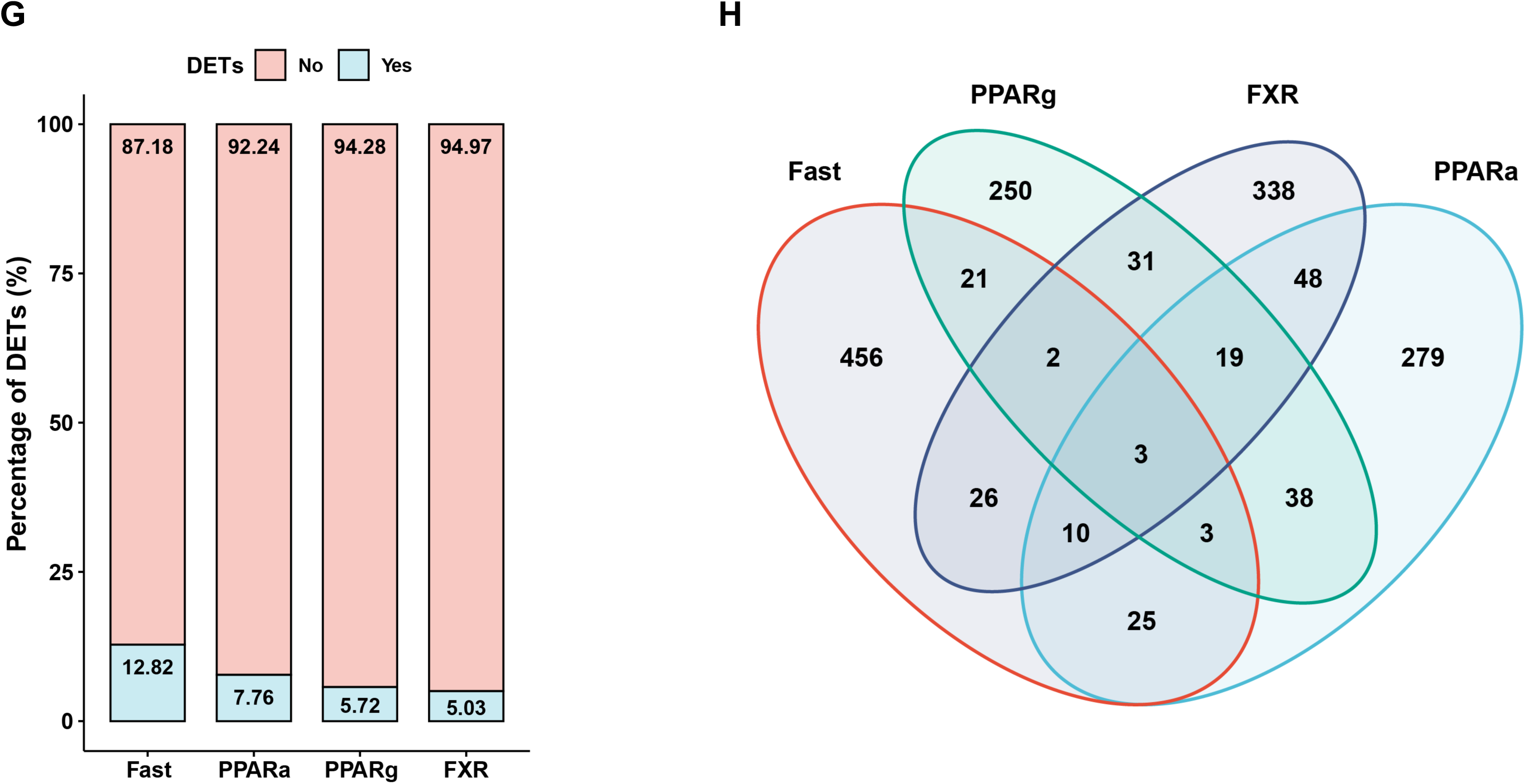
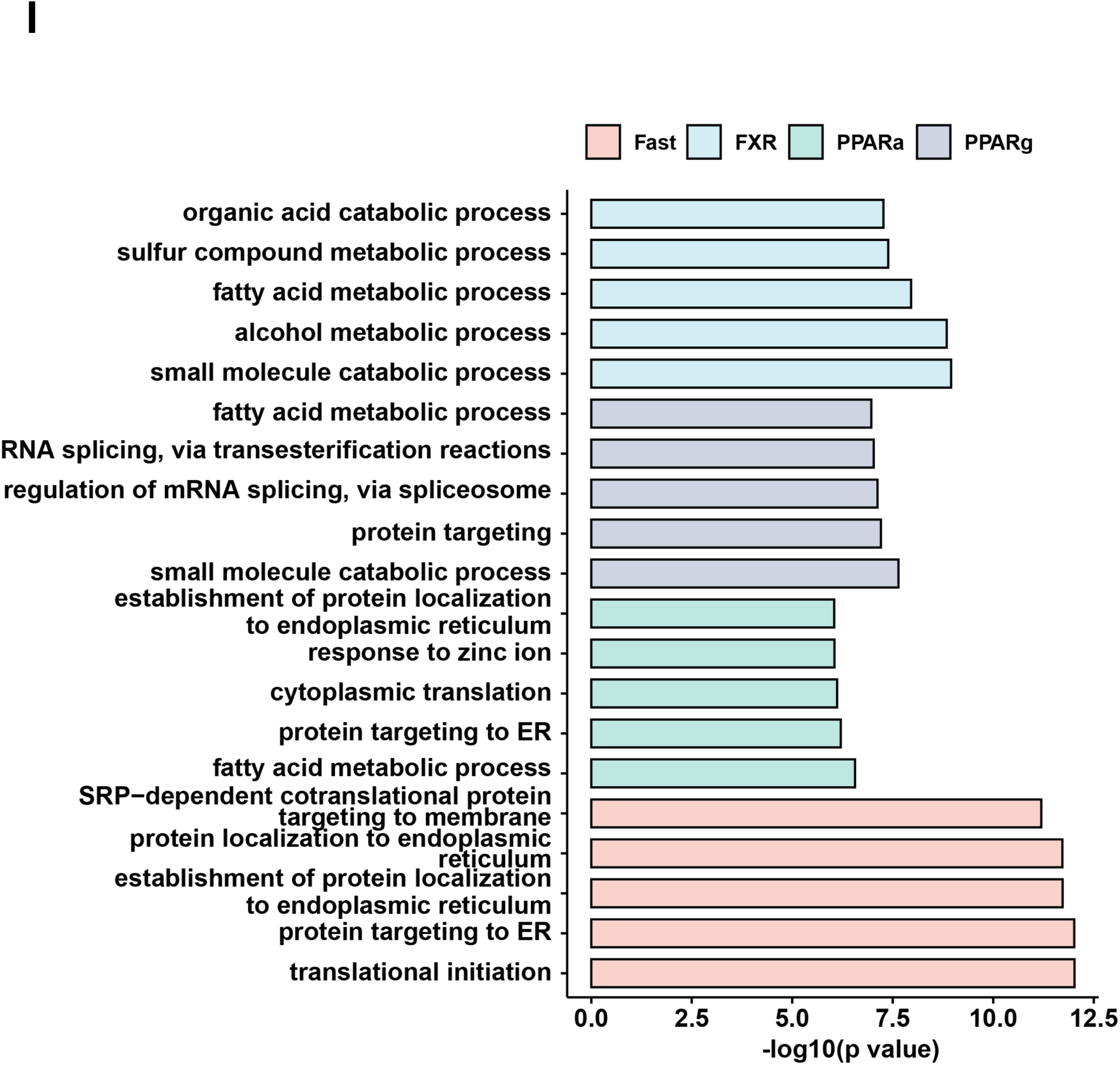
Dynamics of m6A modification and Poly(A) tail length of RNA transcripts in humanized livers. (A) PCA plot of human m6A modification events localized in 3’ UTR regions with different treatments. (B) Metagene analysis of Fast and AL specific m6A modifications in human. (C) Heatmap of human m6A modification frequencies that were significantly changed between Fast and AL treatments. The frequencies of the modification were generated by normalizing to the total modifications in each sample, and were further normalized using the Z-score method across samples. The overlap percentages between DETs and DEGs, as well as the modification motifs and locations (ranging from 0 to 1, representing the 5’ end to 3’ end), were summarized on the right. (D) The hub genes and the central network diagram for top 5 enriched pathways generating from the significantly changed modifications between AL and Fast. (E) Kruskal-Wallis test for poly(A) tail length variance of gene isoforms with different transcript factor agonist treatments. The dashed line labeled the significant bar (p value 0.05). (F) Left: Box plot of relative expression levels of the two APOA2 transcripts (ENST00000468465.5 in purple and ENST00000468730.2 in dark blue) in different treatments. Right top: Violin plot of poly(A) tail lengths of the transcripts of APOA2 in different treatments. The difference between the two transcripts were analyzed by two-tailed unpaired Student’s t-test. ns p>0.05* p<0.05. Right bottom: Isoform schematic of the isoform structures of APOA2 transcripts. (G) Percentage of DETs in the human transcripts displaying differentially changed Poly(A) tail lengths (DPTLs). (H) Venn diagram of the overlapped human transcripts with DPTLs in different metabolic treatments (length difference > 10bp and p value < 0.05). (I) Top 5 significantly enriched pathways (GO BP) for human DPTLs transcripts in different treatments.

Compared to conventional RNA-seq, DRS can also precisely determine poly(A) tail length on transcripts. We quantified the lengths of poly(A) tails on all human transcripts in humanized livers and found that the mean length was around 100 nt (Fig. S3D). Compared to the protein coding RNAs, most of the remaining types of transcripts displayed shorter poly(A) tail length. Surprisingly, the poly(A) tail lengths of lncRNAs were significantly longer than those of protein coding genes (Fig. S3D), suggesting a unique regulatory role of poly(A) tail in lncRNA metabolism and function. We also found that the poly(A) tail lengths of transcripts showed a trend of negative correlation with their expression levels (Fig. S3E), and transcripts containing intron retention usually had longer poly(A) tails (Fig. S3F).

In light of the divergent expression regulations of different transcripts within genes, we explored if poly(A) tail lengths also exhibit transcript-specific regulation in response to metabolic treatments. We checked the variance of poly(A) tail length among the transcripts from the same genes under different treatments and found that the distributions of poly(A) tail length on transcripts varied from gene to gene. For example, while certain genes, such CYP2E1, showed significantly divergent poly(A) tail length among all the transcripts, other genes, such as APOE, displayed similar poly(A) lengths on all transcripts (Fig. S3G).

Furthermore, different treatments regulated the diversities of the poly(A) tail lengths of transcripts from the same gene in a metabolic context dependent manner (Fig. 3E). For example, two major transcripts from APOA2 displayed no expression level changes under any treatment. While the transcripts displayed no significant difference in Poly(A) tail length in DMSO and PPARγ agonist treatments, they had completely different poly(A) tail lengths in PPARα and FXR agonist treatments (Fig. 3F), suggesting that the regulation of hyperadenylation of RNA transcripts is highly condition- and transcript-dependent.

To further identify metabolically sensitive transcripts displaying high degree of dynamics in their Poly(A) tail lengths, we analyzed the tail difference in response to different treatments. Interestingly, most transcripts whose poly(A) tail lengths were regulated by metabolic treatments had similar expression levels under both conditions suggesting a decoupling of the two events (Fig. 3G). Furthermore, only a small percentage of transcripts showed common changes across all the treatments and most of the dynamic poly(A) tails were specifically regulated by specific metabolic treatment (Fig. 3H and Table S7). To ascertain the potential biological functions of these transcripts with poly(A) tail changes under different treatments, we performed pathway analysis and found that FXR, PPARα, PPARγ activation all impacted major metabolic processes (Fig. 3I).

Our data here were among the first to directly characterize m6A and Poly(A) tail length of human liver transcripts in vivo and revealed that both of them are dynamically regulated and could impact diverse metabolic pathways. This information could serve as a useful resource to understand how these modifications impact human metabolism and the development of metabolic disorders. Our results also indicate that many of these modifications are condition-dependent and often exhibit dynamic changes in response to specific metabolic stimuli, and deciphering their physiological role would require an accessible system of human liver which allows for careful analysis of their dynamics under a series of metabolic conditions.

### Divergent transcriptome architecture and dynamics between human and mouse livers

Mouse is the dominant animal model in biomedical research largely due to their low cost and being conducible to genetic manipulation. Information of gene regulation in human livers are often inferred from mouse studies. However, growing evidence indicates that human and mouse often display inconsistent gene regulation^3,4^. Furthermore, the information from existing studies were mostly generated with short-read RNA-seq which mainly focuses on the gene-level analyses, and as shown previously, are insufficient to capture the overall transcriptome dynamics. Here, to further understand the robustness of mouse as a model for human physiology, we compared gene regulations between human and mouse based on transcript-level analysis of gene expressions in humanized livers. Remarkably, we found that human and mouse shared few commonly regulated DEGs (less than 3%) in response to metabolic treatments indicating that homologous genes in human and mouse may undergo substantially different regulations (Fig. 4A). It is challenging to use short-read RNA-seq to compare human and mouse RNA regulation at transcript level due to the redundancy of exonic elements between multiple isoforms of the same homologous genes ^31^. DRS can detect the full-length native RNAs allowing an unambiguous identification of the expressed isoforms and making it possible to investigate transcript diversity for human and mouse homologous genes^45^. In order to compare transcript-level dynamics of homologous genes between human and mouse, we examined the coefficient of variation (CV) of transcript distribution, which reflects the distributions of expression levels of transcripts and the transcript dynamics among genes. This analysis revealed an inconsistent ranking trend of CV value changes between human and mouse homologous genes indicating divergent transcript regulation between the two species (Fig. 4B). For example, CYP2E1, a central regulator of lipid synthesis, displayed high CV values in both human and mouse but the expression levels of their transcripts displayed substantial divergence. More than half of human CYP2E1 transcripts exhibited decreased expression levels during Fast but one transcript, which was the highest expressed on, was up-regulated (Fig. 4C). However, mouse Cyp2e1 transcripts were all increased by Fast treatment. These results suggest that many human and mouse genes may undergo differential and even opposite transcript-level regulations in response to metabolic stimuli.

**Fig. 4.**
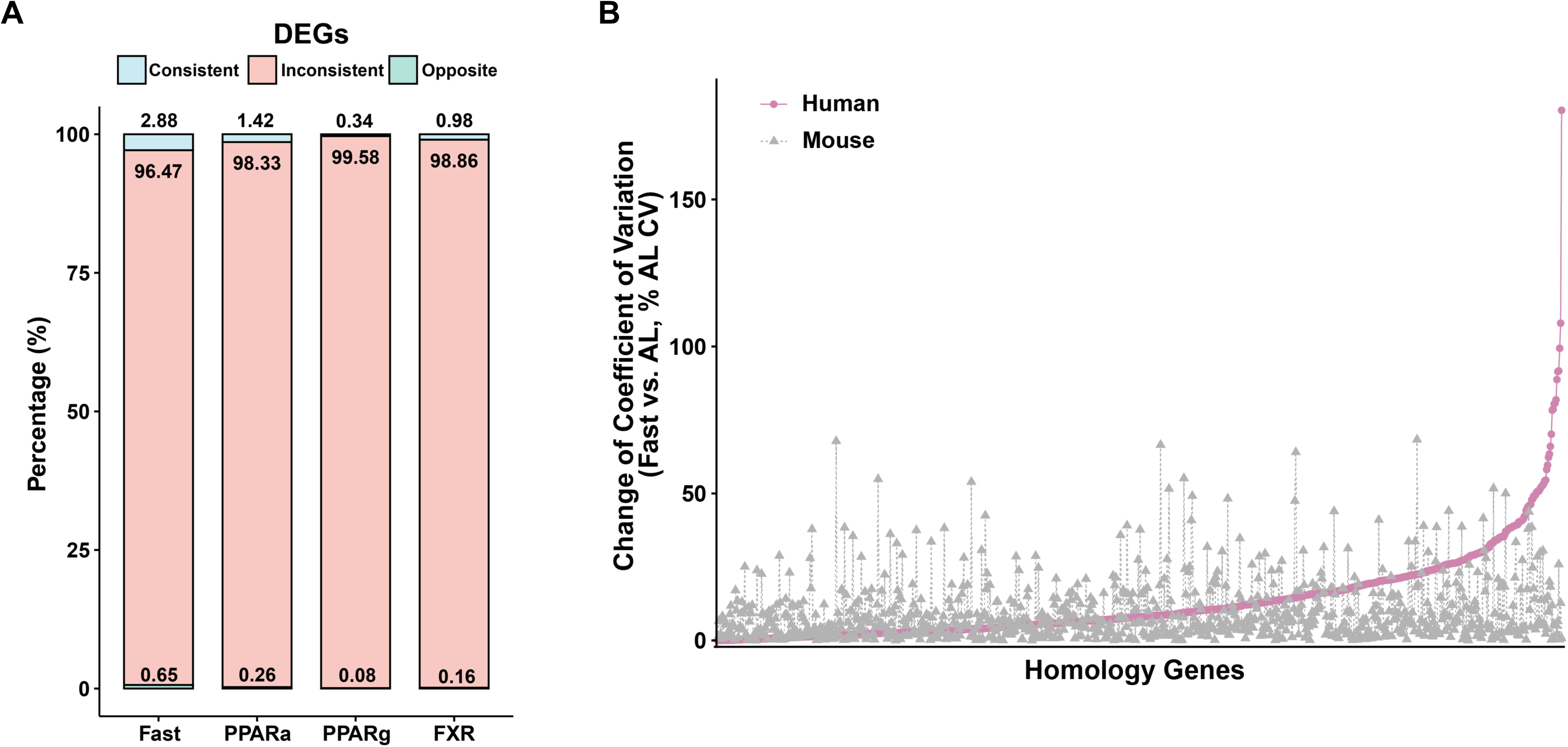
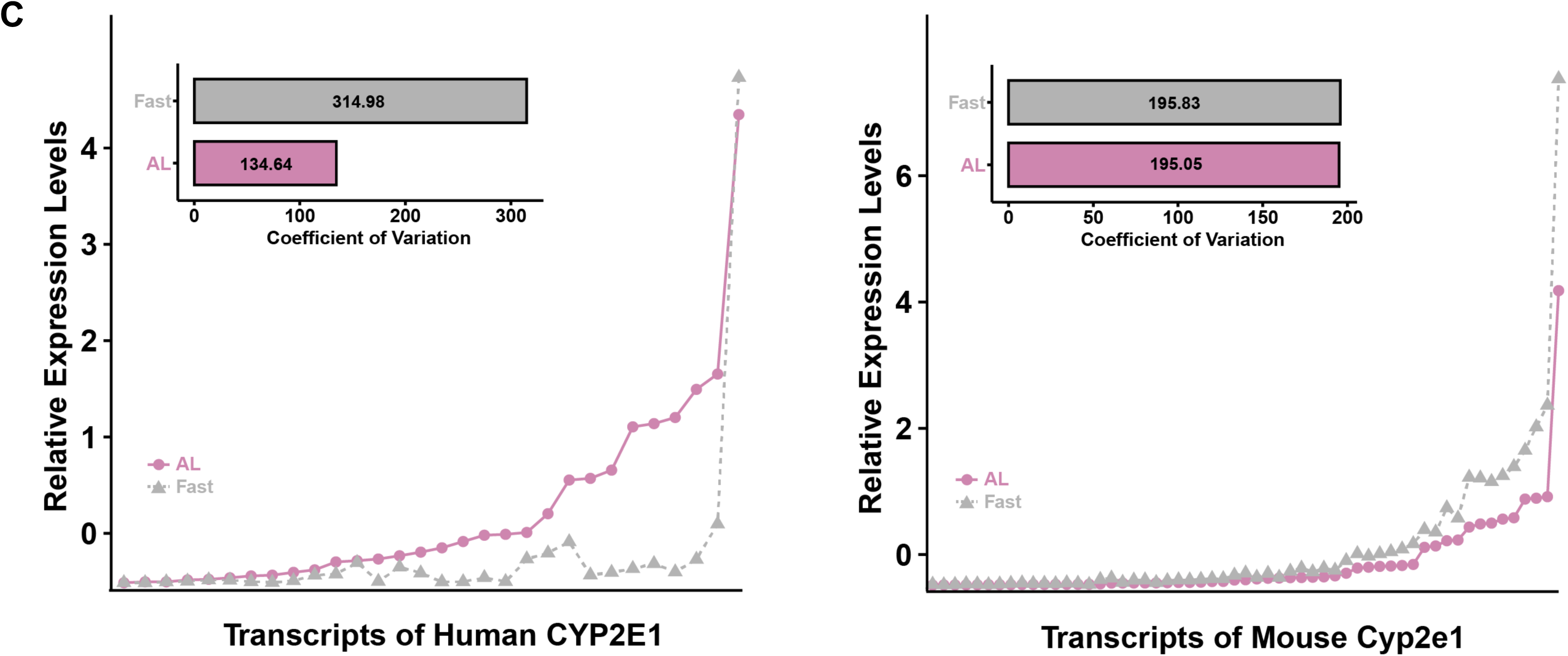
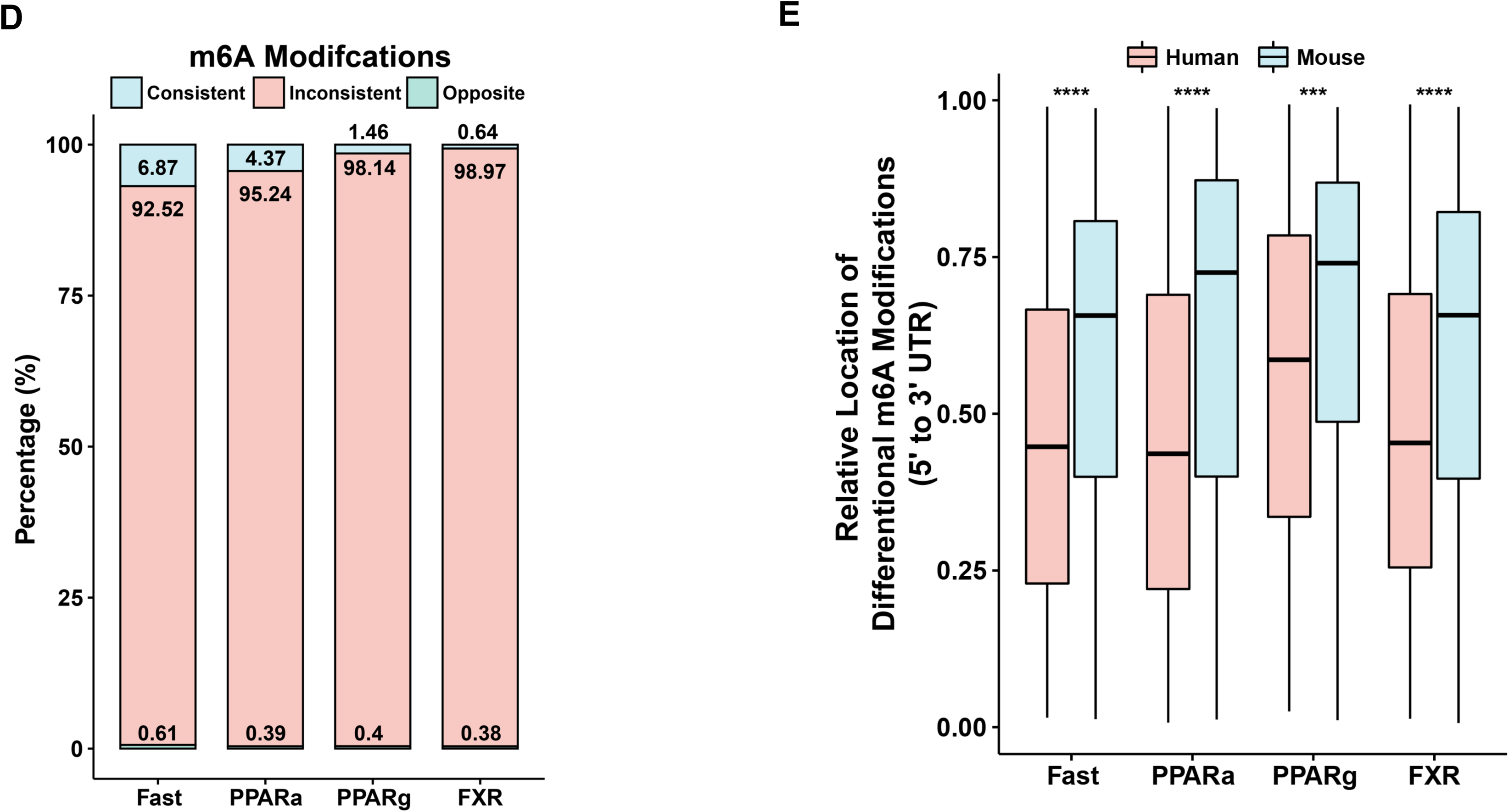
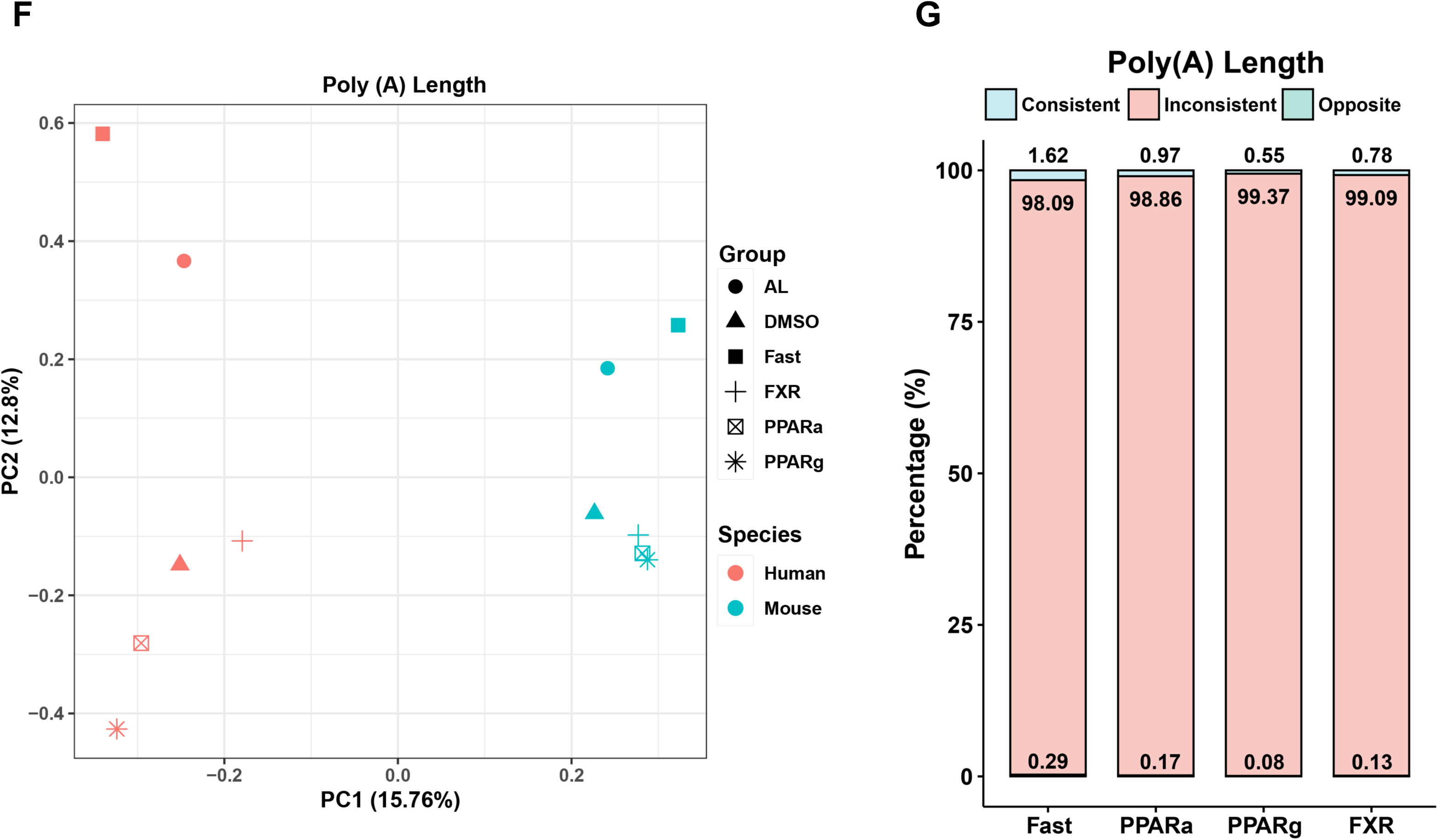
Divergent transcriptome architecture and dynamics between human and mouse livers. (A) The composition of gene level changes for human and mouse homologous genes in different treatments. The expression level of a gene was calculated based on the sum of all transcripts from the gene detected by DRS. For homologous gene comparison, only genes that significantly changed in both mouse and human were considered for consistent or opposite regulation. Homology genes that changed in the same direction were defined as consistent (Consistent), and those that changed in opposite directions were defined as opposite (Opposite). The remaining homologous genes were defined as inconsistent (Inconsistent). (B) The dynamic of coefficient of variation (CV) values for transcript expression variabilities of homologous genes in response to Fast and AL treatment. The homologous gene pairs were ranked by human CV values (human in purple and mouse in grey). (C) Bottom: Dot plots of human and mouse CYP2E1 (Cyp2e1 for mice) transcript expression levels. The transcript expression levels were generated from the mean TPM values from same group. Top: CV values of CYP2E1/Cyp2e1 transcript expression levels in AL and Fast treatments respectively. (D) The compositions of RNA m6A modification changes for human and mouse homologous genes in different treatments. (E) Box plot of relative location of m6A modification loci in human and mouse under different treatments. The modification loci were normalized from 0 to 1 with closer to 1 meaning closer to the 3’ UTR in the gene body. ***p < 0.001, ****p < 0.0001, two-tailed unpaired Student’s t-test. (F) PCA plot of Poly(A) tail length of homologous genes under different treatments in human (red) and mouse (blue). (G) The compositions of gene poly(A) tail length changes for human and mouse homologous genes in different treatments.

Furthermore, the regulations of m6A modifications on homologous protein coding genes also displayed substantial differences in human and mouse (Fig. 4D). Interestingly, the dynamic m6A modification loci in mouse showed a clear shift towards the 3’ end in response to both dietary treatment or transcription factor activations (Fig. 4E). Since this trend was absent in humans, this suggests the m6A modification distribution and regulation machinery may be fundamentally different in the two species.

We also examined poly(A) tail length in human and mouse and found that their overall patterns separated nicely in a PCA analysis indicating clear divergence of poly(A) tail length distribution between the two species (Fig. 4F). Furthermore, genes that had significantly changed poly(A) tail length in response to metabolic stimuli showed little overlap in the two species (Fig. 4G). Pathway analysis indicated that many genes with oppositely regulated poly(A) tail length play a role in critical metabolic processes such as lipid catabolic process (Fig. S4A). For example, while the opposite genes (PRDX6, HSD11B1, PABP1, ECI2 and APOA1) enriched in lipid catabolic process displayed longer poly(A) tail length during Fast treatment in human, they were significantly shorter on corresponding mouse RNAs (Fig. S4B). These data suggested that human and mouse undergo divergent regulation in poly(A) tail length which may underlie the differential gene expression regulations in the two species.

Although mouse studies provide a useful reference for gene expressions in human, our results here indicated that transcriptome dynamics in mouse and human liver cells showed substantial differences at the levels of gene, transcript, m6A and poly(A) length. Thus, it is unlikely that mouse studies could provide the deep specifics of gene regulation that are required to understand the molecular basis of human pathophysiology and such information need to be derived from a human or humanized system reflecting human biology.

### Divergent transcriptome architecture and dynamics between individuals of different genetic backgrounds

Growing evidence supports that genetic background has a strong influence on an individual’s gene regulation and disease susceptibility^46,47^. Due to the inaccessibility of human livers to treatments, however, it is extremely challenging to directly and experimentally examine the impact of genetic background on gene regulation at transcript and RNA modification levels in human liver. Since the humanized mouse model allows us to produce humanized livers of specific genetic background and Nanopore DRS is capable of detecting RNA isoforms and modifications, we used this setting to compare gene regulations in humanized livers of different genetic background.

To evaluate the differences in transcriptomes between individuals, we performed DRS on livers of humanized mice that were generated from a second independent donor and had also been subjected to Fast treatment. We found that approximately 55% of RNA transcripts detected in the second donor (donor2) matched to those in the first one (donor1) (Fig. 5A). This similarity, however, was much higher than that between donor 2 and the GENCODE reference (37%), which is a collective population annotation. Nonetheless, around 20% of transcript isoforms were conclusively different between these two donors (Fig. 5A). Interestingly, only 39 (around 40%) novel genes detected in the donor1 were expressed in donor2 (Fig. 5B). As an example, the full length of the novel gene, ebabaf4b, was detected by Nanopore DRS in donor2 and corroborated by short read RNA-seq peaks (Fig. 5C), but no signal was detectable in the same region in donor1 through either method. This finding supports personalized activation of certain genes which could also explain the absence of certain novel genes in the public reference. In response to metabolic changes, such as Fast, there were only around 10% of the transcripts that were consistently changed in the two donors (Fig. 5D). Furthermore, the overall patterns of enriched pathways based on transcripts specifically regulated in donor1 and donor2 were clearly different (Fig. 5E). To assess the consistency of inter-individual regulations, we further checked the expression levels of commonly changed transcripts during Fast. While most of them displayed consistent regulations in the two donors by Fast, around 20% of transcripts were oppositely regulated (Fig. 5F). We found that transcripts consistently changed in both donors were involved in major metabolism pathways including Organic Acid Catabolic Process, Fatty Acid Metabolic Process and Carboxylic Acid Catabolic Process. Oppositely regulated genes were mostly enriched in inflammatory defense responses such as respiratory burst and defense response to bacterium (Fig. 5G).

**Fig. 5.**
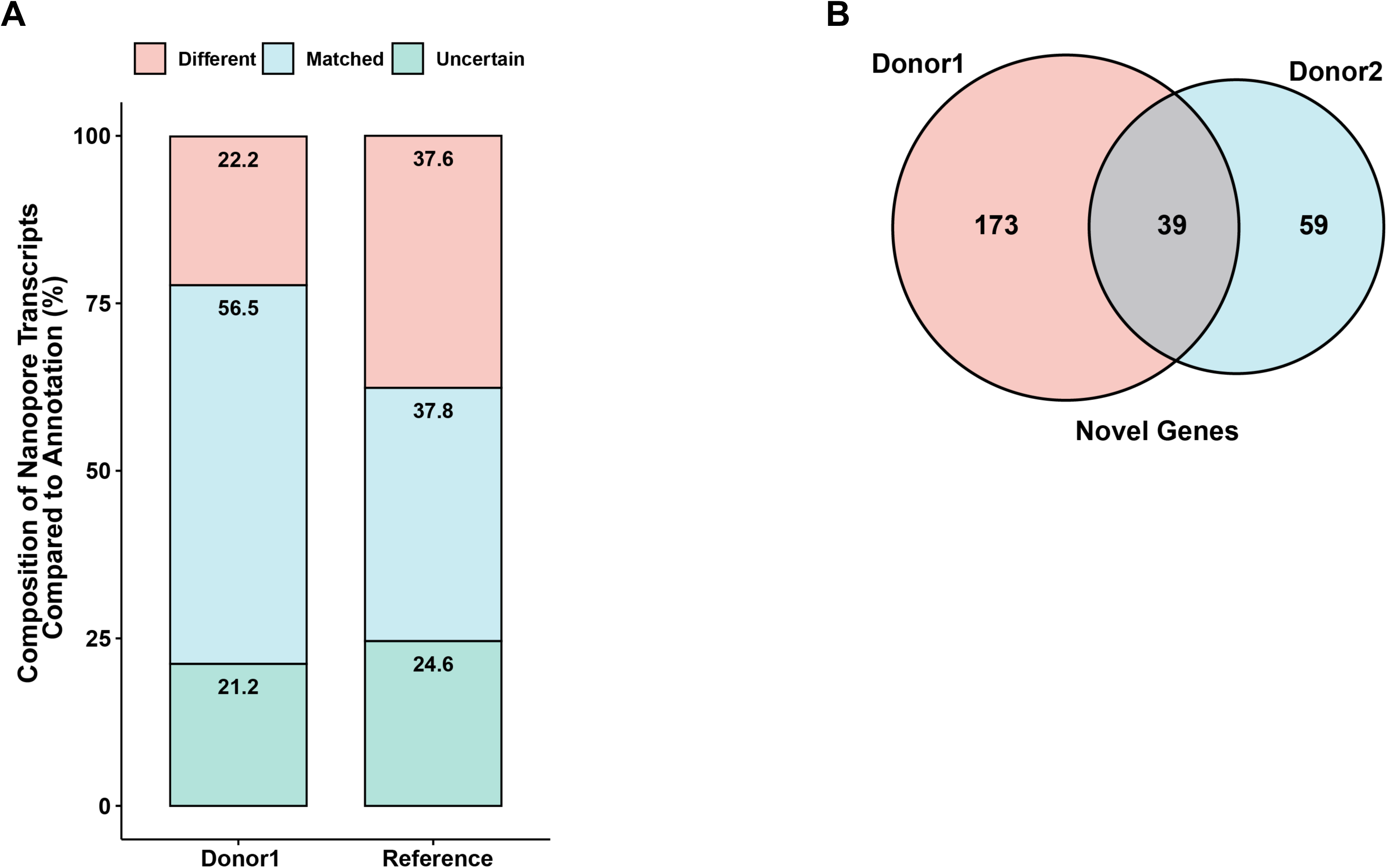
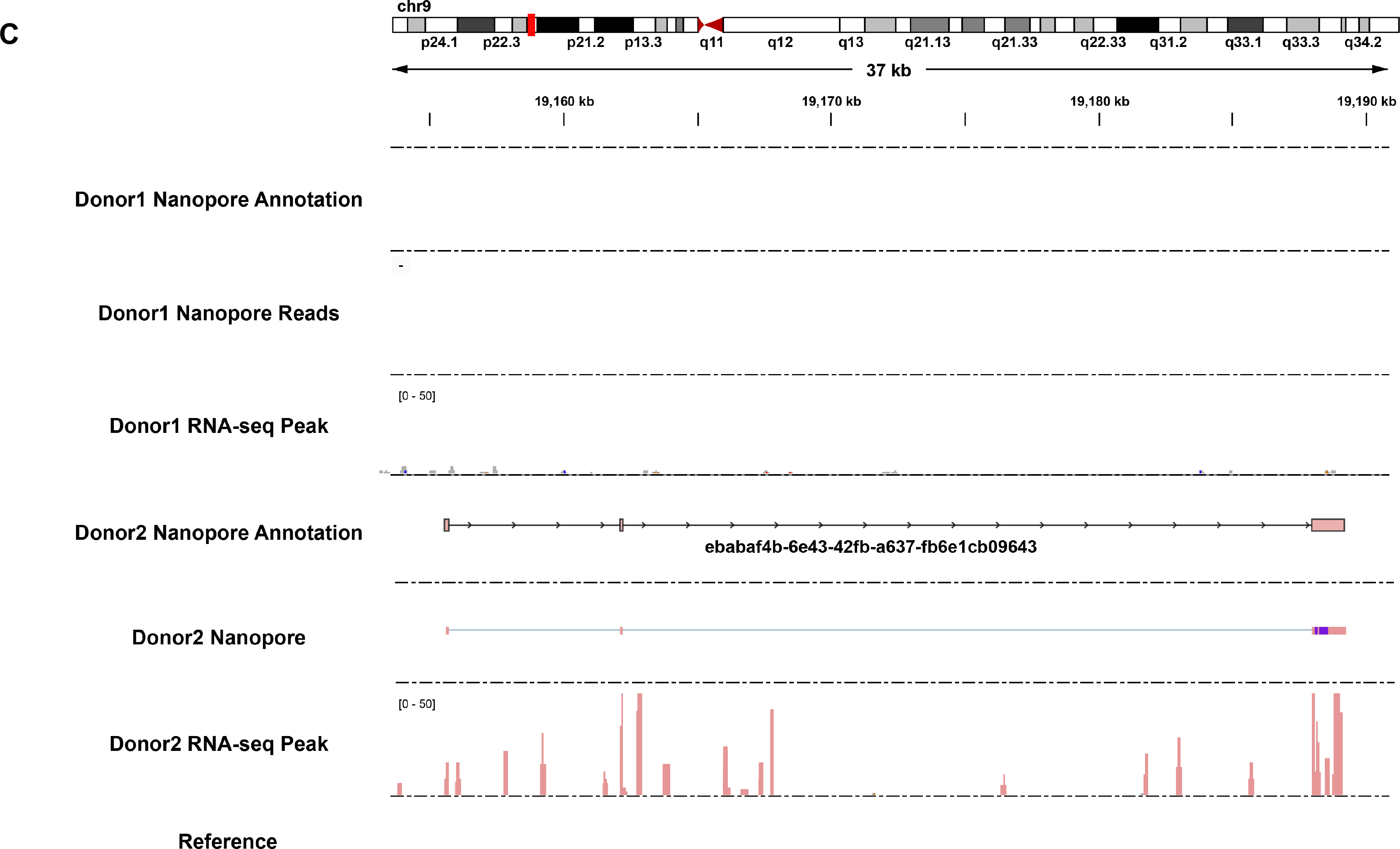
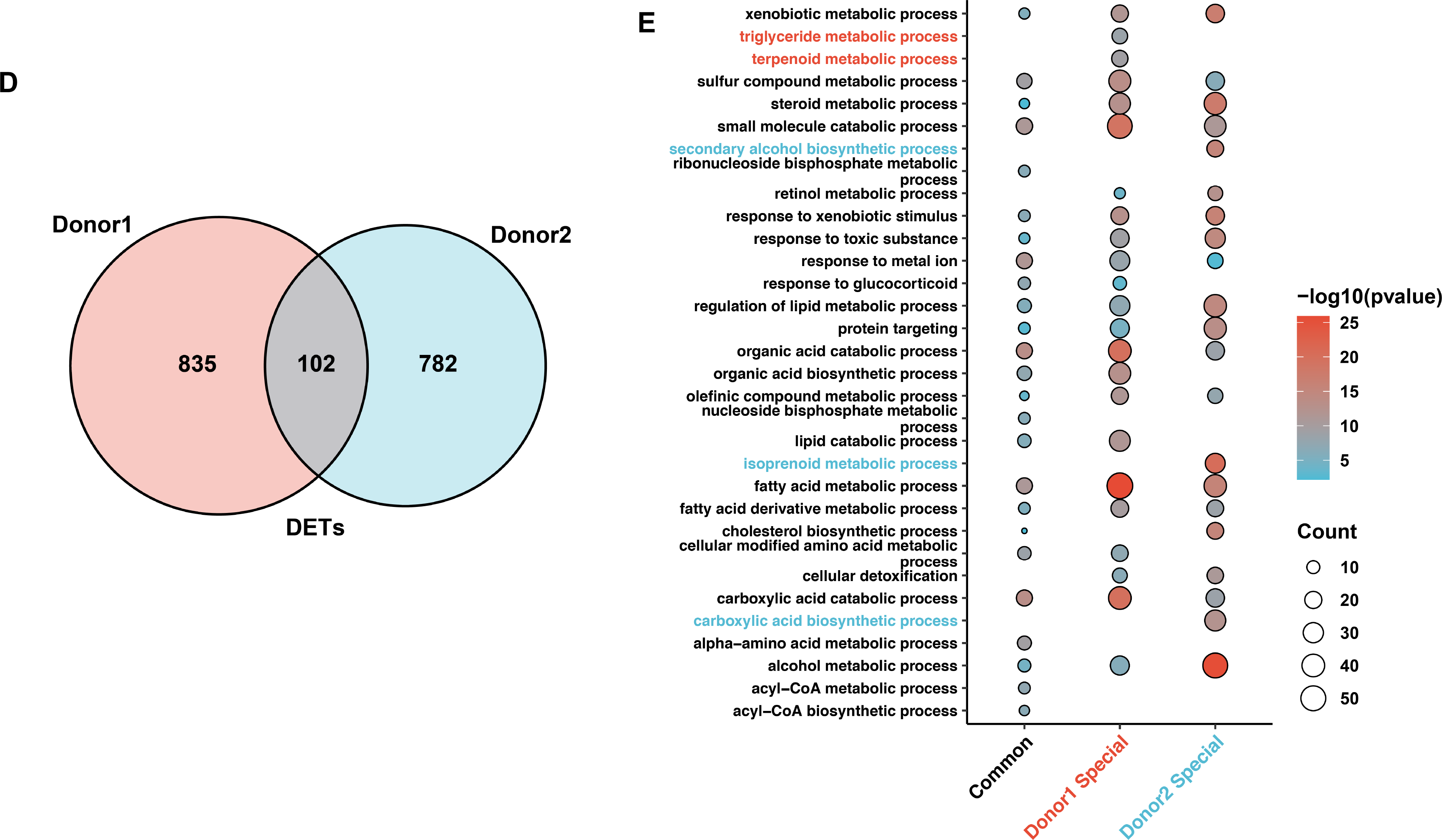
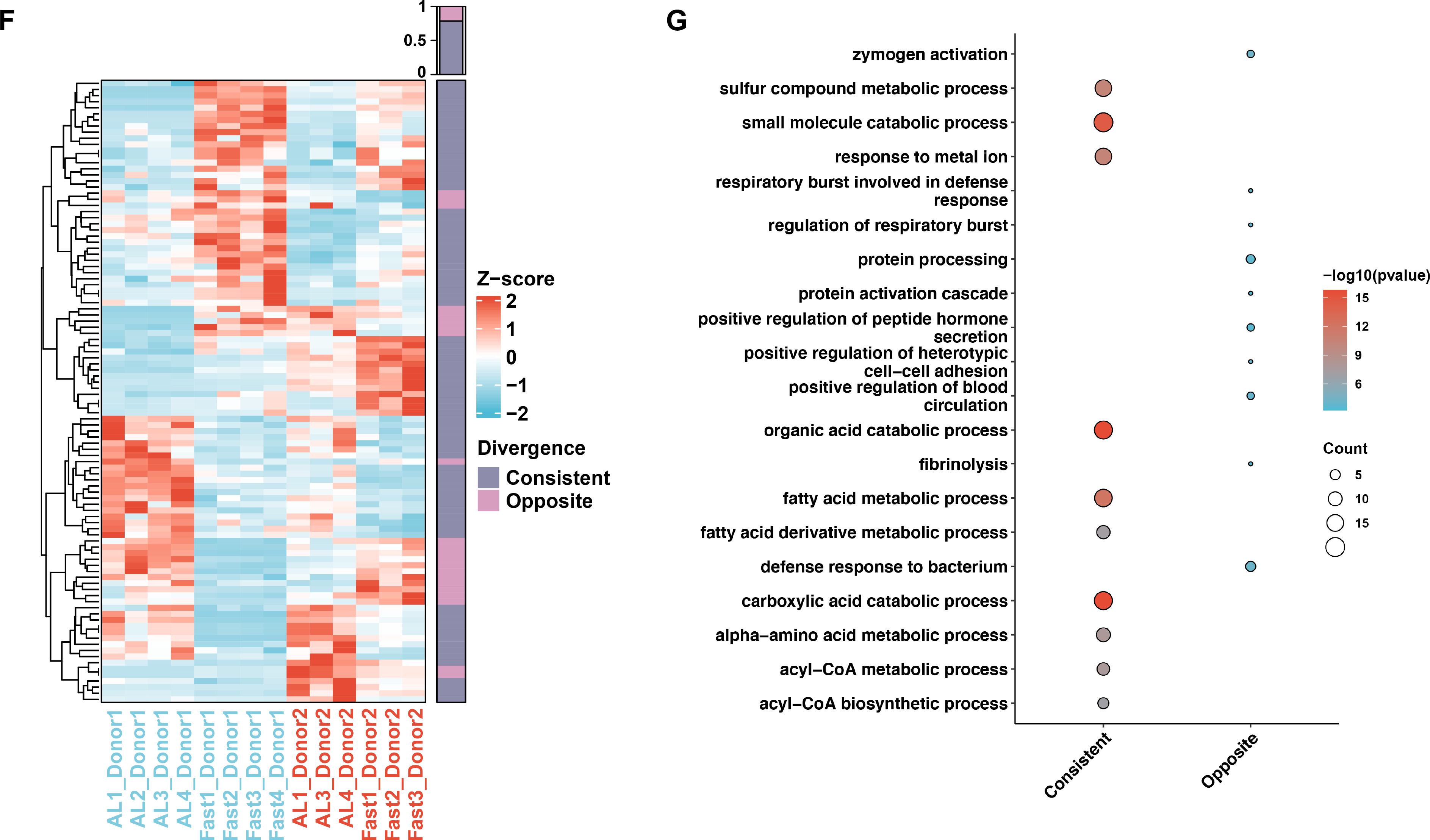
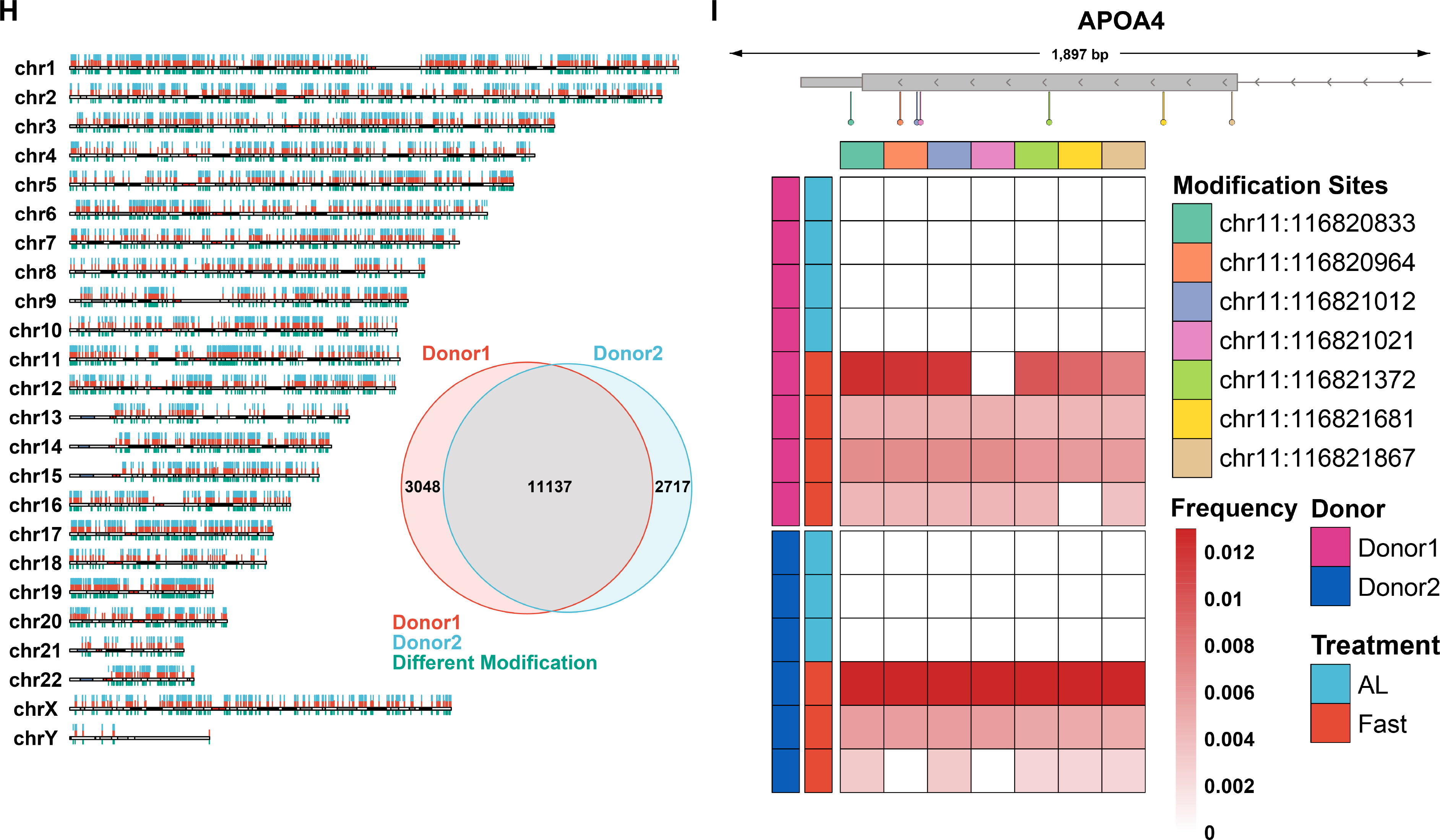
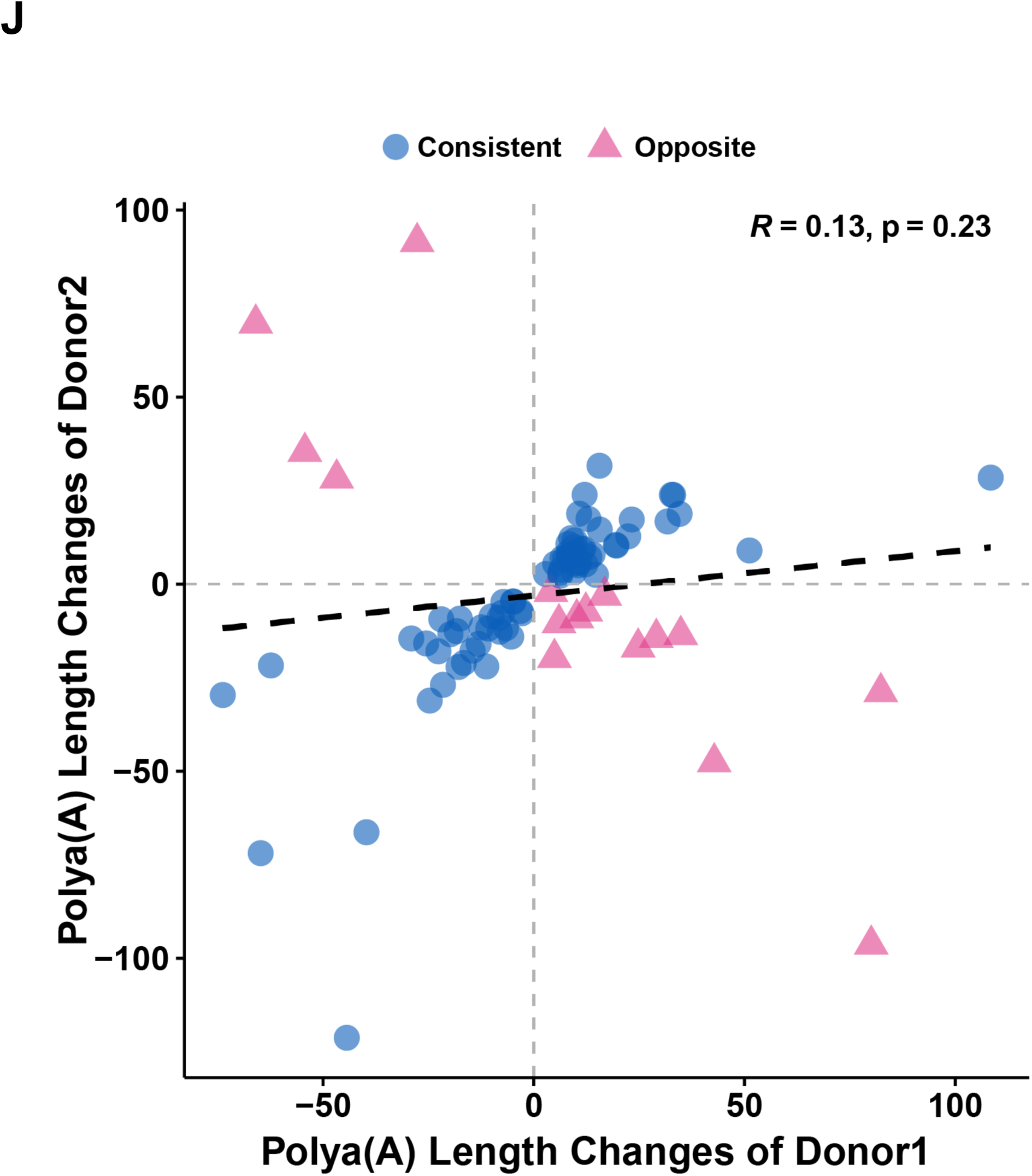
Divergent transcriptome architecture and dynamics between individuals of different genetic backgrounds. (A) Comparison of DRS transcripts between donor2 and donor1 or between donor 2 and the reference annotation. (B) Venn diagram of the overlapped novel genes between donor1 and donor2. (C) Isoform schematic of novel gene from donor2. Donor1 and Donor2’s raw RNA-Seq and DRS reads were also displayed. (D) Venn diagram of the overlapped DETs between donor1 and donor2. Only DETs with the matched isoforms were considered as overlapped ones. (E) Comparison of the top enriched pathways (GO BP) for common, donor1-, and donor2-specific DETs. The donor1 specifically enriched pathways were labeled as red and the donor2 specifically enriched pathways were labeled as blue. (F) Heatmap of the expression levels of commonly regulated DETs in response to AL and Fast treatments in donor1 and donor2. The summary of DETs was on the right. Only DETs with the matched isoforms between the two donors were included. (G) The most significantly enriched pathways (GO BP) for the consistent and opposite DETs between donor1 and donor2. (H) Left top: Distribution of m6A modification sites for donor1 (in red) and donor2 (in blue), with different modifications between them labeled as green. Right bottom: Venn diagram on the bottom right showed the overlapping of m6A modifications between the two donors. (I) Top: Diagram schematic of m6A modification loci on APOA4 gene body. Bottom: Heatmap of m6A modification frequency in APOA4 gene body from donor1 and donor2. The modifications were labeled by different colors. (J) Pearson’s correlation analysis of genes with commonly significantly changed Poly(A) tail lengths in donor1 and donor2. The consistent ones were labeled as dark blue and opposite ones were labeled as pink.

We also compared m6A modifications on human RNA transcripts in the humanized livers of different genetic backgrounds and found that most modification sites (∼80%) could be found in both donors (Fig. 5H). For example, m6A modifications were absent in the 3’ end of APOA4 in AL group and seven of them were induced by Fast treatment in this region for both donors (Fig. 5I). Only 30% of common m6A modifications, however, were regulated similarly in donor1 and donor2, and donor-specifically regulated modifications were related to important pathways such as long-chain fatty acid metabolic process for donor1 and secondary alcohol biosynthetic for donor2 (Fig. S5A and S5B).

We observed a similar trend in the changes in Poly(A) tail length between the donors and only around 10% of regulated transcripts overlapped in donor1 and donor2 (Fig. S5C). Most of the commonly changed genes displayed consistent regulation but a small percentage of them were regulated oppositely in the two donors (Fig. 5J). Pathway analysis showed that donor-specifically regulated genes were involved in cellular pathways such as regulation of mRNA metabolic process in donor1 and mitochondrial translation in donor2 (Fig. S5D).

Our results here illustrated the substantial differences in the composition of genes and transcripts between two individuals of independent genetic backgrounds. Furthermore, gene regulations at transcript, m6A and poly(A) tail length levels were also strongly influenced by an individual’s genetic background. Although comparing gene regulations in patient samples based on a reference genome might be able to detect commonly regulated genes, which accounted for a small fraction of regulated genes, substantial information would have been lost and the overall picture would not be complete. Thus, our result supported an individualized approach in which the changes in global gene network reflecting a patient’s pathophysiology needs to be defined in an experimental system that can define transcriptome dynamics in the context of a specific genetic background.

## Discussion

In this study, we combined an isogenic humanized mouse model and Nanopore DRS to establish a de novo annotation of human liver transcriptome and subsequently studied the regulation of all detectable liver RNA transcripts by representative metabolic conditions. Our work uncovered a large number of novel transcripts including novel genes, many of which were regulated by physiologically relevant metabolic stimuli. Furthermore, our analysis of RNA transcripts at the levels of isoform expression, m6A modification and poly(A) tail length identified substantial dynamic changes of metabolic pathways and gene networks that are otherwise undetectable in conventional experimental design and short-read RNA-seq. We also presented evidence that the holistic landscape of metabolic responsive transcriptome revealed in our study was very unlikely to be attainable in animal or patient cohort-based studies. Thus, our work represents one of the first efforts to directly and comprehensively define metabolism-focused transcriptome dynamics of human liver and uncover a complex metabolic responsive landscape of the transcriptome that could help understand human liver physiology.

In order to study transcriptome dynamics of human liver under a physiological context, it is essential to establish an inclusive annotation of human liver transcriptome and study gene regulation in conjunction. To ensure an inclusive annotation, we subjected a humanized mouse model to diverse metabolic treatments and used DRS to analyze full length native RNAs to resolve key elements of the transcriptome: transcript isoforms, precise transcript-level dynamics, m6A and poly(A) tail length. Profiling transcripts under multiple conditions here was instrumental in capturing a wide range of transcripts and RNA modifications to study their regulation. Indeed, we found that over 30% of novel genes and transcripts identified in this annotation were only expressed under one specific condition which would otherwise be invisible if only samples of one condition were analyzed. When this work was being prepared, a report that analyzed Genotype-Tissue Expression (GTEx) samples using long-read sequencing was published uncovering a large number of novel transcripts ^48^, which is consistent with the conclusion of our work. Whereas this study collected samples from deceased individuals of random conditions, our annotation was regulation-oriented and was based on samples under well-controlled and representative metabolic conditions. Since our annotation was based on samples under metabolic conditions which we intended to study, all conditionally expressed transcripts and RNA modifications were already included. Thus it is no surprise that the regulated metabolic pathways identified by our transcript-level analyses were drastically different from those by conventional short-read RNA-seq, which mainly detects gene-level changes in transcripts that are documented in current reference annotations. Moreover, our study also revealed for the first time the dynamic changes in m6A modifications and poly(A) tail length of many key metabolic genes, which might constitute a new mechanism for the liver to regulate the related metabolic pathways at transcript isoform level. At a deeper level, our work was designed to reveal a holistic picture of human liver transcriptome dynamics in a physiological context, which has never been achieved before. In a cell, all gene expression networks are intertwined and strongly impact each other, and the intrinsic connections among gene networks can only be established under a condition where information of all gene expressions are available. But in a study involving individuals of different genetic backgrounds, such as a patient-based study, the holistic view would be lost once genes that were specifically regulated in certain patients were removed by cohort statistics. The humanized livers are clones of the same human liver thus addressing this critical issue. Although only humanized livers of a specific genetic background were used in this study and the expression of certain genes might be dependent on this background, many of the intrinsic connections among gene networks uncovered in this study could be universal and the information can be harnessed to uncover novel connections among metabolic gene networks.

Our data reinforced the limitations of studying human metabolism using mice, although they are no doubt a useful reference. It is usually difficult to directly compare gene regulation in human and mouse liver cells as it is impossible to place them under the identical in vivo condition. The humanized livers we utilized, however, have the advantage of being chimera in which human and mouse hepatocytes share the same circulation and tissue environment^16^. In addition to gene-level difference between the two species as previously reported^4^, our work also revealed much more profound differences at transcript level which would serve as a strong cautionary note to use mouse transcriptome dynamics to understand human metabolism. Of course, the humanized mouse model we were currently using has its limitations. The mice are immune-deficient and are not suitable to study the impact of immune cells on metabolism^16^. In addition, only hepatocytes were humanized in this model and some of the differences in human and mouse gene expressions revealed by this study were caused by the unequivalent cell populations. But as human liver is inaccessible to treatments, this model provides the closest possible approximation of human livers under representative metabolic conditions, at least at this stage, and this study represents one of the first direct analyses of metabolism-related transcriptome dynamics in human liver.

Our work could address a broad spectrum of readers. Basic scientists, particularly those who mainly use mouse as a model to study human metabolism, can use the extensive regulatory information of human genes we provided to infer the potential relevance of their genes of interest to human physiology and diseases. Physician scientists who study patient samples can potentially use the annotation and regulatory information of RNA transcripts to better pinpoint the changes in metabolic networks underlying a specific pathophysiology. Geneticists and epidemiologists can use the transcript-level details to further corroborate the importance of gene loci identified in their analyses. Finally, this work could serve as a framework and model for designing additional tools to gain direct knowledge of gene regulation in human organs under important pathophysiological conditions, which is the foundation to understanding diseases and developing effective therapies.

## Materials and Methods

### Animal experiments

All animal experiments were performed in accordance and with approval from the NHLBI Animal Care and Use Committee or the Animal Care Committee of the Central Institute for Experimental Animals (CIEA, Kanagawa, Japan). The TK-NOG mice (Cat. 12907, Taconic Biosciences) aged 8-10 weeks received intraperitoneal (i.p.) injection of Ganciclovir (NDC: 63323-315-10, APP Pharmaceuticals) at a dose of 20 mg/kg (4 µl/g) one week before surgery. Thawed Primary human hepatocytes were recovered in Cryopreserved Hepatocyte Recovery Medium (Cat. CM700, Thermo Fisher Scientific, Waltham, MA, USA), and re-suspended in ice-cold HBSS prior to surgery. Mice were anesthetized with Ketamine/Xylazine by i.p. injection and were injected with a 50 µl suspension containing 1×10^6^ primary human hepatocytes via the spleen. Eight weeks after the surgery, the humanized mice with serum human albumin levels above 0.5 mg/ml were utilized in experiments. For the dietary treatment, the humanized mice were allowed free access to food (AL, n=4 for donor1, n=3 for donor2) or subjected to twenty-four hours of food withdrawal (Fast, n=4 for donor1, n=3 for donor2). The livers were collected and stored in liquid nitrogen immediately upon sacrifice. For the transcription factor agonist treatments, the humanized mice were injected with DMSO (n=4), fenofibrate (n=4, 50mg/kg, Sigma-Aldrich, Cat. F6020), rosiglitazone (n=4,10mg/kg, Sigma-Aldrich, Cat. R2408) and GW4064 (n=4, 30mg/kg, Sigma-Aldrich, Cat. G5172) by i.p. injection. The livers were collected after 6 hours of fasting and stored in liquid nitrogen immediately upon sacrifice.

### RNA extraction

Approximately 40 mg of frozen liver tissue powders were gently homogenized 20-25 times using a Dounce homogenizer in 1.2 ml of Trizol reagent (Invitrogen, Cat. 15596018) on ice. The samples were transferred to new nuclease-free 2ml Eppendorf (EP) tubes and incubated on ice for 5 minutes. The lyses were centrifuged at 10,000 x g for 2 minutes (4℃), and 1 ml of each supernatant was transferred to new nuclease-free 1.5 ml EP tubes. Each sample received 200 μl per ml CHCl3 (Chloroform), and was then gently mixed. The mixtures were then incubated at room temperature for 5 minutes and centrifuged at 12,000 x g (4 °C) for 10 minutes. Five hundred microliters of the supernatants were transferred to new EP tubes. Five hundred microliters of isopropanol was added and then samples were incubated at room temperature for 15 minutes while being gently mixed. The samples were centrifuged at 12,000 x g (4 °C) for 10 minutes, and then the supernatants were removed. The RNA pellets were washed with 500 μl of 75% ethanol followed by centrifugation at 12,000 x g (4 °C) for 5 minutes. The supernatants were removed and the washing step was repeated once. The RNA pellets were stored at -80 °C in 75% ethanol or they can be processed to remove the supernatants and air-dried for 10 minutes. The air-dried RNA pellets were resuspended in 100 μl nuclease-free water and used immediately.

### Analysis of RNA-Seq data

RNAs that have been extracted by Trizol were purified using the MagMAX RNA extraction kit (Thermo Fisher Scientific, Cat.AM1830). Strand-specific sequencing libraries were constructed using the TruSeq Stranded Total RNA Prep kit (Illumina). DNA sequencings were performed at the NHLBI DNA Sequencing and Genomics Core. The original FASTQ read files were trimmed and cleaned using fastp/0.23.2, then quality analysis was performed using FastQC/0.11.8. A custom genome reference index was created by combining human (GRCh38.p13) and mouse (GRCm38.p6) genome sequences for alignment of humanized mouse chimeric liver RNA-seq data as previously described ^4,14^. The primary assemblies were obtained from the GENCODE database (https://www.gencodegenes.org/). The trimmed reads were aligned with default settings of HISAT2/2.2.1.0 and quantified using featureCounts (subread/2.0) with human annotation GENCODE v33 and mouse GENCODE vm24. The combined raw count files generated by the subread featureCounts tool were imported into the R package DESeq2/3.1.0, and were used for differential expression genes (DEGs) analysis. A cutoff of log2 (fold change) more than 0.5 and p value less than 0.05 was used for differential expression for all the liver samples.

### Analysis pipeline for public human RNA-Seq data

A dataset of RNA-Seq data used for cross-sectional human studies was retrieved from BioProject PRJNA523510, consisting of 57 liver samples from a Nonalcoholic fatty liver disease (NAFLD) population ^28^. This study included the liver biopsies from normal individuals (n=14), obese individuals (n=12), NAFLD patients (n=15) and nonalcoholic steatohepatitis (NASH) patients (n=16) which were determined from quantitative histomorphometry of liver fat, inflammation and fibrosis. The RNA-seq data were downloaded using sratoolkit/3.0.2 and decompressed into FASTQ read files. The FASTQ read files were trimmed and cleaned using fastp/0.23.2, then quality analysis was performed using FastQC/0.11.8. The reads were then aligned using HISAT2/2.2.1.0 with an index created using the GRCh38 genome and annotations generated from the Nanopore direct RNA-seq (DRS), respectively. RNA-Seq aligned reads sorting, quantification and DEGs analysis were conducted in the same way as humanized mouse RNA-Seq analysis. A log2 (fold change) cutoff greater than 0.5 and a p-value lower than 0.05 were used to identify DEGs.

### Isolation of nuclei from frozen liver tissues

The ATAC-Seq assay was performed as previously described with some modifications^49^. Briefly, around 40mg frozen humanized liver tissue powders from the same group were pooled together and homogenized gently using Dounce homogenizer with 20-25 strokes in ice-cold 2 ml homogenization buffer. The homogenized tissues were pre-sieved through a 100 µm filter, then centrifuged at 100 x g for 1 minute. Four hundred microliters of the supernatants were transferred to 2 ml round-bottomed Lo-Bind EP tubes. Then, 400 µl of 50% Iodixanol solutions (Sigma-Aldrich, Cat. D1556) were added and mixed to produce a final concentration of 25% Iodixanol solution mixture. Six hundred microliters of 29% Iodixanol solutions were added under the layer of 25% iodine acetone mixture. Then 600 µl of 35% Iodixanol solutions were added to the 29% iodine acetone mixture. The samples were centrifuged at 3,000 x g for 20 minutes using a swinging bucket centrifuge. Two hundred microliters of top layers were aspirated to new nuclease-free Lo-Bind EP tubes. The cell nuclei were counted and 50,000 of which were placed in 1 ml resuspension buffer containing 0.1% Tween-20. Then, the nuclei were centrifuged at 500 x g for 10 minutes at 4°C, and the supernatants were aspirated. Twenty-five microliters of 2x TD buffer, 2.5 µl of transposase (illumine, catalog number 20034210), 16.5 µl of PBS, 0.5 µl of 1% DAPI, 0.5 µl of 10% Tween-20, and 5 µl of H2O were added to the pellet and mixed thoroughly. They were then incubated for 30 minutes at 37°C mixing at 1,000 rpm/min. DNAs were purified using the MinElute Reaction Cleanup Kit (Qiagen, catalog number 28206) according to the manufacturer’s instructions. The purified DNAs were stored at -80 °C or processed to ATAC-Seq library preparation.

### ATAC-Seq library preparation, sequencing and analysis

The ATAC-Seq DNA library generations were processed using Kaestner lab’s pipeline^50^. The purified DNAs were amplified using TruSeq i7 index primers (Illumina) via PCR. The amplified DNAs were purified and size-selected using AMPure XP beads (Beckman Coulter) to remove primer dimers and fragments greater than 1,000 bp. The library quality and quantity were evaluated using Bioanalyzer and Qubit, and the final library was sequenced at NHLBI DNA Sequencing and Genomics Core.

The raw FASTQ reads were processed using the pipeline from Kundaje Laboratory [https://github.com/kundajelab/atac_dnase_pipelines], which included adapter trimming, alignment with genome using Bowtie2, and peak calling using MACS2. The genome used for Bowtie2 alignment was established using the same method as that used in the analysis of the humanized mouse RNA-Seq data. For peak calling analysis, filtered reads were used, which were retained after removal of unmapped reads and duplicates. Significant peaks were defined as those with a false discovery rate (FDR) Q value less than 0.01, which indicated the chromatin open regions.

### Polyadenylated RNA isolation and Nanopore direct RNA sequencing (DRS)

The 75 ug of fresh Trizol isolated RNAs were adjusted to a volume of 100 μL with nuclease-free water. The polyadenylated RNAs were selected using the Dynabeads™ mRNA Purification Kit (Invitrogen, Cat. 61006) following the manufacturer’s instructions. The resulting polyadenylated RNAs were resuspended with nuclease-free water. The quality and quantity of RNAs were assessed using the NanoDrop 2000 spectrophotometer (Thermo Fisher Scientific). The 500 ng of fresh isolated polyadenylated RNAs were used for Nanopore direct RNA sequencing (DRS) generally following the Oxford Nanopore Technologies (ONT) SQK-RNA002 kit protocol, including the optional reverse transcription step recommended by ONT. Libraries were loaded onto ONT R9.4 flow cells (Oxford Nanopore Technologies) and were sequenced using the GridION platform, with the standard MinKNOW/19.12.5 protocol script being used.

### Isoform-identification of DRS

The analysis flow was performed as previously described^38,51^. The ONT Guppy workflow (version 2.1.0) was used for basecalling the DRS data with the parameters “--flowcell FLO-MIN106 --kit SQK-RNA002 -x cuda:all --records_per_fastq 0” being employed^52^. The FASTQ files were aligned to the GRCh38 human genome reference and GRCm38 mouse genome reference respectively using minimap2/2.17, with the parameters “-ax splice -uf -k14” being employed. The FLAIR pipeline (https://github.com/BrooksLabUCSC/flair) with some modifications was employed to identify the isoform from the aligned reads. Firstly, reads with deletion length greater than 100 nt were removed. Secondly, only reads with 5’ ends that overlapped with chromatin open regions as indicated by ATAC-Seq information were considered. Next, the splice-site boundaries of DRS reads were corrected by matching them with corresponding short read splice junctions and only the junctions that were supported by at least three uniquely mapped short reads were considered valid and included ^51^. Finally, FLAIR *collapse* with the default settings was utilized to generate DRS isoform annotations for both human and mouse, including splicing sites and sequence information.

### Quantification, differential expression and alternative splicing analyses of DRS

The Nanopore reads were quantified to DRS annotation using FLAIR *quantify* by default parameters. Only alignments with quality scores of 1 or greater were counted. The quantified isoform counts were analyzed using FLAIR *diffExp* for characterization of differentially expressed genes (DEGs) and transcripts (DETs), as well as FLAIR *diffSplice* for analysis of alternative splicing (AS) events. The default parameters were used for all analyses. To find the DEGs and DETs, a threshold of log2 (fold change) larger than 0.5 and a p value less than 0.05 were used. The events having a p value of less than 0.05 were considered to have significantly altered AS.

### m6A Modification Detection and Analysis

The RNA m6A modifications of DRS were examined using the MINES (https://github.com/YeoLab/MINES)^43^. Briefly, the DRS reads and modification values were aligned by Tombo (https://github.com/nanoporetech/tombo) with either a genomic or a cDNA (transcriptomic) reference using the default *re-squiggle* and de novo detection settings, respectively. Genomic references (GRCh38/hg38 for human and GRCm38/mm10 for mouse) were downloaded from GENCODE, and cDNA references came from DRS isoform identification step. A new set of regions was created by extending 10 bp on either side of the “A” in the DRACH motifs, which allowed for the identification of all areas in the reference that had a DRACH motif. These areas were then further filtered to have a minimum of five reads of coverage. In the same treatment group, modification events that occurred in less than half of the samples were excluded. The distance measure and metagene analysis of the modification sites were performed by MetaPlotR (https://github.com/olarerin/metaPlotR) using default setting^53^. DESeq2/3.1.0 was used to examine the differential modification events with its default settings. The modification event counts within the same genes were combined and sent for analysis for gene level comparison. A log2 (fold change) cutoff greater than 0.5 and a p-value lower than 0.05 were used to identify significantly changed modifications.

### Poly(A) tail length analysis of DRS

The pipeline-poly(A)-ng (https://github.com/nanoporetech/pipeline-poly(A)-ng) was used to call the estimated lengths of poly(A) tails from Nanopore DRS. The transcript sequence data extracted from the DRS isoform identification step was used as transcriptome reference. Reads with a mapping quality of less than 5 were excluded. The other options were set to be the defaults.

Pipeline-poly(A)-diff (https://github.com/nanoporetech/pipeline-poly(A)-diff) was used to compare the changes in transcript poly(A) tail length between treatments. The parameters were all set to their default values. For gene level comparison, the transcripts within the same genes were pooled and sent for Student’s t-test analysis. For testing significant shifts in transcript poly(A) tail lengths, a cutoff of length changes greater than 10 nt and p value less than 0.05 was applied.

### Protein Structure Prediction and Visualization

The putative peptide sequences for the GSTM1 transcripts were predicted by CPC2 (http://cpc2.gao-lab.org/index.php)54, using the default parameters. AlphaFold/2.1.2 algorithm was employed to protein structures using the default parameters as “run_singularity-- model_preset=monomer_casp14--fasta_paths=data.fasta--max_template_date=2020-05-14 -- output_dir=$PWD”, (https://github.com/deepmind/alphafold)55. The top ranked structure models were sent to PyMOL/2.3 for the visualization and comparison.

### Novel Transcripts and Novel Gene Identification

The reference annotations were downloaded from GENCODE database (https://www.gencodegenes.org/,GENCODE v33 for human and GENCODE vM24 for mouse). The DRS and public reference annotation were compared using the GffCompare (https://github.com/gpertea/gffcompare) using default settings^56^. The output GTF file’s attribute value from “class code” was used to categorize the transcripts/isoforms. The transcripts that had a class code of “=” were considered to be matched transcripts since their intron chains matched the reference annotation exactly. Novel/different transcripts were defined as the ones that were assigned with the “class code” of either “m, j, o, x, i, y, or u” and that were found to have different isoform structures from the reference annotation. Transcripts that were labeled as “others” were those that were assigned with the “class code” of either “c, k, n, e, s, p or r” and that had been found to have partially matched splicing sites or no actual overlap with the reference annotation.

Additionally, the transcripts from novel transcripts with the “class code” of “u” were categorized as new genes since they showed no overlap with any genes in the reference annotation.

### Protein sequences comparison of novel transcripts

The novel transcripts that showed the highest levels of expression within their respective genes were identified as the dominant ones in both human and mouse. The dominant novel transcript sequences were sent to ORFfinder/0.4.3 with the default settings to search for open reading frames (ORFs), and those with lengths less than 50 amino acids were excluded. To compare the ORF protein sequences with reference protein databases, the longest ORFs for each transcript were sent to blastp /2.13.0 with default settings. The publicly available reference protein-coding transcript translation sequences were retrieved from GENCODE (https://www.gencodegenes.org/, GENCODE v33 for human and GENCODE vM24 for mouse). The ORFs that had an alignment coverage percentage of greater than 90% and an E-value of less than 0.01 were considered to be the matched ORFs.

### Coding Potential Analysis

The sequences of the novel transcripts from lncRNA regions were sent to five coding potential prediction tools with different intrinsic sequence-related features (composition, structural properties and motifs) and divergent filtering steps respectively. The potential coding transcripts were defined as following cutoff in each algorithm: CPC2 (score > 0.5, http://cpc2.gao-lab.org/index.php)^54^, CPAT (score > 0.36, http://lilab.research.bcm.edu/cpat/index.php) ^57^, CNCI (score > 0, https://github.com/www-bioinfo-org/CNCI)58, FEElnc (default parameters, https://github.com/tderrien/FEELnc)^59^ and PLEK (score > 0, https://sourceforge.net/projects/plek/files/)^60^.

### Novel Gene Cloning, Gel Analysis and Sequencing

The Gateway BP cloning system (Thermo Fisher Scientific) was employed for gene cloning according to the manufacturer’s instructions. The attB PCR primers were designed by adding the attB sequence to the 5’-terminal of the forward (GGGGACAAGTTTGTACAAAAAAGCAGGCT) and reverse (GGGGACCACTTTGTACAAGAAAGCTGGGT) primers, respectively. (Table 4). Humanized liver cDNA samples were used to clone the novel gene DNA fragments using PCR-amplifying. Then, the attB-flanked PCR products with the correct sizes, as determined by gel electrophoresis, were purified and cloned into pDONR221 vectors using the Gateway BP reaction system. Sequencing by Synthesis (SBS) sequencing was used to identify the novel gene sequences following transformation and plasmid extraction. The similarity between the gene sequences obtained from SBS sequencing and the Nanopore annotation was determined using BLAST (https://blast.ncbi.nlm.nih.gov/Blast.cgi).

### Primary human hepatocytes and knockdown assays

The primary human hepatocytes (PHH, Lonza) were thawed and plated following the manufacturer’s introduction. After 5 hours of plating, they were switched to five chemicals’ (5C) long-term functional maintenance medium^61^. 24 hours after plating, human primary hepatocytes were transfected with siRNA negative control (siNC) or three different siRNAs (siRNA1: GCAGACAGAUUUCCUAGAAAU; siRNA2: GGUGUGUUCAAAGAGCAAAGA; siRNA3: GGCAAAUGAAGGAUUGAAACG) at a dose of 10 nM for 48 hours using Lipofectamine™ RNAiMAX (Thermo Fisher Scientific, Cat. 13778150). RNAs were then extracted from these cells using MagMAX RNA Extraction Kit (Thermo Fisher Scientific, Cat. AM1830) and sent for transcriptome analysis using RNA-Seq. Each group included three independent duplicates.

### Gene Set Enrichment Analysis (GSEA)

Gene Set Enrichment Analysis (GSEA) was performed to identify gene sets that were differentially expressed between the knockdown (siRNA1, siRNA2, and siRNA3) and control groups using the RNA-Seq data of hNovel3. The GSEA analysis was conducted using the software provided by the Broad Institute (http://www.gsea-msigdb.org/gsea/index.jsp). The RNA-Seq was preprocessed as descripted and normalized by DESeq2/3.1.0. The Sterol Biosynthetic Process pathway gene set was obtained from the MSigDB database (https://www.gsea-msigdb.org/gsea/msigdb/). The normalized enrichment score (NES) and the False Discovery Rate (FDR) q value for the gene set were calculated. The enrichment plots were then generated to visualize the results.

### Gene Ontology (GO), Kyoto Encyclopedia of Genes and Genomes (KEGG) and Gene Set Variation Analysis (GSVA) Pathway Enrichment Analyses

The GO and KEGG pathway enrichment analyses were performed and visualized by R package clusterProfiler/3.18.0 using the default setting (https://github.com/YuLab-SMU/clusterProfiler). Short read and Nanopore DRS gene expression matrixes were sent to GSVA/1.38.0 (https://github.com/rcastelo/GSVA) to estimate the enrichment signatures of pathways for each sample^62^. The limma/3.46.0^63^ R package was used to investigate the differentially enriched pathways based on GSVA scores between Fast and AL. The GSVA enrichment scores for the differently enriched pathways were visualized by R package pheatmap/1.0.12 (adjust p value 0.05) after z-score normalization.

### Principal component analysis (PCA)

The DESeq2/3.1.0 package in R was utilized to do the principal component analysis (PCA) on expression data, alternative splicing events, and m6A modification coverages. The “rlog” function in DESeq2 was utilized to regularize the data’s log transformation, which improved its suitability for PCA by reducing the data’s variance. The “plotPCA” function in DESeq2 was then utilized to conduct PCA on the transformed data. The first two principal components were plotted and samples were colored according to a grouping variable.

### Homology gene analysis between human and mouse

The list of homologous genes for humans and mice was downloaded from the MGI database (https://www.informatics.jax.org/homology.shtml). For RNA m6A modifications and poly(A) tail length analyses, the transcript level data were collapsed to gene levels. Only the genes that significantly changed in both mouse and human were considered for consistently or oppositely regulated genes. Among which, the homologous genes that changed in the same direction were defined as consistent, while those that changed in opposite directions were defined as opposite. Homology genes that only changed significantly in one species or showed no changes in both species were defined as inconsistent.

### Statistics

The two-tailed unpaired Student’s t-test was used for comparisons between two groups shown in Fig. 1E, Fig. 2B, Fig. 3F, and Fig. 4E and Fig. S1J, 3D-F, and 4B. For comparisons of Poly(A) tail length variance, Kruskal–Wallis one-way analysis was used in Fig. 3E and Fig. S3G. A p value of less than 0.05 was considered significant.

## Study approval

All animal experiments were performed in accordance with and with approval from the NHLBI Animal Care and Use Committee or the Animal Care Committee of the CIEA, Kawasaki, Japan. All human-related data sets were downloaded from public domains.

## Data availability

The source data underlying Figures, Extended Data Figures and Tables are provided as a Source Data file. The raw sequencing data can be accessed at GEO through the SuperSeries dataset GSE224281, which consists of multiple SubSeries, including GSE130525 and GSE224279 for RNA-Seq data of donor 1, GSE126587 for RNA-Seq data of donor 2, GSE224277 for ATAC-Seq data, and GSE224278 for nanopore direct RNA sequencing data. All data is available from the corresponding author upon reasonable request.

## Author Contributions

CJ, PL and HC designed the workflow. CJ carried out the Nanopore direct RNA sequencing and bioinformatics analyses. CJ and YM processed humanized liver samples and analyzed the function of a novel gene. NY, MN, YO and HS prepared and treated humanized mice. CJ, PL and HC wrote the manuscript. PL and HC conceived and supervised the study.

## Acknowledgments

We thank Yan Luo, Poching Liu and Yuesheng Li (NHLBI DNA Sequencing and Genomics Core) for RNA-seq analysis. This study was funded by NHLBI Division of Intramural Research funds to HC (1ZIAHL006103, 1ZIAHL006159).

## Supplementary Figure Legend

**Fig. S1 A de novo annotation of human liver transcriptomes reflecting pathophysiologically relevant metabolic responses**

(A) Length distribution of the human and mouse DRS transcripts. (B) Heatmaps of common DEGs’ relative expression levels in short read RNA-seq and DRS under the Fast treatments in human and mouse, respectively. The sequencing types and treatments are labeled at the top of the heatmap. The expression levels were normalized by Z-score method. (C) Central network plot of top pathways (GO BP) and hub genes for the DRS DEGs of Fast treatment in human and mouse, respectively. (D) Heatmaps of the relative pathway enrichment scores that were differentially changed between Fast and AL treatments in human and mouse, respectively (|log2 (fold change)| >0.5 and p value <0.05). The pathway enrichment scores for short read and Nanopore DRS were estimated by Gene Set Variation Analysis (GSVA) and were further normalized by Z-score method. (E) Pie plots of the composition of transcripts that were specifically expressed in response to dietary intervention (left) and transcription factor activation treatments (right). Transcripts were characterized as specific (only expressed in one condition), common (expressed in all conditions), or multi-condition (expressed in two or more but not all conditions). (F) Comparison of Nanopore DRS and reference annotations for the different/novel transcripts that were dominantly expressed in human and mouse, respectively. (G) Top: Heatmap of the coding protein potential results for novel/different transcripts from lncRNA regions. Five coding potential analysis algorithms (CPC2, PLEK, CPAT, CNCI, FEELnc) were used to examine the novel/different transcript sequences, and the positive hints for coding transcripts were colored as yellow. Bottom: UpSet plot of intersection between the estimated coding transcripts generated from different algorithms. (H) Top: Isoform schematics of DRS novel transcript (495df9d1, red) and reference transcript (ENST00000652241, dark green) in lncRNA AC005538.3 genome region. The raw reads from short-reads RNA-seq and Nanopore DRS in humanized mice livers were labeled as light blue. Bottom: The table of the results of coding potential analysis using CPC2 for the new transcript (495df9d1) and the reference transcript (ENST00000652241). (I) Heatmap of relative expression levels of novel gene DEGs in the samples of Non-alcoholic fatty liver disease database (NAFLD, PRJNA523510). (J) Expression levels of hNovel3 in the humanized mouse liver in DMSO control and FXR agonist treatment samples. Data represent mean ± SEM, **p < 0.01, two-tailed unpaired Student’s t-test. (K) Isoform schematics of hNovel 3 from DRS (dark brown) and reference (no transcript). The reads peaks of ATAC-Seq, H3K27Ac ChIP-seq and short-reads RNA-seq from humanized mice livers in hNovel3 genome region were labeled as brown. The signal peaks of short-read RNA-seq in primary human hepatocytes from the control group were labeled blue, and those from the different hNovel knockdown groups were labeled as black.

**Fig. S2 Human liver transcriptome dynamics in response to representative metabolic treatments**

(A) Network plot of the top enriched pathways (GO BP) and hub genes for the commonly regulated DETs in the treatments of Fast and FXR, PPARα, and PPAR γ agonists. (B) Heatmap of the expression levels of DETs that are commonly changed across all treatments. The summaries of the different transcript types compared to the reference annotation and their overlap with DEGs from short read RNA-seq data were displayed on the right. (C) Heatmap of the expression levels of transcripts specifically changed in one condition. The types of transcription factor agonists were labeled on the left. (D) Bar plot of the top enriched pathways for the conditionally changed DETs (PPARα in purple, PPAR γ in dark blue, and FXR in gray). (E) PCA plot of human splicing events across samples treated with AL or Fast. (F) APOA1 splice-site usage (left) and full isoforms (right) for the proximal chr11: 116,837,604 (purple) and distal chr11: 116,837,558 (dark blue) sites from 5′ to 3′. The alternative acceptor event is boxed in grey in the isoform schematic.

**Fig. S3 Dynamics of m6A modification and Poly(A) tail length of RNA transcripts in humanized livers**

(A) Metagene analysis of human m6A modification sites with different treatments. (B) PCA plot of human m6A modification events localized in CDS regions with different treatments. (C) Left top: Distribution of human m6A modification sites for AL (in blue) and donor2 (in red), with different modifications between them labeled as green. Right bottom: Venn diagram of the overlapping m6A modification locations between the AL and Fast. (D) Box plot of the Poly(A) tail length for different types of human genes, including protein-coding genes, lncRNAs and other genes (pseudogenes and other non-coding genes). ****p < 0.0001, two-tailed unpaired Student’s t-test. (E) Box plot of the Poly(A) tail length for transcripts with different expression levels, with low expression in red, middle expression in blue, and high expression in green. The transcripts were sorted by expression level, with the top 25% considered high expression, the bottom 25% considered low expression, and the middle 50% considered middle expression. ****p < 0.0001, two-tailed unpaired Student’s t-test. (F) Violin plot of the Poly(A) tail length of transcripts with and without intron-retained splicing modifications. ****p < 0.0001, two-tailed unpaired Student’s t-test. (G) Kruskal-Wallis test for poly(A) tail length variance of gene isoforms in Fast and AL treatments. The dashed line labeled the significant bar (p value 0.05).

**Fig. S4 Divergent transcriptome architecture and dynamics between human and mouse livers**

(A) Bar plot of the top enriched pathways (GO BP) for the homologous genes that had consistently or oppositely regulated Poly(A) tail length between human and mouse in different transcription factor agonist treatments. (B) Boxplot of poly(A) tail lengths of homologous genes from the lipid catabolic process pathway that showed opposite regulation patterns between human and mouse in AL and Fast treatments. * p<0.05, ** p<0.01, ***p < 0.001,****p < 0.0001, two-tailed unpaired Student’s t-test.

**Fig. S5 Divergent transcriptome architecture and dynamics between individuals of different genetic backgrounds**

(A, C) Venn diagram of the overlapping genes showing the significantly changed m6A modifications (A) and poly(A) tails (C) in response to Fast treatment between donor1 and donor2. (B, D) Top enriched pathways of genes for common, donor1-specific and donor2-specific changed m6A modifications (B) and poly(A) tails (D). The donor1 specifically enriched pathways were labeled as red, the donor2 specifically enriched pathways were labeled as blue and commonly changed pathways were labeled as black.

