## Supplementary Figures for "Single transcript-level metabolic responsive landscape of human liver transcriptome"

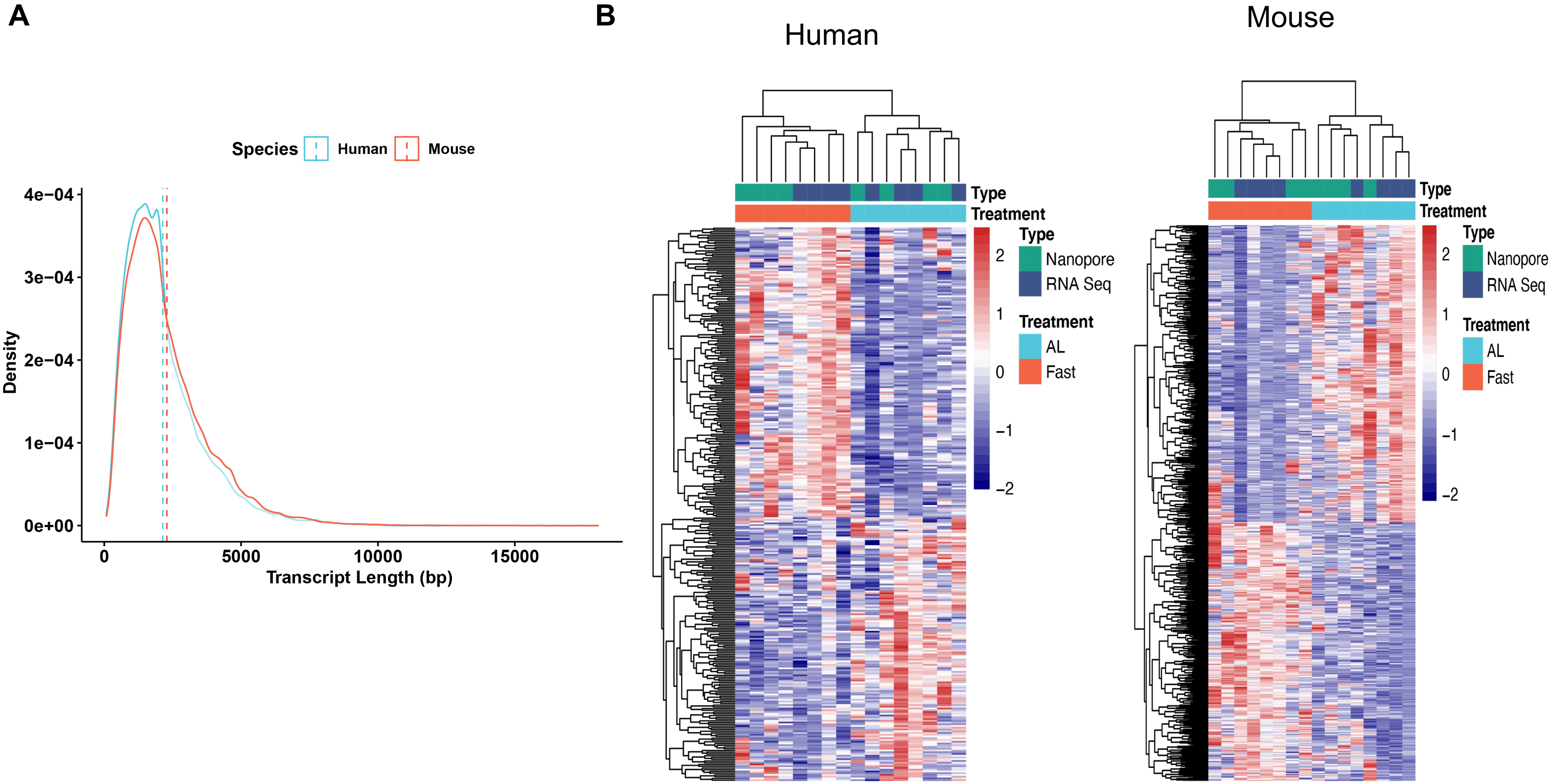
A
Species
Human
Mouse
4e−04
3e−04
2e−04
1e−04
0e+00
Density
0
5000
10000
15000
Transcript Length (bp)
B
Human
Mouse
Type
Treatment
Type
Nanopore
RNA Seq
Treatment
AL
Fast
2
1
0
−1
−2

Fig. S1

C

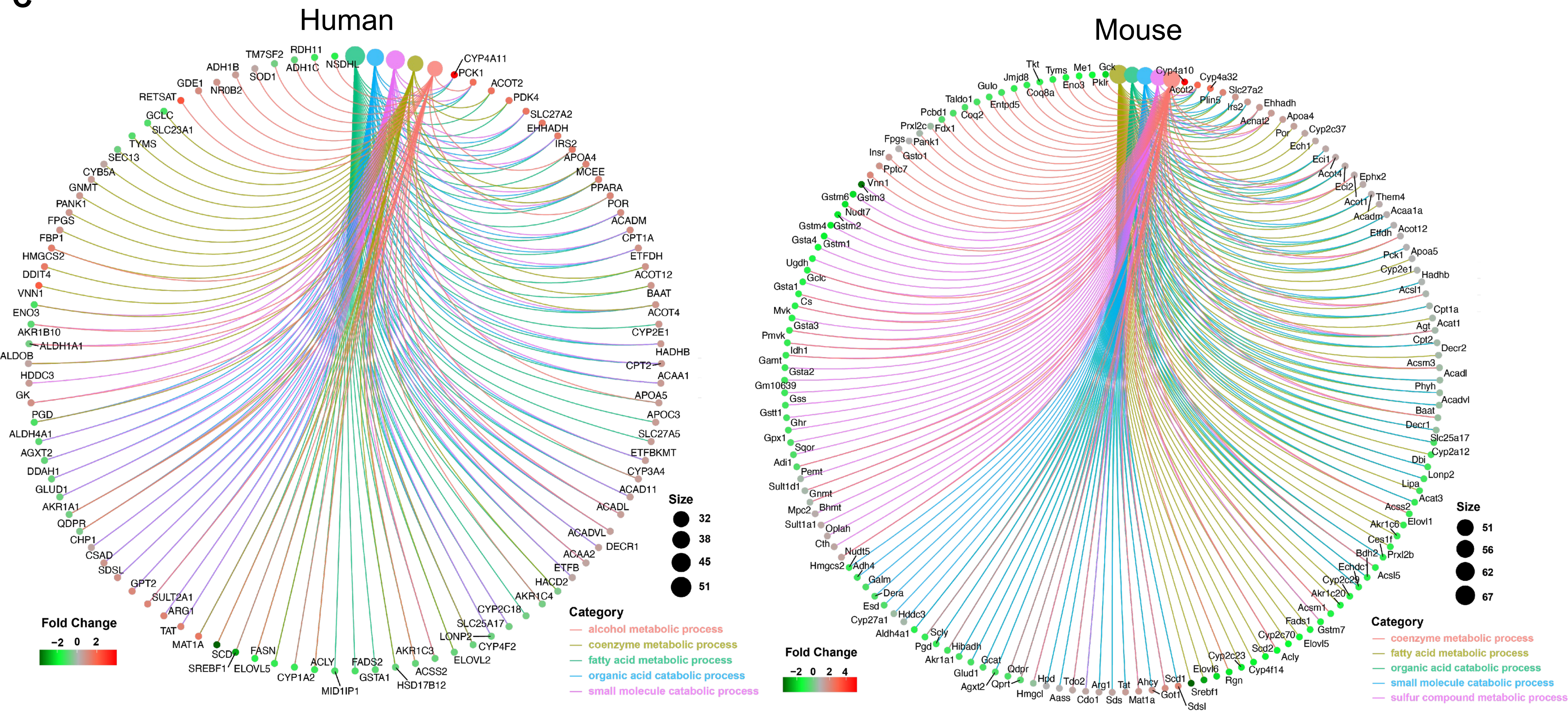

Human
Mouse
TM7SF2
RDH11
ADH1B
ADH1C
NSDHL
SOD1
GDE1
NR0B2
RETSAT
GCLC
SLC23A1
TYMS
SEC13
CYB5A
GNMT
PANK1
FPGS
FBP1
HMGCS2
DDIT4
VNN1
ENO3
AKR1B10
ALDH1A1
ALDOB
HDDC3
GK
PGD
ALDH4A1
AGXT2
DDAH1
GLUD1
AKR1A1
QDPR
CHP1
CSAD
SDSL
GPT2
SULT2A1
ARG1
TAT
MAT1A
SCD
SREBF1
ELOVL5
FASN
CYP1A2
ACLY
MID1IP1
FADS2
GSTA1
HSD17B12
AKR1C3
ACSS2
ELOVL2
LONP2
CYP4F2
SLC25A17
CYP2C18
AKR1C4
HACD2
ETFB
ACAA2
DECR1
ACADVL
ACADL
ACAD11
CYP3A4
ETFBKMT
SLC27A5
APOC3
APOA5
ACAA1
CPT2
HADHB
CYP2E1
ACOT4
BAAT
ACOT12
ETFDH
CPT1A
ACADM
POR
PPARA
MCEE
APOA4
IRS2
EHHADH
SLC27A2
PDK4
ACOT2
PCK1
CYP4A11
Size
32
38
45
51
Fold Change
-2 0 2
Category
alcohol metabolic process
coenzyme metabolic process
fatty acid metabolic process
organic acid catabolic process
small molecule catabolic process
Tkt
Jmjd8
Tyms
Me1
Gck
Gulo
Eno3
Pklr
Coq8a
Taldo1
Entpd5
Pcbd1
Coq2
Prxl2c
Fdx1
Fpgs
Pank1
Insr
Gsto1
Pptc7
Vnn1
Gstm6
Gstm3
Nudt7
Gstm4
Gstm2
Gsta4
Gstm1
Ugdh
Gclc
Gsta1
Cs
Mvk
Gsta3
Pmvk
Idh1
Gamt
Gsta2
Gm10639
Gss
Gstt1
Ghr
Gpx1
Sqor
Adi1
Pemt
Sult1d1
Gnmt
Mpc2
Bhmt
Sult1a1
Oplah
Cth
Nudt5
Hmgcs2
Adh4
Galm
Dera
Esd
Cyp27a1
Hddc3
Aldh4a1
Scly
Pgd
Hibadh
Akr1a1
Gcat
Glud1
Qdpr
Agxt2
Qprt
Hpd
Hmgcl
Tdo2
Aass
Cdo1
Arg1
Sds
Tat
Mat1a
Ahcy
Scd1
Got1
Sdsl
Srebf1
Elovl6
Rgn
Cyp2c23
Cyp4f14
Scd2
Acly
Cyp2c70
Elovl5
Fads1
Gstm7
Acsm1
Akr1c20
Cyp2c29
Echdc1
Acsl5
Bdh2
Prxl2b
Ces1f
Akr1c6
Elovl1
Acss2
Acat3
Lipa
Lonp2
Dbi
Cyp2a12
Slc25a17
Decr1
Baat
Acadvl
Phyh
Acadl
Acsm3
Decr2
Cpt2
Agt
Acat1
Cpt1a
Acsl1
Hadhb
Cyp2e1
Pck1
Apoa5
Etfdh
Acot12
Acadm
Acaa1a
Acot1
Them4
Eci2
Ephx2
Acot4
Eci1
Ech1
Por
Cyp2c37
Apoa4
Acnat2
Ehhadh
Irs2
Plin5
Slc27a2
Acot2
Cyp4a32
Cyp4a10
Size
51
56
62
67
Fold Change
-2 0 2 4
Category
coenzyme metabolic process
fatty acid metabolic process
organic acid catabolic process
small molecule catabolic process
sulfur compound metabolic process


Fig. S1

D

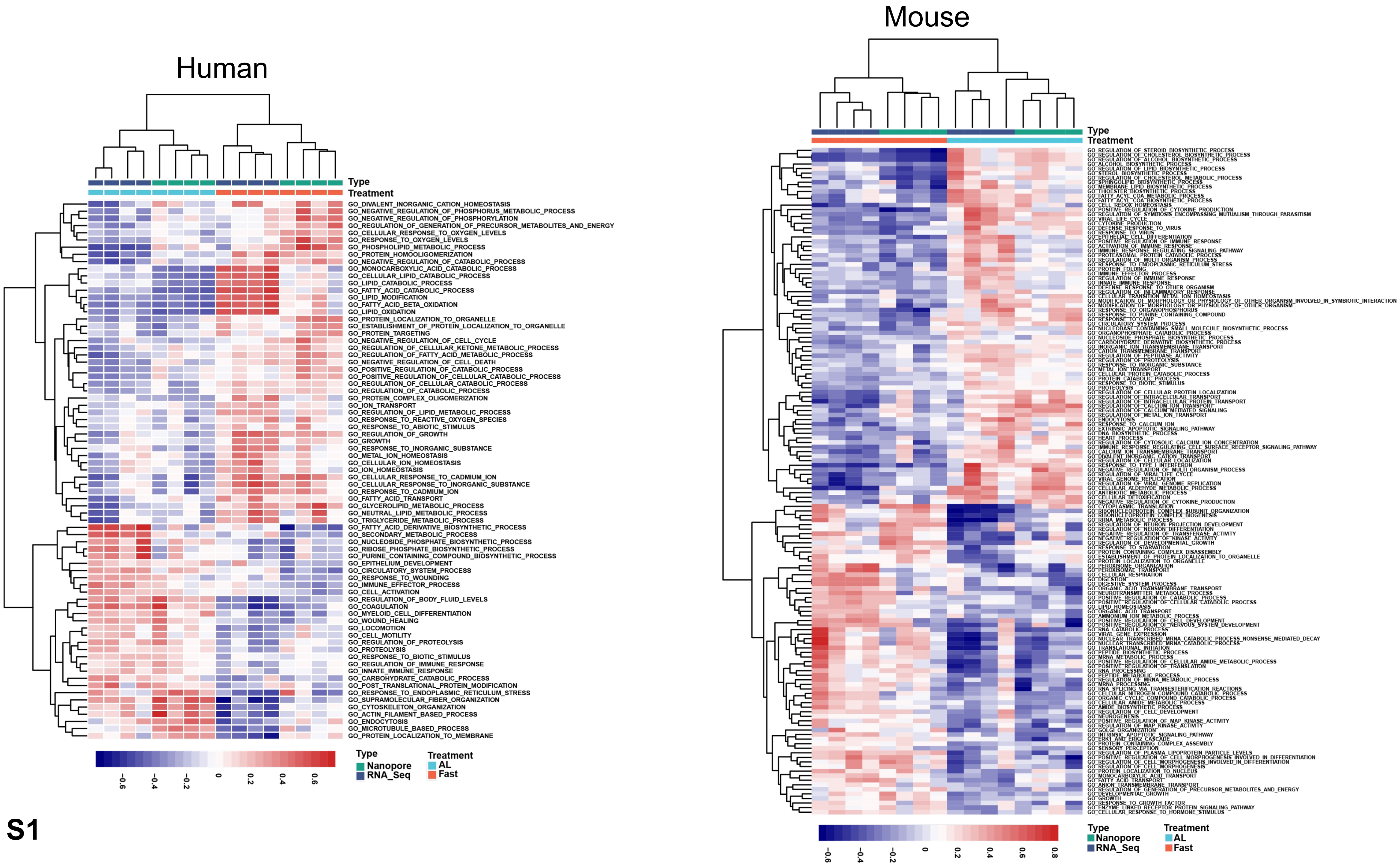

Human
Type
Treatment
GO_DIVALENT_INORGANIC_CATION_HOMEOSTASIS
GO_NEGATIVE_REGULATION_OF_PHOSPHORUS_METABOLIC_PROCESS
GO_NEGATIVE_REGULATION_OF_PHOSPHORYLATION
GO_REGULATION_OF_GENERATION_OF_PRECURSOR_METABOLITES_AND_ENERGY
GO_CELLULAR_RESPONSE_TO_OXYGEN_LEVELS
GO_RESPONSE_TO_OXYGEN_LEVELS
GO_PHOSPHOLIPID_METABOLIC_PROCESS
GO_PROTEIN_HOMOOLIGOMERIZATION
GO_NEGATIVE_REGULATION_OF_CATABOLIC_PROCESS
GO_MONOCARBOXYLIC_ACID_CATABOLIC_PROCESS
GO_CELLULAR_LIPID_CATABOLIC_PROCESS
GO_LIPID_CATABOLIC_PROCESS
GO_FATTY_ACID_CATABOLIC_PROCESS
GO_LIPID_MODIFICATION
GO_FATTY_ACID_BETA_OXIDATION
GO_LIPID_OXIDATION
GO_PROTEIN_LOCALIZATION_TO_ORGANELLE
GO_ESTABLISHMENT_OF_PROTEIN_LOCALIZATION_TO_ORGANELLE
GO_PROTEIN_TARGETING
GO_NEGATIVE_REGULATION_OF_CELL_CYCLE
GO_REGULATION_OF_CELLULAR_KETONE_METABOLIC_PROCESS
GO_REGULATION_OF_FATTY_ACID_METABOLIC_PROCESS
GO_NEGATIVE_REGULATION_OF_CELL_DEATH
GO_POSITIVE_REGULATION_OF_CATABOLIC_PROCESS
GO_POSITIVE_REGULATION_OF_CELLULAR_CATABOLIC_PROCESS
GO_REGULATION_OF_CELLULAR_CATABOLIC_PROCESS
GO_REGULATION_OF_CATABOLIC_PROCESS
GO_PROTEIN_COMPLEX_OLIGOMERIZATION
GO_ION_TRANSPORT
GO_REGULATION_OF_LIPID_METABOLIC_PROCESS
GO_RESPONSE_TO_REACTIVE_OXYGEN_SPECIES
GO_RESPONSE_TO_ABIOTIC_STIMULUS
GO_REGULATION_OF_GROWTH
GO_GROWTH
GO_RESPONSE_TO_INORGANIC_SUBSTANCE
GO_METAL_ION_HOMEOSTASIS
GO_CELLULAR_ION_HOMEOSTASIS
GO_ION_HOMEOSTASIS
GO_CELLULAR_RESPONSE_TO_CADMIUM_ION
GO_CELLULAR_RESPONSE_TO_INORGANIC_SUBSTANCE
GO_RESPONSE_TO_CADMIUM_ION
GO_FATTY_ACID_TRANSPORT
GO_GLYCEROLIPID_METABOLIC_PROCESS
GO_NEUTRAL_LIPID_METABOLIC_PROCESS
GO_TRIGLYCERIDE_METABOLIC_PROCESS
GO_FATTY_ACID_DERIVATIVE_BIOSYNTHETIC_PROCESS
GO_SECONDARY_METABOLIC_PROCESS
GO_NUCLEOSIDE_PHOSPHATE_BIOSYNTHETIC_PROCESS
GO_RIBOSE_PHOSPHATE_BIOSYNTHETIC_PROCESS
GO_PURINE_CONTAINING_COMPOUND_BIOSYNTHETIC_PROCESS
GO_EPITHELIUM_DEVELOPMENT
GO_CIRCULATORY_SYSTEM_PROCESS
GO_RESPONSE_TO_WOUNDING
GO_IMMUNE_EFFECTOR_PROCESS
GO_CELL_ACTIVATION
GO_REGULATION_OF_BODY_FLUID_LEVELS
GO_COAGULATION
GO_MYELOID_CELL_DIFFERENTIATION
GO_WOUND_HEALING
GO_LOCOMOTION
GO_CELL_MOTILITY
GO_REGULATION_OF_PROTEOLYSIS
GO_PROTEOLYSIS
GO_RESPONSE_TO_BIOTIC_STIMULUS
GO_REGULATION_OF_IMMUNE_RESPONSE
GO_INNATE_IMMUNE_RESPONSE
GO_CARBOHYDRATE_CATABOLIC_PROCESS
GO_POST_TRANSLATIONAL_PROTEIN_MODIFICATION
GO_RESPONSE_TO_ENDOPLASMIC_RETICULUM_STRESS
GO_SUPRAMOLECULAR_FIBER_ORGANIZATION
GO_CYTOSKELETON_ORGANIZATION
GO_ACTIN_FILAMENT_BASED_PROCESS
GO_ENDOCYTOSIS
GO_MICROTUBULE_BASED_PROCESS
GO_PROTEIN_LOCALIZATION_TO_MEMBRANE
-0.6
-0.4
-0.2
0
0.2
0.4
0.6
Type
Nanopore
RNA_Seq
Treatment
AL
Fast
Mouse
Type
Treatment
-0.6
-0.4
-0.2
0
0.2
0.4
0.6
0.8
Type
Nanopore
RNA_Seq
Treatment
AL
Fast


Fig. S1

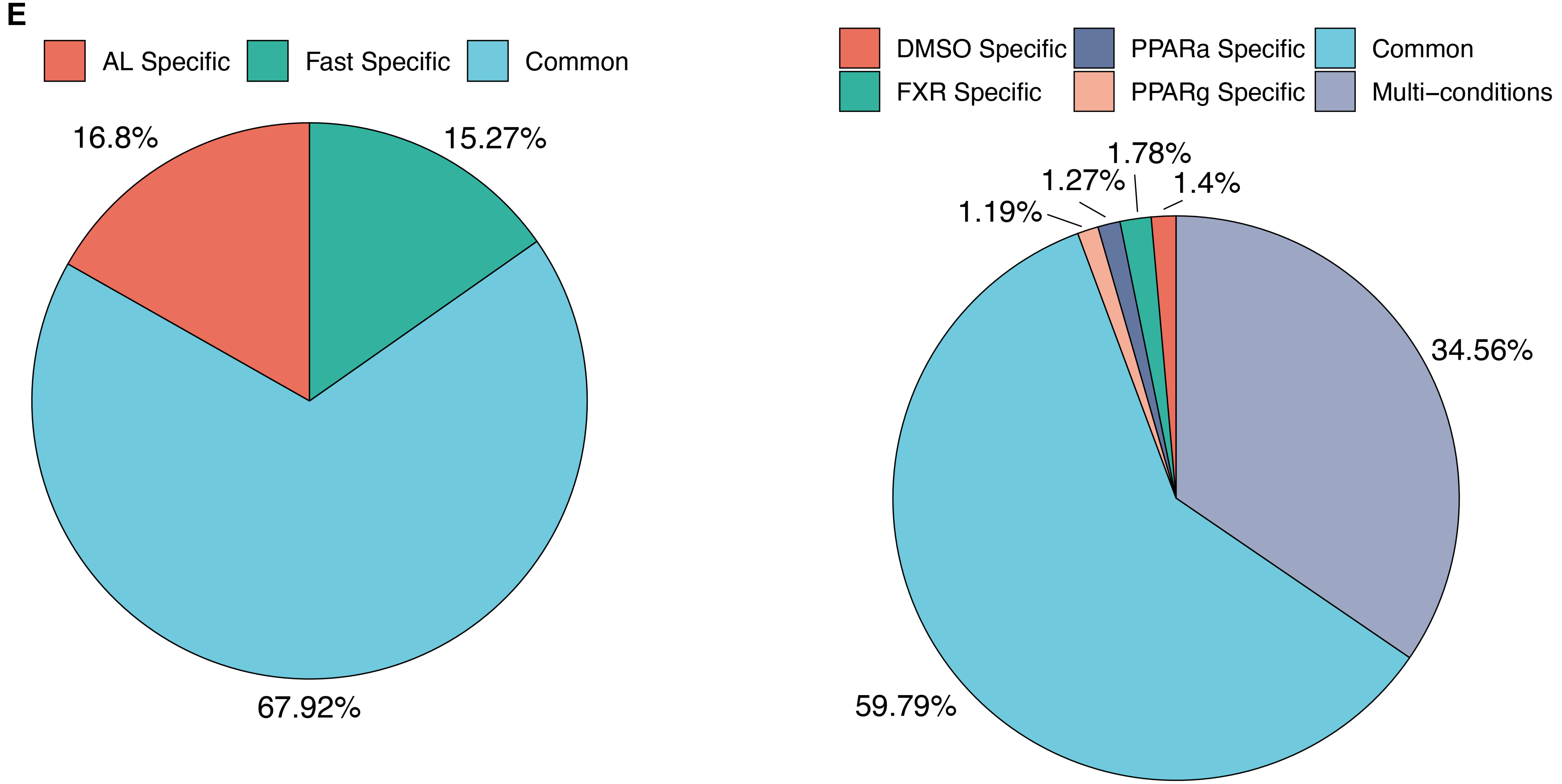
E
AL Specific
Fast Specific
Common
16.8%
15.27%
67.92%
DMSO Specific
PPARa Specific
Common
FXR Specific
PPARg Specific
Multi−conditions
1.78%
1.27%
1.4%
1.19%
34.56%
59.79%

Fig. S1

F

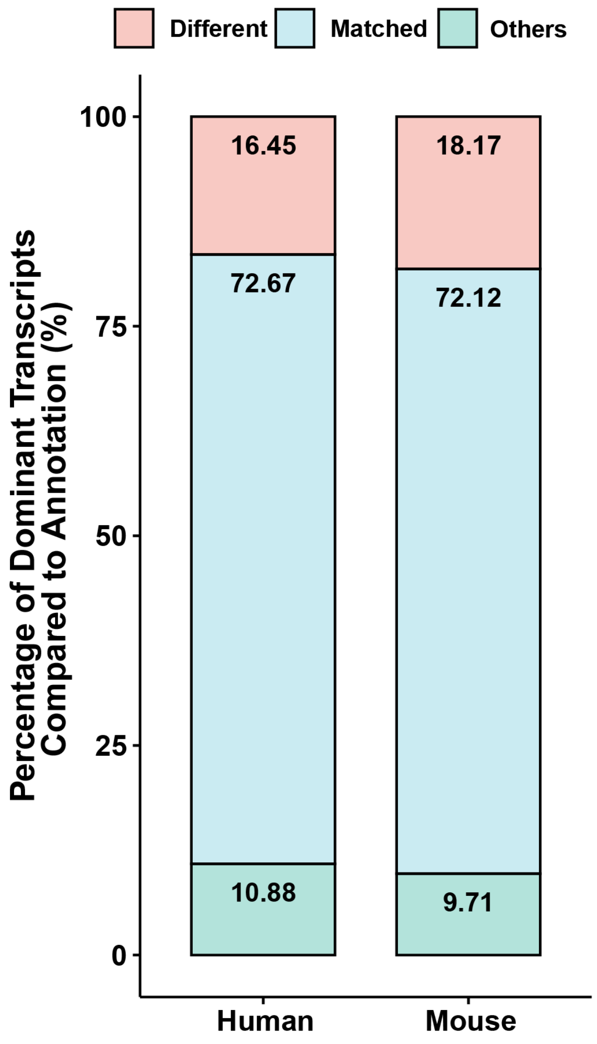

Different
Matched
Others
Percentage of Dominant Transcripts Compared to Annotation (%)
100
75
50
25
0
16.45
72.67
10.88
18.17
72.12
9.71
Human
Mouse


G

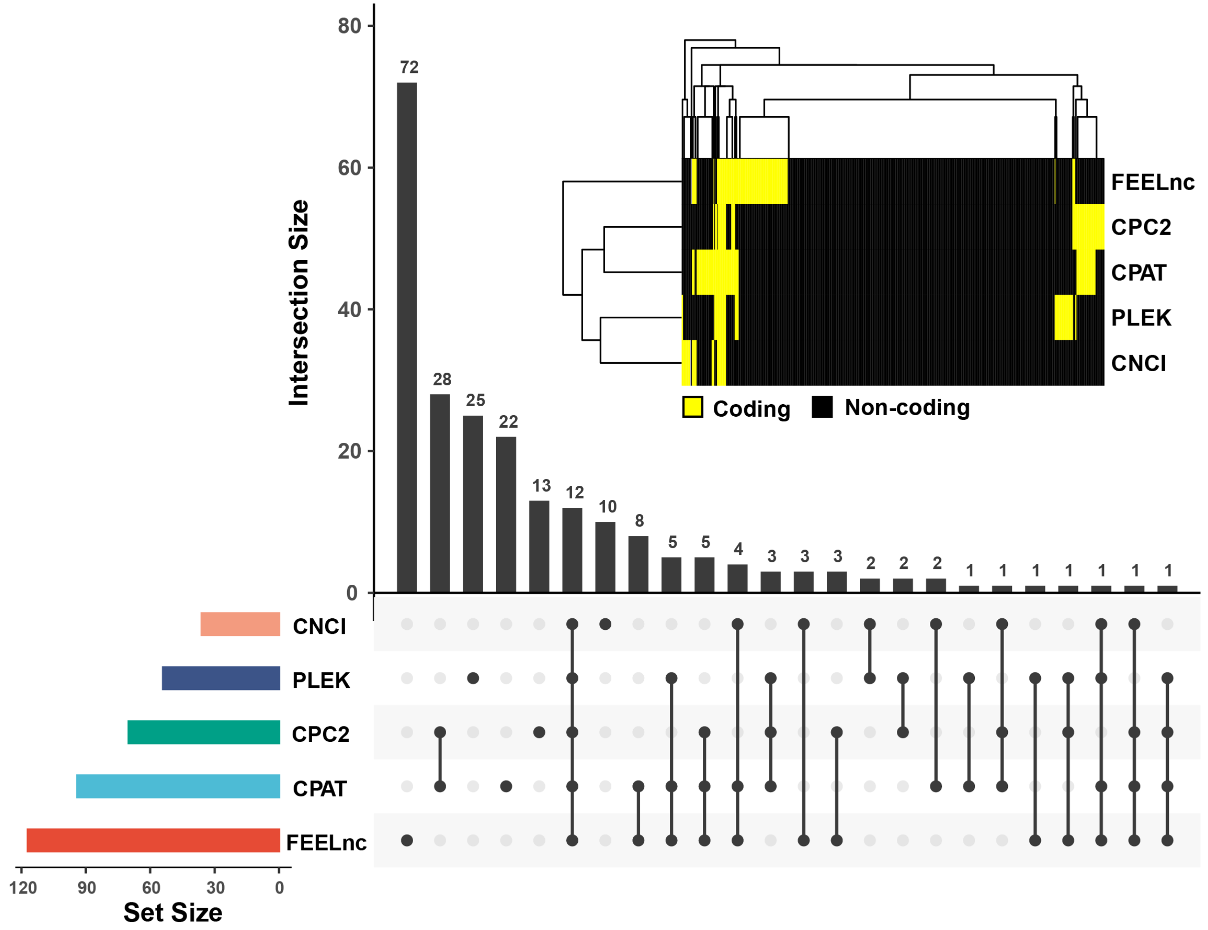

Intersection Size
80
60
40
20
0
72
28
25
22
13
12
10
8
5
5
4
3
3
3
2
2
2
1
1
1
1
1
1
1
FEELnc
CPC2
CPAT
PLEK
CNCI
Coding
Non-coding
CNCI
PLEK
CPC2
CPAT
FEELnc
120
90
60
30
0
Set Size


Fig. S1

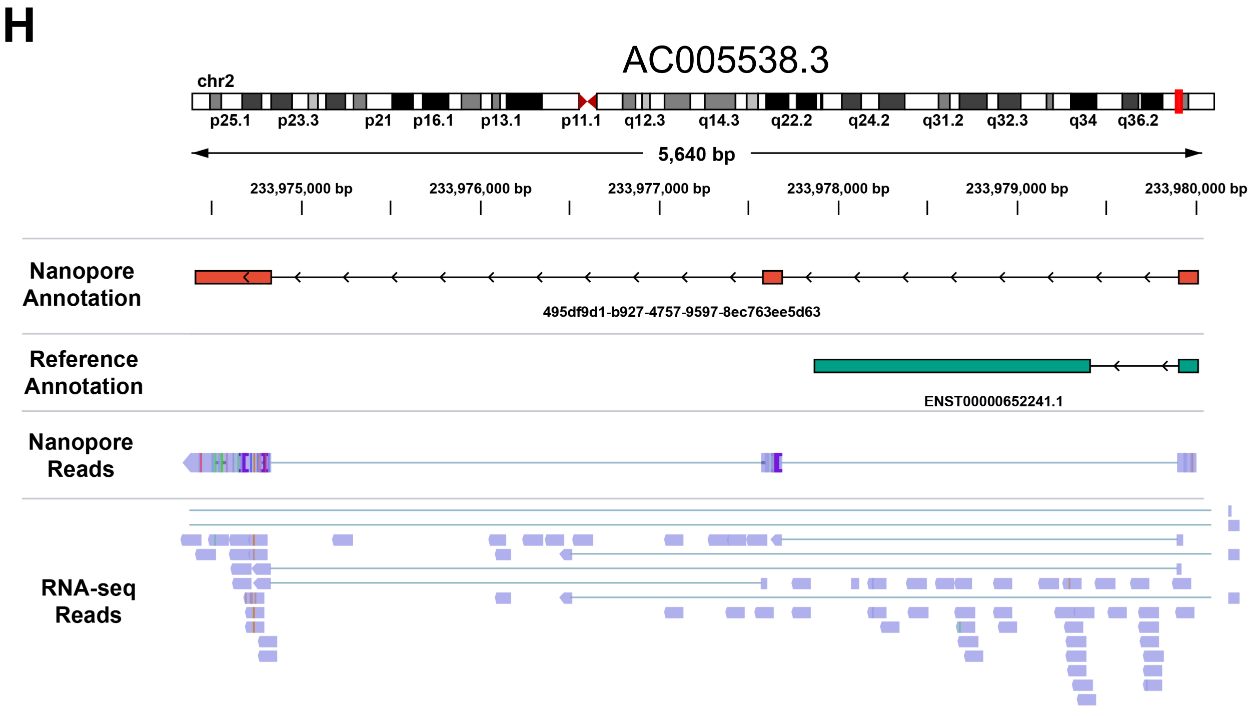


| ID | Label | Coding probability | Peptide length(aa) | Fickett score | Isoelectric point | ORF integrity |
|---|---|---|---|---|---|---|
| 495df9d1 | coding | 0.801057 | 127 | 0.43929 | 9.78997802734 | complete |
| ENST00000652241 | noncoding | 0.154281 | 99 | 0.35741 | 11.1637573242 | complete |

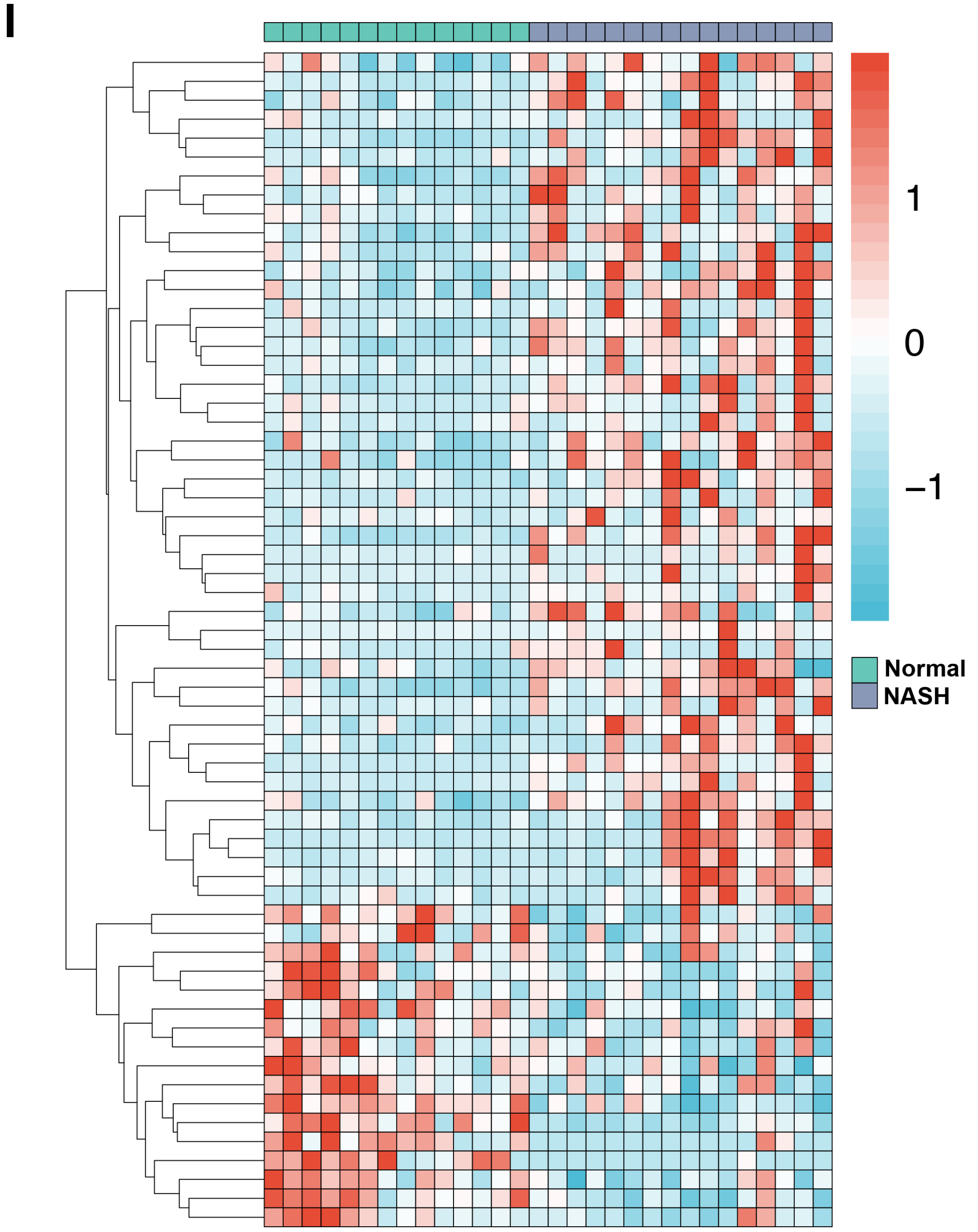


**Fig. S1**

J

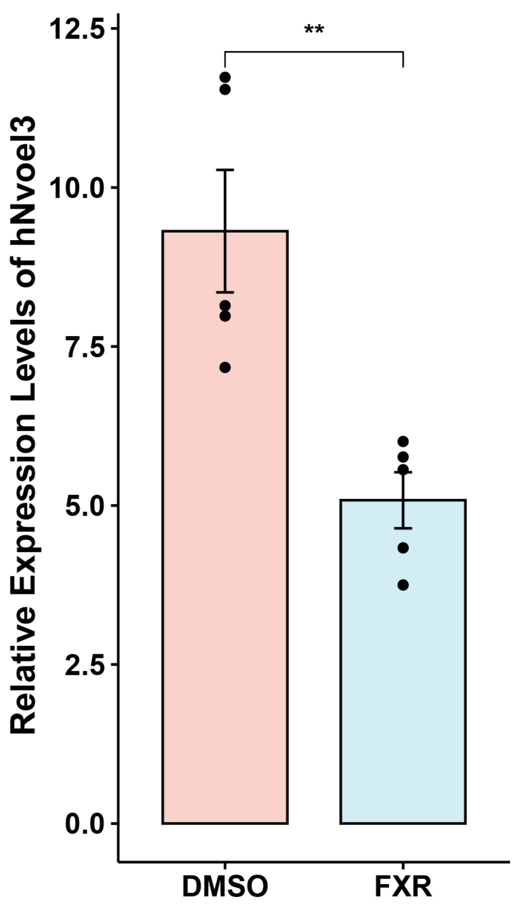

Relative Expression Levels of hNvoel3
12.5
10.0
7.5
5.0
2.5
0.0
**
DMSO
FXR


K

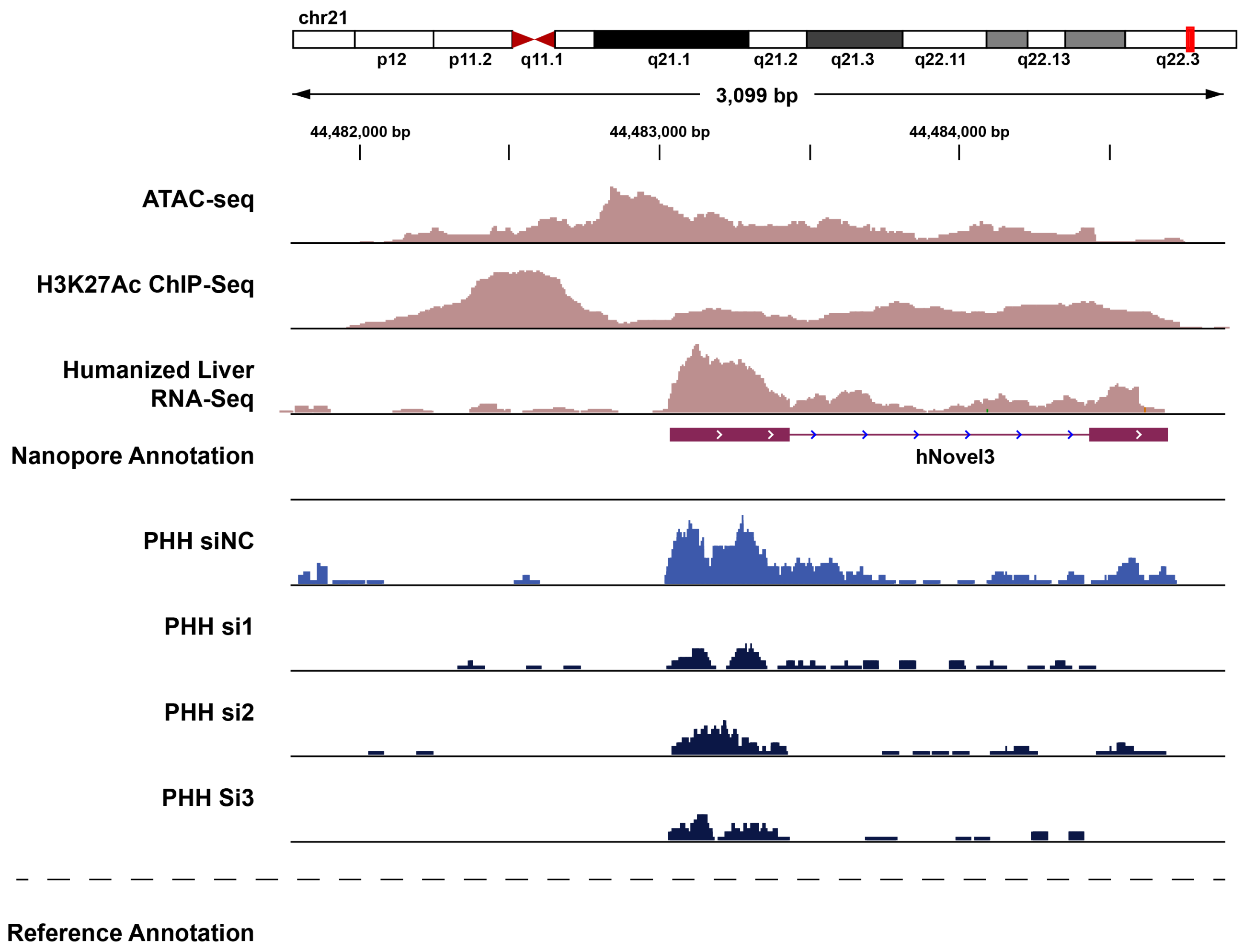

chr21
p12
p11.2
q11.1
q21.1
q21.2
q21.3
q22.11
q22.13
q22.3
3,099 bp
44,482,000 bp
44,483,000 bp
44,484,000 bp
ATAC-seq
H3K27Ac ChIP-Seq
Humanized Liver RNA-Seq
Nanopore Annotation
hNovel3
PHH siNC
PHH si1
PHH si2
PHH Si3
Reference Annotation


Fig. S1

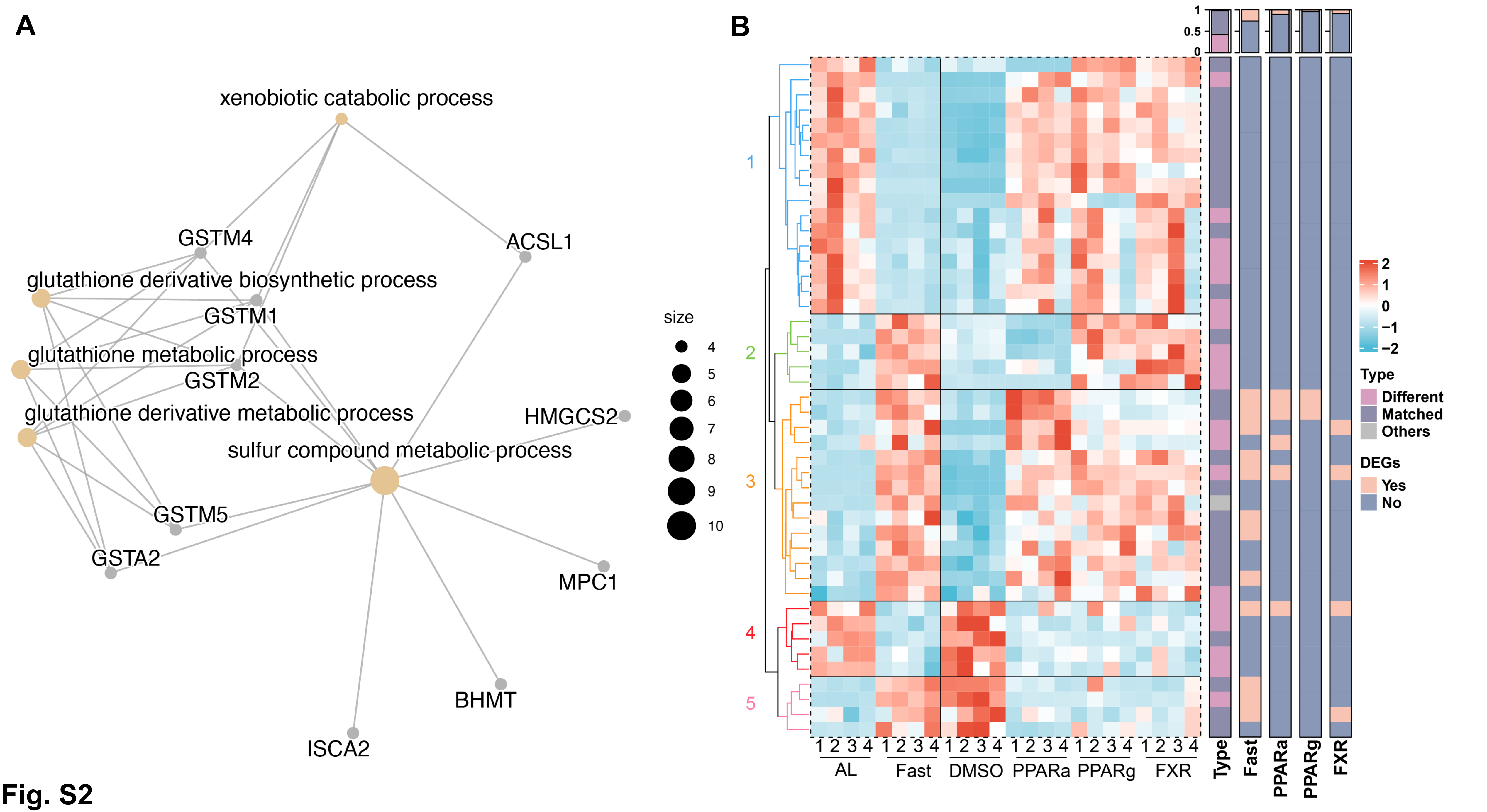
A
xenobiotic catabolic process
GSTM4
ACSL1
glutathione derivative biosynthetic process
GSTM1
glutathione metabolic process
GSTM2
glutathione derivative metabolic process
HMGCS2
sulfur compound metabolic process
GSTM5
GSTA2
MPC1
BHMT
ISCA2
size
4
5
6
7
8
9
10
B
1
2
3
4
5
1 2 3 4
AL
Fast
DMSO
PPARa
PPARg
FXR
Type
Fast
PPARa
PPARg
FXR
2
1
0
−1
−2
Type
Different
Matched
Others
DEGs
Yes
No

Fig. S2

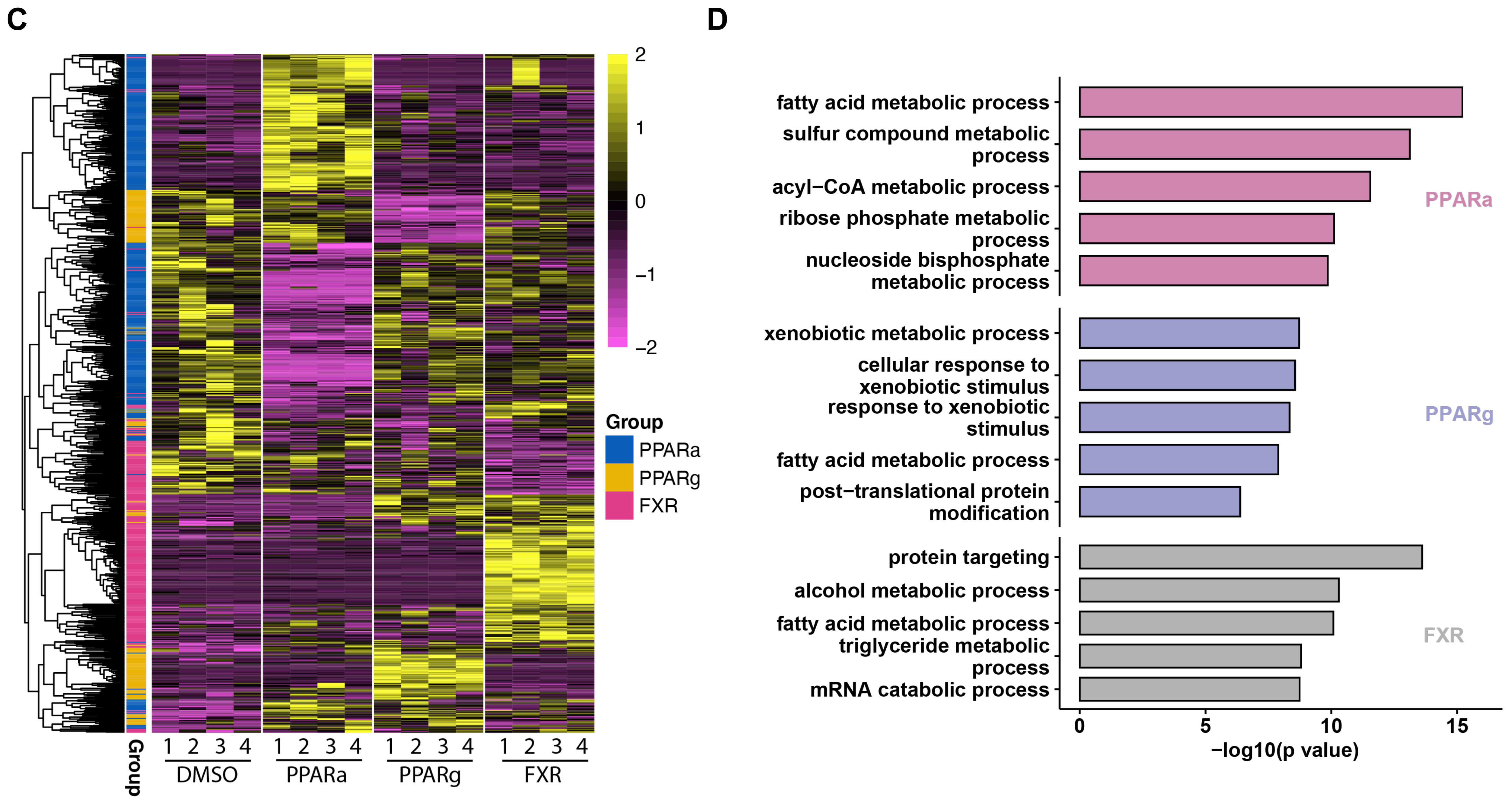
C
2
1
0
−1
−2
Group
PPARa
PPARg
FXR
Group
1 2 3 4
DMSO
1 2 3 4
PPARa
1 2 3 4
PPARg
1 2 3 4
FXR
D
fatty acid metabolic process
sulfur compound metabolic process
acyl−CoA metabolic process
ribose phosphate metabolic process
nucleoside bisphosphate metabolic process
PPARa
xenobiotic metabolic process
cellular response to xenobiotic stimulus
response to xenobiotic stimulus
fatty acid metabolic process
post−translational protein modification
PPARg
protein targeting
alcohol metabolic process
fatty acid metabolic process
triglyceride metabolic process
mRNA catabolic process
FXR
0
5
10
15
−log10(p value)

Fig. S2

E

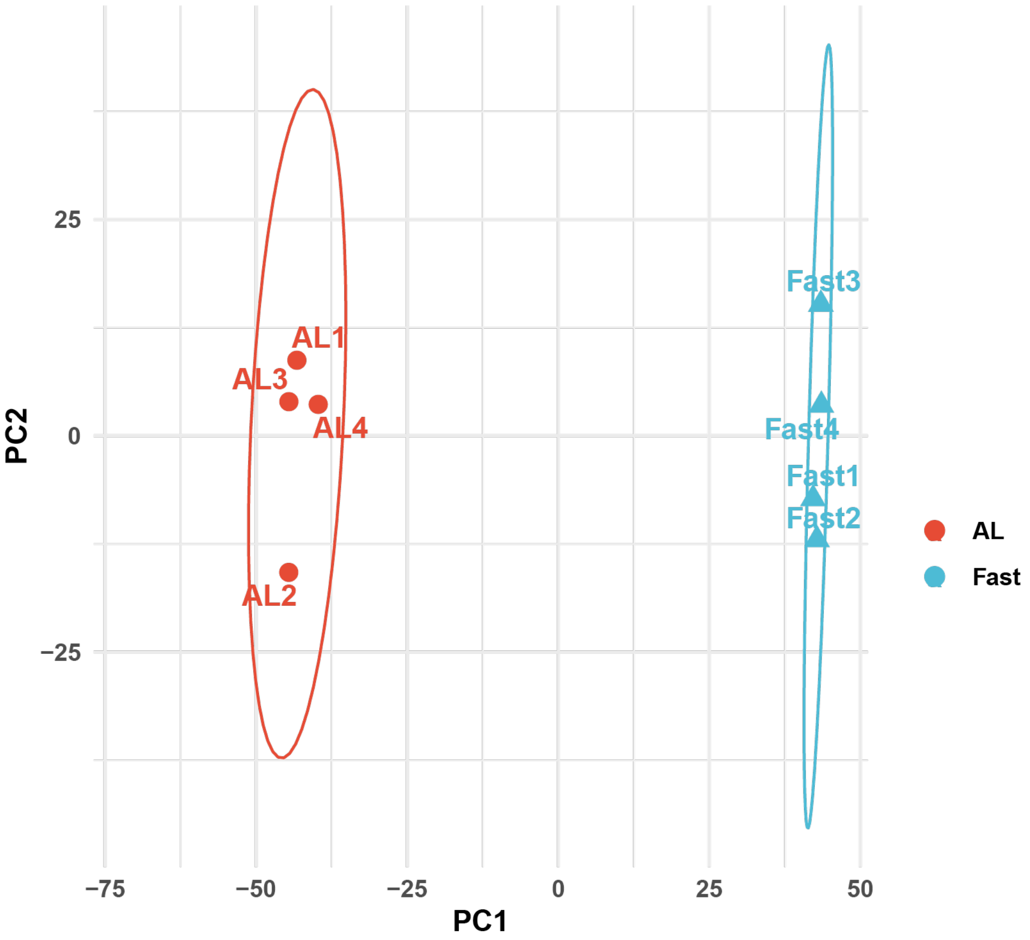
PC2
PC1
25
0
−25
−75
−50
−25
0
25
50
AL1
AL3
AL4
AL2
Fast3
Fast4
Fast1
Fast2
AL
Fast

F

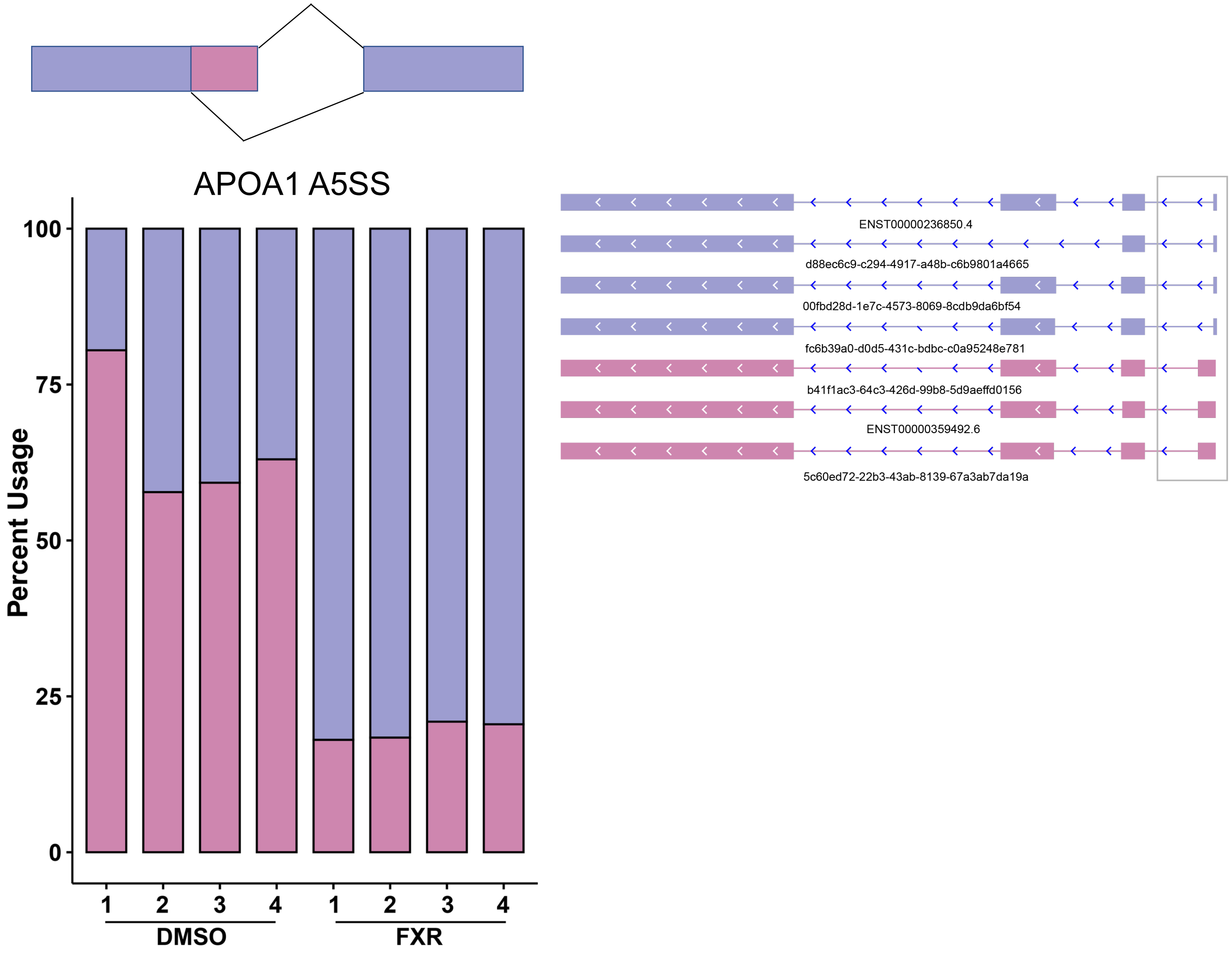
APOA1 A5SS
Percent Usage
100
75
50
25
0
1
2
3
4
1
2
3
4
DMSO
FXR
ENST00000236850.4
d88ec6c9-c294-4917-a48b-c6b9801a4665
00fbd28d-1e7c-4573-8069-8cdb9da6bf54
fc6b39a0-d0d5-431c-bdbc-c0a95248e781
b41f1ac3-64c3-426d-99b8-5d9aeffd0156
ENST00000359492.6
5c60ed72-22b3-43ab-8139-67a3ab7da19a

Fig. S2

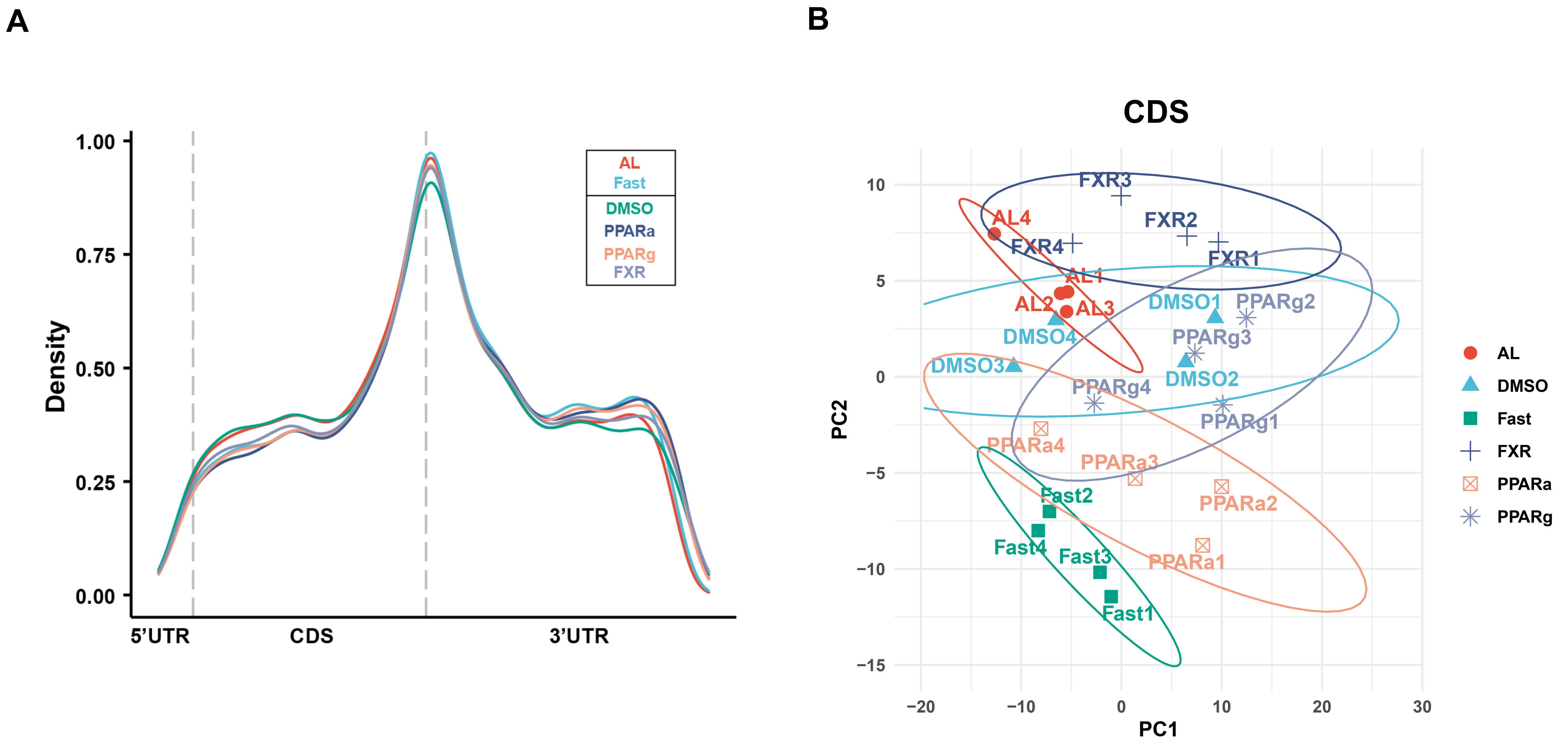
A
Density
1.00
0.75
0.50
0.25
0.00
5'UTR
CDS
3'UTR
AL
Fast
DMSO
PPARa
PPARg
FXR
B
CDS
PC1
PC2
AL
DMSO
Fast
FXR
PPARa
PPARg

Fig. S3

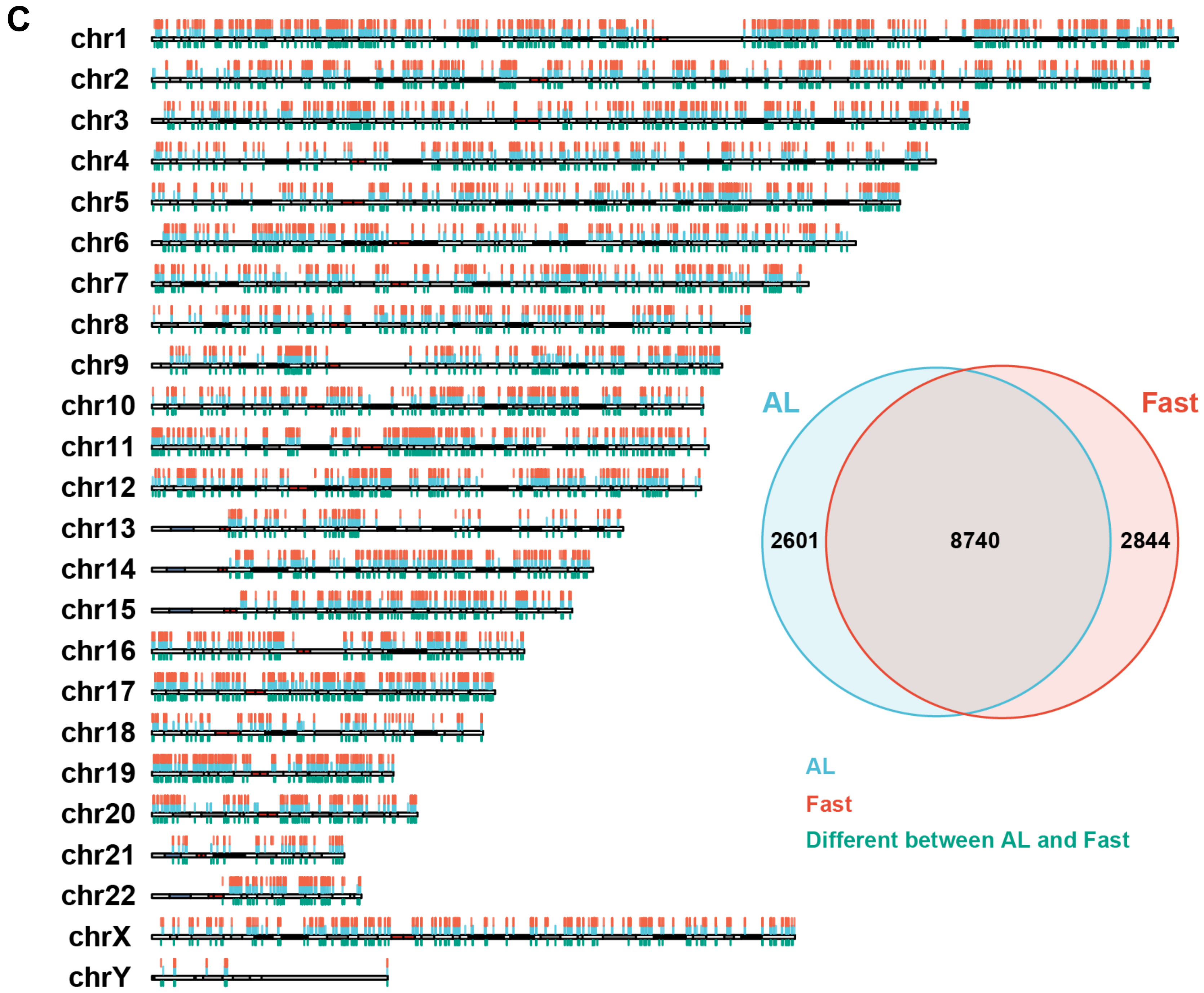
C
chr1
chr2
chr3
chr4
chr5
chr6
chr7
chr8
chr9
chr10
chr11
chr12
chr13
chr14
chr15
chr16
chr17
chr18
chr19
chr20
chr21
chr22
chrX
chrY
AL
Fast
2601
8740
2844
AL
Fast
Different between AL and Fast

Fig. S3

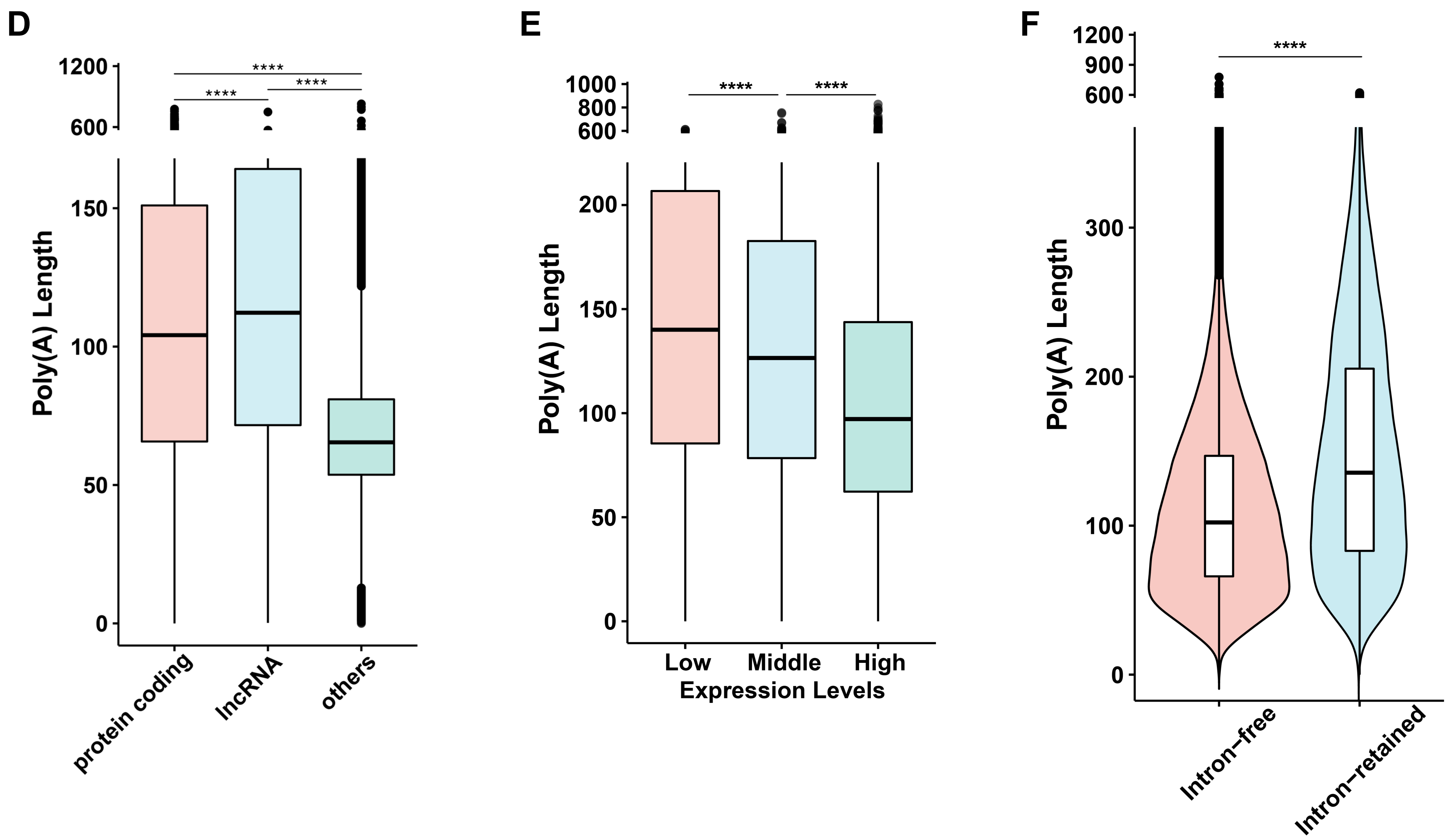
D
1200
600
150
100
50
0
Poly(A) Length
****
****
****
protein coding
lncRNA
others
E
1000
800
600
200
150
100
50
0
Poly(A) Length
****
****
Low
Middle
High
Expression Levels
F
1200
900
600
300
200
100
0
Poly(A) Length
****
Intron−free
Intron−retained

Fig. S3

G

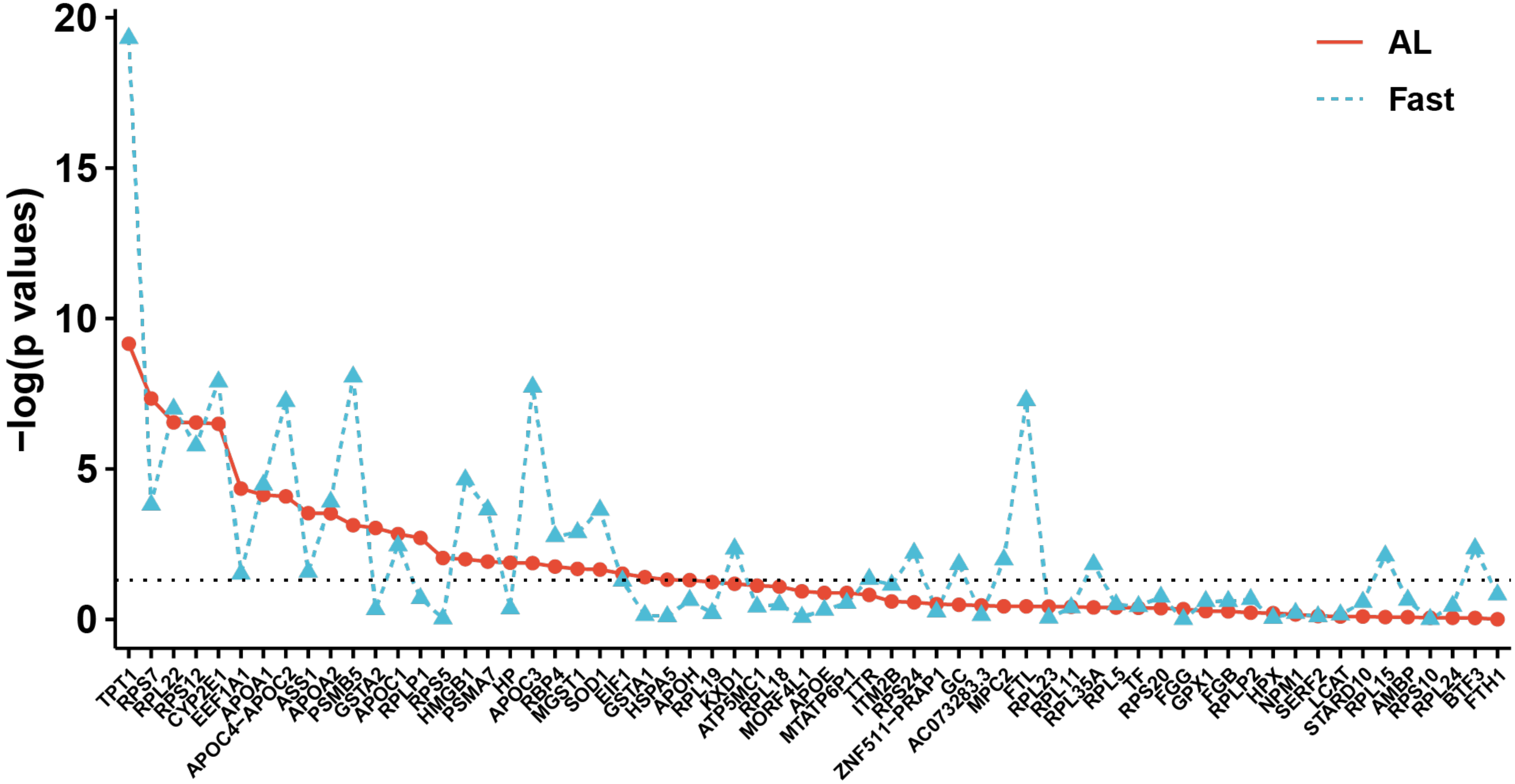

-log(p values)
20
15
10
5
0
AL
Fast
TPT1
RPS7
RPL22
RPS12
CYP2E1
EEF1A1
APOA1
APOC4-APOC2
ASS1
APOA2
PSMB5
GSTA2
APOC1
RPLP1
RPS5
HMGB1
PSMA7
HP
APOC3
RBP4
MGST1
SOD1
EIF1
GSTA1
HSPA5
APOH
RPL19
KXD1
ATP5MC1
RPL18
MORF4L1
APOE
MTATP6P1
TTR
ITM2B
RPS24
ZNF511-PRAP1
GC
AC073283.3
MPC2
FTL
RPL23
RPL11
RPL35A
RPL5
TF
RPS20
FGG
GPX1
FGB
RPLP2
HPX
NPM1
SERF2
LCAT
STARD10
RPL15
AMBP
RPS10
RPL24
BTF3
FTH1


Fig. S3

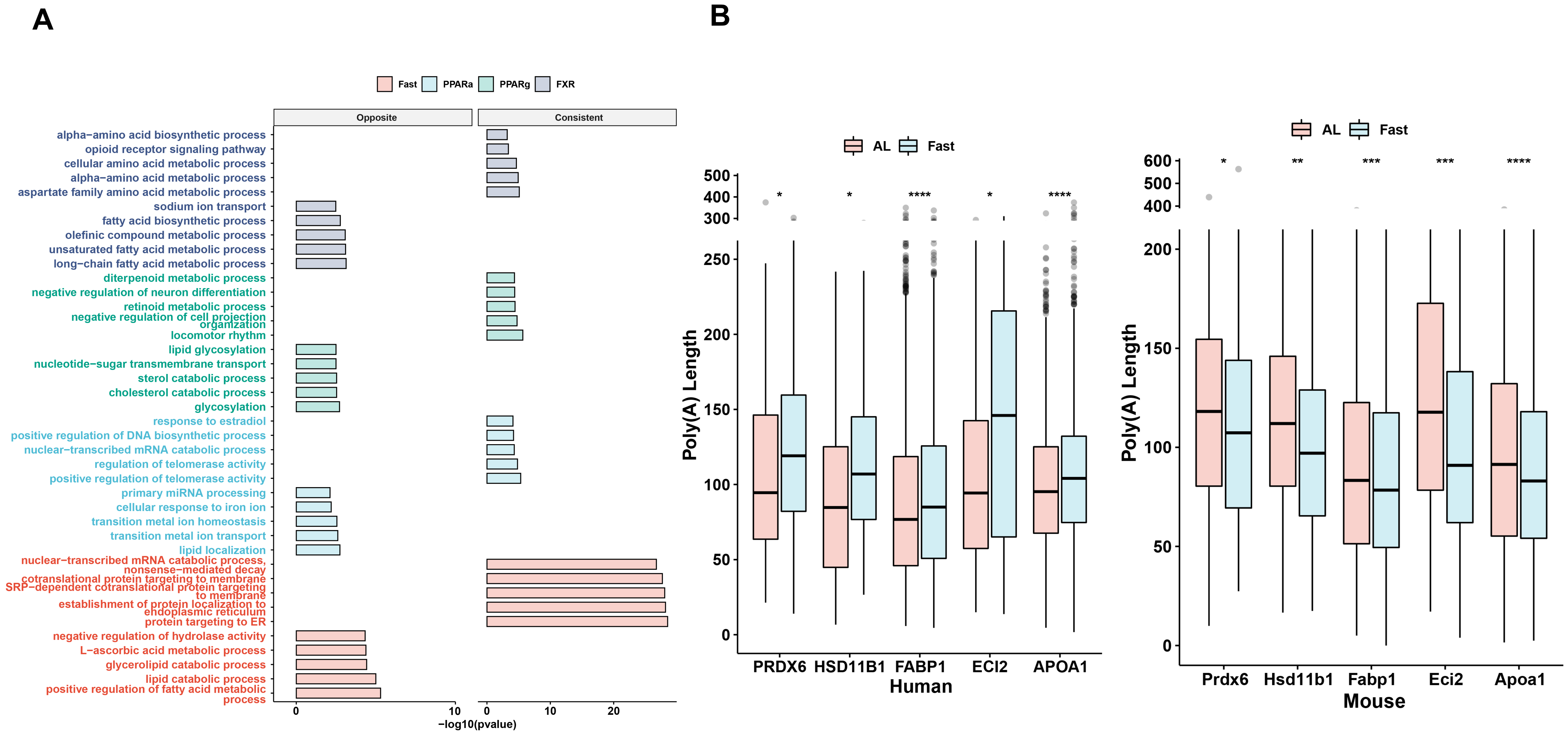
A
Fast
PPARa
PPARg
FXR
Opposite
Consistent
alpha−amino acid biosynthetic process
opioid receptor signaling pathway
cellular amino acid metabolic process
alpha−amino acid metabolic process
aspartate family amino acid metabolic process
sodium ion transport
fatty acid biosynthetic process
olefinic compound metabolic process
unsaturated fatty acid metabolic process
long−chain fatty acid metabolic process
diterpenoid metabolic process
negative regulation of neuron differentiation
retinoid metabolic process
negative regulation of cell projection organization
locomotor rhythm
lipid glycosylation
nucleotide−sugar transmembrane transport
sterol catabolic process
cholesterol catabolic process
glycosylation
response to estradiol
positive regulation of DNA biosynthetic process
nuclear−transcribed mRNA catabolic process
regulation of telomerase activity
positive regulation of telomerase activity
primary miRNA processing
cellular response to iron ion
transition metal ion homeostasis
transition metal ion transport
lipid localization
nuclear−transcribed mRNA catabolic process, nonsense−mediated decay
cotranslational protein targeting to membrane
SRP−dependent cotranslational protein targeting to membrane
establishment of protein localization to endoplasmic reticulum
protein targeting to ER
negative regulation of hydrolase activity
L−ascorbic acid metabolic process
glycerolipid catabolic process
lipid catabolic process
positive regulation of fatty acid metabolic process
0
10
0
10
20
−log10(pvalue)
B
AL
Fast
Poly(A) Length
500
400
300
250
200
150
100
50
0
*
*
****
*
****
PRDX6
HSD11B1
FABP1
ECI2
APOA1
Human
600
500
400
200
150
100
50
0
*
**
***
***
****
Prdx6
Hsd11b1
Fabp1
Eci2
Apoa1
Mouse

Fig. S4

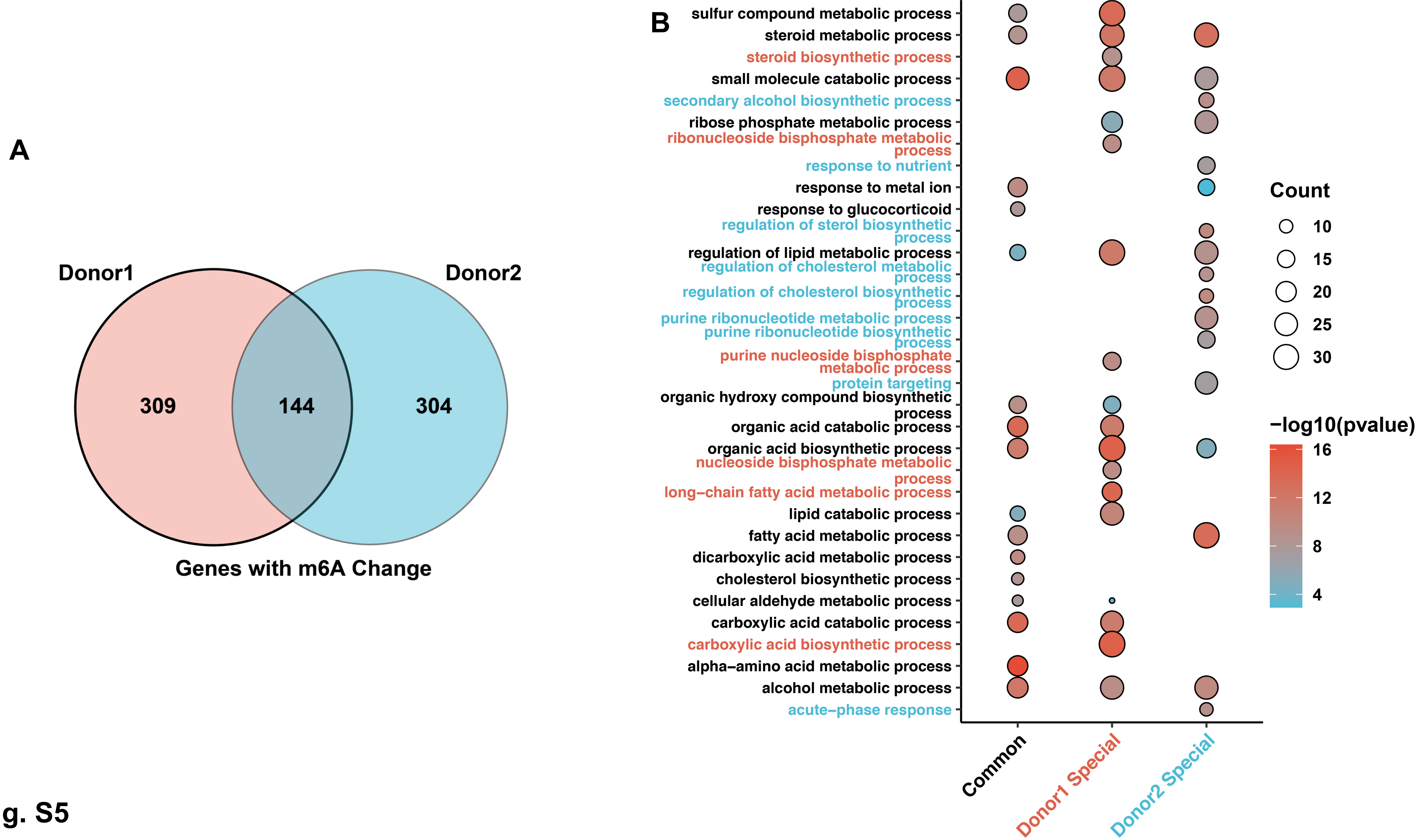
A
Donor1
Donor2
309
144
304
Genes with m6A Change
B
sulfur compound metabolic process
steroid metabolic process
steroid biosynthetic process
small molecule catabolic process
secondary alcohol biosynthetic process
ribose phosphate metabolic process
ribonucleoside bisphosphate metabolic process
response to nutrient
response to metal ion
response to glucocorticoid
regulation of sterol biosynthetic process
regulation of lipid metabolic process
regulation of cholesterol metabolic process
regulation of cholesterol biosynthetic process
purine ribonucleotide metabolic process
purine ribonucleotide biosynthetic process
purine nucleoside bisphosphate metabolic process
protein targeting
organic hydroxy compound biosynthetic process
organic acid catabolic process
organic acid biosynthetic process
nucleoside bisphosphate metabolic process
long−chain fatty acid metabolic process
lipid catabolic process
fatty acid metabolic process
dicarboxylic acid metabolic process
cholesterol biosynthetic process
cellular aldehyde metabolic process
carboxylic acid catabolic process
carboxylic acid biosynthetic process
alpha−amino acid metabolic process
alcohol metabolic process
acute−phase response
Common
Donor1 Special
Donor2 Special
Count
10
15
20
25
30
−log10(pvalue)
16
12
8
4

Fig. S5

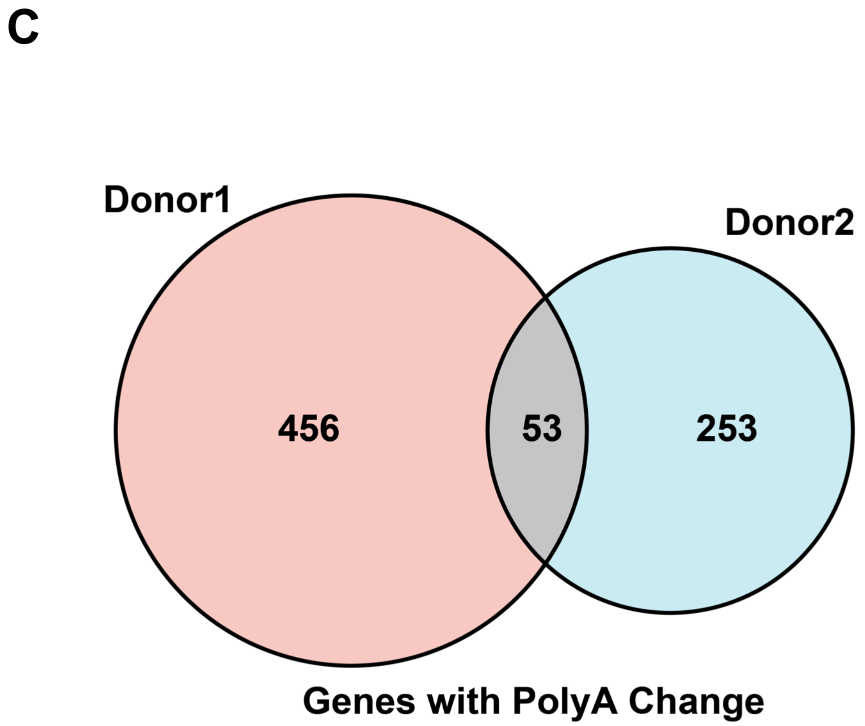
C
Donor1
Donor2
456
53
253
Genes with PolyA Change

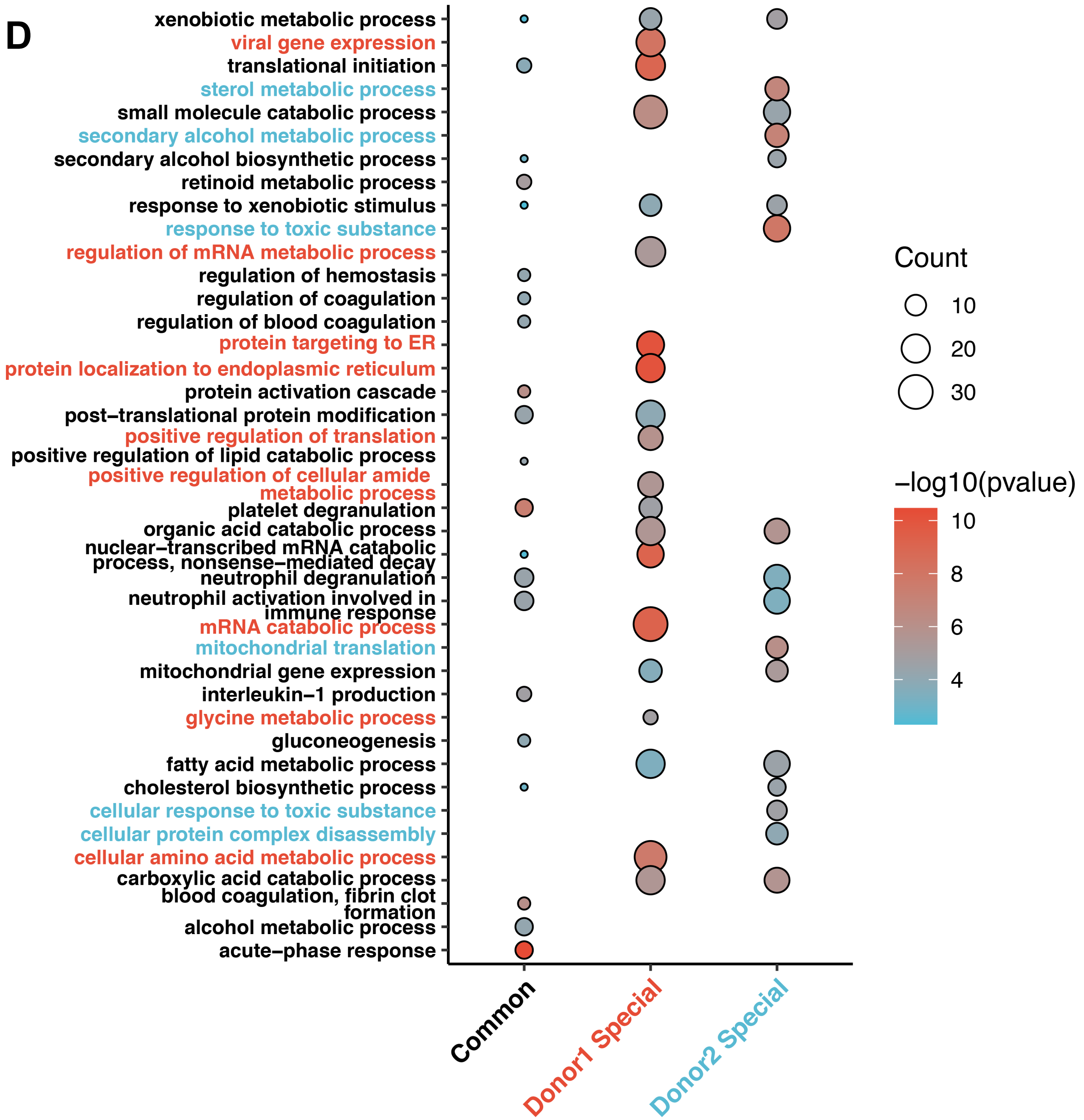
D
xenobiotic metabolic process
viral gene expression
translational initiation
sterol metabolic process
small molecule catabolic process
secondary alcohol metabolic process
secondary alcohol biosynthetic process
retinoid metabolic process
response to xenobiotic stimulus
response to toxic substance
regulation of mRNA metabolic process
regulation of hemostasis
regulation of coagulation
regulation of blood coagulation
protein targeting to ER
protein localization to endoplasmic reticulum
protein activation cascade
post−translational protein modification
positive regulation of translation
positive regulation of lipid catabolic process
positive regulation of cellular amide metabolic process
platelet degranulation
organic acid catabolic process
nuclear−transcribed mRNA catabolic process, nonsense−mediated decay
neutrophil degranulation
neutrophil activation involved in immune response
mRNA catabolic process
mitochondrial translation
mitochondrial gene expression
interleukin−1 production
glycine metabolic process
gluconeogenesis
fatty acid metabolic process
cholesterol biosynthetic process
cellular response to toxic substance
cellular protein complex disassembly
cellular amino acid metabolic process
carboxylic acid catabolic process
blood coagulation, fibrin clot formation
alcohol metabolic process
acute−phase response
Common
Donor1 Special
Donor2 Special
Count
10
20
30
−log10(pvalue)
10
8
6
4

Fig. S5
